# T2T genome of the basal nematode *Xiphinema index* reveals major adaptive events associated with plant parasitism and epigenetic regulation

**DOI:** 10.64898/2025.12.19.695367

**Authors:** Julia Truch, Karine Robbe-Sermesant, Arthur Péré, Dominique Colinet, Corinne Rancurel, Elodie Drula, Martine Da Rocha, Laetitia Perfus-Barbeoch, Ercan Seçkin, Ulysse Julien-Portier, Céline Lopez-Roques, Carole Iampietro, Amalia Sayeh, Marie Gislard, Roxane Boyer, Léo Luvisutto, Daniel Esmenjaud, Etienne GJ Danchin, Cyril Van Ghelder

## Abstract

**Background:** Nematodes constitute a particularly speciose animal phylum, occupying an exceptionally broad range of biomes and encompassing free-living microbe-feeders, predators, and notorious plant or animal parasites. However, the limited availability of genome resources for early-branching nematode clades has constrained our understanding of the molecular determinants underlying these multiple evolutionary adaptations.

**Results:** We present the genome of the plant-parasitic nematode *Xiphinema index*, constituting the first telomere-to-telomere assembly for the Dorylaimida, an early-branching nematode order. Phylogenetic analyses suggest that horizontal gene transfers (HGTs) have played a pivotal role in the emergence of plant parasitism in this phylum. Remarkably, we identify the first documented case of a plant-derived HGT in a nematode genome. This gene encodes a predicted secreted effector of parasitism, underscoring the diversity of evolutionary events that contributed to the parasitic arsenal. Indeed, beyond bacteria and fungi, plants seem to have themselves provided crucial genes for parasitism that may now be used by the nematodes to manipulate them. Our comparative genomic analyses across more than 70 nematode species uncovered an epigenetic toolkit unprecedented in any other invertebrate genome. Indeed, *X. index* harbors a complete set of canonical DNA methyltransferases, a full-length ATRX chromatin remodeler, and a homolog of the 3D-chromatin architectural CTCF protein. Consistent with this repertoire, we detected genome-wide CpG methylation, which inversely correlates with gene expression and chromatin accessibility.

**Conclusions:** Overall, these findings reveal a vertebrate-like epigenetic machinery in an early-branching nematode and call for a fundamental reassessment of the prevailing epigenetic paradigm in nematodes and, more broadly, in animals.

## Background

Nematodes are particularly species-rich [1] and are considered the most abundant animals on Earth [2,3]. Despite their marine origin [4–6], they are also present in terrestrial and freshwater environments as a sign of their adaptive evolutionary success. Their ecology is diverse and encompasses free-living microbe-feeders, predators, and notorious plant or animal parasites. This diversity of trophic habits results from multiple independent evolutionary transitions. Plant parasitism, for example, has independently arisen at least four times during nematode evolution [5,7]. Each transition from a free-living to a plant-parasitic lifestyle has been accompanied by convergent morphological adaptations, most notably the repeated evolution of a needle-like stylet in the mouthpart, which is used to secrete effector proteins and extract nutrients from host plants [8]. At the molecular level, acquisition of plant cell wall-degrading enzymes via horizontal gene transfer (HGT) has been a key evolutionary step towards adaptation to plant parasitism. For instance, cellulase enzymes have been acquired, from bacteria, in major plant parasites such as cyst and root-knot nematodes, while cellulases of fungal origin are present in foliar nematodes [9–11]. Yet, most of our knowledge on the molecular determinants underlying adaptation to plant parasitism remains restricted by the strong phylogenetic bias towards late-branching clades. Notably, no substantial genome resources are currently available for basal clades of plant parasites. Consequently, it remains unclear whether evolutionary events such as HGT are systematically associated with adaptation to plant parasitism in nematodes. Preliminary transcriptomic data in *Xiphinema index* and *Longidorus elongatus*, two plant-parasitic nematodes with basal phylogenetic positions, suggest HGT events took place in early-branching clades as well [12]. However, in the absence of a genome assembly, distinguishing HGT events from contamination in transcriptomic data alone is barely possible.

Beyond trophic adaptation, how the epigenetic mechanisms regulating gene expression and maintaining chromatin integrity have evolved in nematodes remains poorly explored. In most vertebrates, cytosine methylation (5mC) of CpG dinucleotides is a fundamental modification of the epigenetic landscape with a large proportion of CpG dinucleotides being methylated [13,14]. CpG methylation at promoter regions is often associated with gene repression. Deposition and maintenance of 5mC is primarily orchestrated by the interplay of DNA methyl transferase (DNMT) enzymes. Generally, DNMT3 is responsible for *de novo* DNA deposition of CpG, whereas DNMT1 is involved in the maintenance of these epigenetic marks. DNMT2, however, is often associated with RNA methylation [15]. By contrast, invertebrates, which represent the vast majority of animals, display differing DNA methylation patterns and machineries. For instance, nematodes were so far considered to have a null or sparse methylation on cytosine (5mC) across genomes along with a depletion of essential components of the machinery required for the deposition and/or maintenance of the latter one [16–20]. Indeed, the presence of canonical DNMTs is sparse among nematode genomes. However, previous studies showed that methylation of adenine (6mA) could be a fundamental epigenetic mark in *C. elegans* [18], and a modulator of development and parasitism in root-knot nematodes [21,22]. Other key elements of the epigenetic machinery described in mammals include factors influencing three-dimensional organization of the genome such as the insulator CTCF and chromatin remodelers, among which the tumor suppressor ATRX (alpha-thalassemia mental retardation X-linked) [23]. ATRX mutations or knock-down affect important processes including DNA repair and methylation and gene expression. In vertebrates, ATRX is composed of two main domains, a N-terminal ADD domain and a C-terminal helicase/ATPase domain [24]. Its orthologs, named Xnp, in *Drosophila melanogaster* and *C. elegans* are truncated and deprived of ADD domain. Interestingly, besides ATRX, the ADD domain has been only found in DNMT3 proteins. Both are involved in DNA methylation, and the latter two insect and nematode model species show rare or absent DNA methylation on cytosine [25].

Despite recent efforts deployed to produce genomes in basal nematode clades [26], no chromosome-level genome assembly is available for an early-branching plant parasite so far. Therefore, to better understand the evolutionary events associated with adaptation to plant parasitism and the epigenetic regulation of gene expression, we have produced a high-quality telomere-to-telomere (T2T) chromosome-scale genome for the dagger nematode *Xiphinema index*. This is the first representative genome of plant parasites in the basal order Dorylaimida. *X. index* is a virus vector and migratory ectoparasite dreaded by the viticulture sector. This species feeds on actively growing grapevine roots causing malformation, swelling, galling made of multinucleate cells, or necrosis [27,28]. Although high inoculum pressure of *X. index* may cause growth retardation to host plants, the severity of its impact on grapevine, *Vitis vinifera*, is mostly due to its ability to transmit the Grapevine Fanleaf Virus (GFLV) [28]. This nepovirus is the main cause of the lethal grapevine degeneration disease, making it the most severe viral disease of grapevines worldwide [29]. Native to the Middle East, *X. index* was dispersed later along with grapevine’s agricultural history, and is nowadays present in most if not all vineyards worldwide [30]. Consequently, *X. index* was the only virus vector to be included in the top 10 plant-parasitic nematodes (PPNs) in terms of agronomical and scientific importance [31]. While both GFLV variants and *Vitis* species and cultivars now have a significant number of omics resources available, these crucial assets were lacking for *X. index* impeding the in-depth study of this three-partner pathosystem.

Our comparative analysis of the *X. index* genome with those of 70 other nematode species, combined with large-scale phylogenetic analyses unveiled two major findings. First, we revealed that plant cell wall-degrading enzymes and other proteins were also acquired by HGT in *X. index*, suggesting these evolutionary events played major roles in the multiple adaptations to plant parasitism across the nematode tree of life. Remarkably, in addition to HGT of bacterial and fungal origin, we also report the first case of HGT of plant origin in a nematode. Second, our comparative genomics analysis revealed in *X. index,* an epigenetic toolkit of unparalleled completeness in nematodes, and potentially one of the most comprehensive reported across invertebrates. We unveiled (i) a full and large arsenal of DNMTs, (ii) a global genome methylation at CpGs and (iii) an inverse relationship between 5mC levels, chromatin accessibility around TSS, and gene expression. Furthermore, we unexpectedly identified a CTCF and a full-length ATRX ortholog. These findings suggest that, in certain respects, the epigenetic arsenal of *X. index* resembles that of vertebrates [32] and impose a rethinking of the evolution of the epigenetic toolkit across the animal tree of life.

## Results

### A first T2T reference genome for the early-branching nematode order Dorylaimida

To generate the *X. index* genome, tens of thousands of individuals (**Figure 1A**) were collected from soil to produce three independent DNA samples. We combined Oxford Nanopore Technologies, PacBio HiFi and Hi-C sequencing approaches, to generate a final assembly of 73 contigs with a N50 length of 23.66 Mb and a L90 of 10, for a total size of 225.83 Mb, which is consistent with the k-mer estimation (ca. 219 Mb haploid) **(Figure S1, Table S1**). Plots of k-mers occurrence and multiplicity in the HiFi reads and in the genome assembly confirmed that the genome is diploid with a relatively high heterozygosity of ∼0.9%, and that the diversity of sequence data present in the raw reads was well incorporated in the assembly **(Figure S1)**. The Hi-C contact map yielded 10 large pseudo-molecules **(Figure 1B),** and the canonical nematode telomeric repeat sequence (TTAGGC)n was present at both ends of each pseduso-molecule **(Figure 1C)**. These results suggest a T2T assembly of 10 chromosomes, n=10, which is consistent with previous cytogenetic observations [33]. In addition to the canonical nematode telomeric DNA repeat, we identified a candidate telomerase (xiphind_chr_06g013610) (**Figure S2**), thus confirming a general trend of co-occurrence of the repeat and the enzyme observed in nematodes. However, other proteins associated with telomeres composing the shelterin complex showed more variations in terms of conservation across nematodes **(Table S2, Figure S2**), consistent with previous observations [34]. A total of 30,614 protein-coding genes were predicted (**Table S3**), and 54.6% of the predicted proteins contained at least one InterPro domain (https://doi.org/10.57745/B4WGC9). Among the proteins without functional annotation, 10,540 had no predicted ortholog in 70 other nematode proteomes used for orthofinder analysis and might be specific to *Xiphinema* (https://doi.org/10.57745/6CUXWM). Proteins with neither functional annotation nor recognizable homology in other nematodes constitute an interesting source of novelty to be further characterized. The secretome of *X. index* was constituted by 2,959 proteins with a predicted signal peptide for secretion but neither transmembrane region nor GPI anchor, hereafter referred to as predicted secreted proteins (PSPs) (https://doi.org/10.57745/TCN7RX). The mitochondrial genome of *X. index* was larger (14,820 bp) than those publicly available for other *Xiphinema* species, which was mostly due to the presence of two duplicated non-coding regions (NCR), showing no similarity with any other sequences (**Figure S3**). Concurrently, we identified several nuclear mitochondrial DNA segments (NUMTs; >500bp), including the largest one (11.4 kb), sharing 96.24% identity with the mitogenome **(Figure 1C)**. This insertion was further confirmed by PCR in two unrelated populations of *X. index* originally from Iran (*X. index “Iran”*) and Samos (genome), indicating an ancient insertion in the nuclear genome (**Figure S3**). However, the functional implications of the presence of untranscribed mitochondrial-related sequences in the genome have yet to be uncovered.

**Figure 1:**
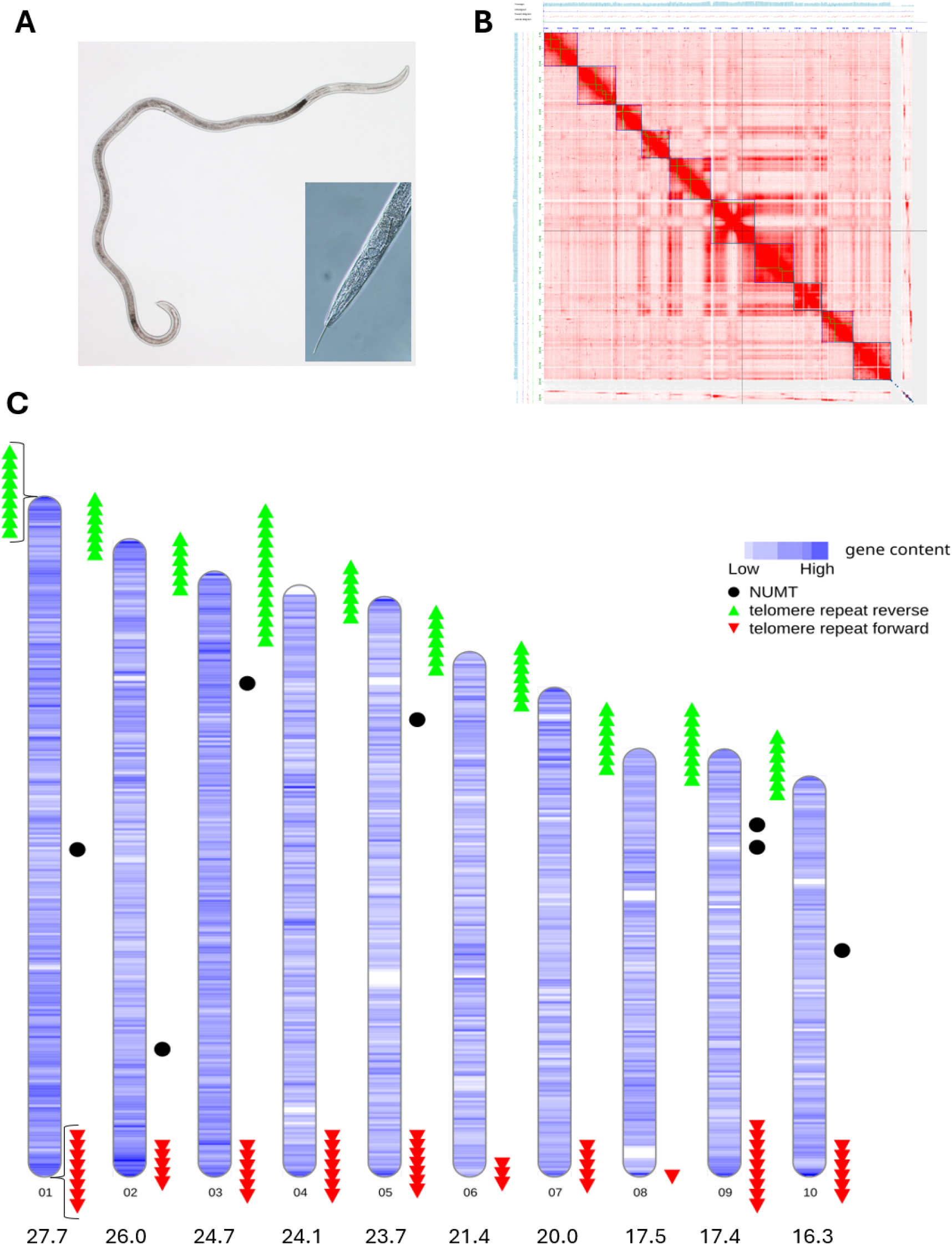
Genome assembly of *Xiphinema index.* (A) *X. index* adult female 3.5mm long with a close-up of its large stylet, a classic apparatus of plant parasitic nematodes. (B) Genome-wide Hi-C contact matrix of *X. index*. (C) Representation of the 10 chromosomes of *X. index* showing the gene density, mitochondrial DNA insertions (NUMT) and positions of oriented telomeric repeats. The number of triangles reflects the size of repeat arrays of the canonical nematode telomeric sequence (TTAGGC)n. The lengths in Mb of the 10 chromosomes are indicated below.

### Horizontal gene transfers have contributed to the nematode parasitic arsenal

In the context of plant-parasite interactions, Carbohydrate-Active enZymes (CAZymes) play an essential role in degrading the plant cell wall and digesting storage sugars. We therefore delineated their repertoire (CAZome) in *X. index*. Using the CAZy database [35], we identified 392 CAZymes made of 473 modules and compared this CAZome to those of five other nematode species from the same Dorylaimia subclass. The number of CAZymes was lower in the four Trichinellida animal parasitic species than in the two Dorylaimida species (*X. index and Mesodorylaimus*) **(Table 1)**. Clustering the species according to the abundance of each CAZyme family module involved in the degradation of carbohydrates recapitulates the phylogeny of the species, with the Trichinellida on one side and the Dorylaimida on the other (**Figure S4**). Beyond Dorylaimia, seven CAZymes were found to be *X. index-*specific, when compared to 70 other highly contiguous nematode genomes using Orthofinder (https://doi.org/10.57745/6CUXWM). Among these specific CAZymes, five were included in the PSPs, highlighting them as candidate effectors to be further functionally assessed. These five CAZymes belonged to the three glycoside hydrolase (GH) enzyme families in *X. index* absent from the other Dorylaimia tested, for which activities are known or predicted: GH12 (cellulose degradation, previously characterized in *X. index* [12]), GH28 (pectin degradation), and GH32 (sucrose degradation). As all these activities are related to degradation or digestion of plant carbohydrates, we then classified Dorylaimia species according to their abundance of CAZymes in which at least one of the possible activities includes degradation of plant sugars [36] (**Figure S5**). In this classification, *X. index* stands out from the rest with a clear outgroup position, reflecting its rich and idiosyncratic enzymatic arsenal for plant sugar degradation.

**Table 1:** Annotation of CAZyme modules in the predicted proteomes of nematodes in the Dorylamia group. Modules Abbreviations GH: Glycoside Hydrolase, GT: Glycosyl Transferase, PL: Polysaccharide Lyase, CE: Carbohydrate Esterase, CBM: Carbohydrate Binding Module, AA: Auxiliary Activity. Note that a protein can bear multiple CAZyme modules.

| Species / CAZyme modules | GH | GT | PL | CE | CBM | AA | Total modules |
| --- | --- | --- | --- | --- | --- | --- | --- |
| *Xiphinema index* | 115 | 284 | 0 | 1 | 59 | 14 | 473 |
| *Mesodorylaimus YZB2_4* | 132 | 250 | 0 | 2 | 74 | 14 | 472 |
| *Trichinella nelsoni* | 88 | 139 | 0 | 4 | 45 | 1 | 277 |
| *Trichinella spiralis* | 43 | 88 | 0 | 1 | 22 | 1 | 155 |
| *Trichuris muris* | 57 | 124 | 0 | 3 | 59 | 3 | 246 |
| *Trichuris trichiura* | 61 | 124 | 0 | 2 | 55 | 1 | 243 |

Previous analyses have shown that a GH12 cellulase was likely acquired via HGT in *X. index* [12]. Furthermore, cellulases of families GH5 and GH45, as well as GH28 polygalacturonases, GH32 invertases, other CAZymes and other enzymes have been reported as acquired via HGT in distantly related plant-parasitic nematodes such as root-knot nematodes, cyst nematodes or foliar nematodes [9,11,37]. Therefore, to investigate whether HGTs have also substantially contributed to the CAZome and the parasitic arsenal of *X. index*, we performed a comprehensive and accurate phylogenetic detection of HGT candidates of non-animal origin. Among the 30,614 *X. index* predicted proteins, 1,449 had an Alien Index (AI) or Aggregate Hit Support (AHS) greater than 0, indicating that they are more similar to non-animal than to animal proteins (**Table 2**). Of these, automated phylogenetic analyses and topology tests identified 224 as possible Longidoridae-specific HGT candidates. After applying stringent phylogenetic criteria (see Methods section), 44 HGT candidates were considered highly likely and validated (**Table 2**).

**Table 2.** Screening of HGT candidates in the predicted proteome of *Xiphinema index*.

|  | *Xiphinema index* |
| --- | --- |
| Number of protein-coding genes | 30,614 |
| Number of predicted proteins with AI or AHS > 0 | 1,449 |
| Number of HGT candidates detected by AvP | 224 |
| - Bacterial origin | 93 |
| - Fungal origin | 38 |
| - Plant origin | 30 |
| - Other origin | 63 |
| Number of validated HGT candidates (and transfer events) | 44 (29) |
| - Bacterial origin | 36 (23) |
| - Fungal origin | 7 (5) |
| - Plant origin | 1 (1) |

These HGTs are widely distributed across the genome, with chromosomes 1 and 4 harbouring the highest number of HGTs whereas chromosomes 7 and 8 contained the fewest, proportionally (**Figure 2A**). In addition, 21 out of 44 HGTs encoded PSPs, making them noteworthy effectors to be further studied (i.e., PSPs are significantly enriched in the HGT subset compared to the whole proteome; Fisher’s exact test, p = 8.5 × 10⁻¹¹). These 44 validated HGTs originated from at least 29 independent transfer events, mainly from putative bacterial donors, while a smaller proportion was presumably acquired from fungal donors, and, notably, one case was attributed to a plant donor (xiphind_chr_06g025050) (**Table 2** and **Table S4**). Consistent with a plant origin, no Metazoa sequences were found in the Diamond homology search with the default maximum of 500 hits, and thus none are present in the phylogenetic tree (**Figure 2B**). Five Metazoa sequences, including two Nematoda sequences, were found in the phylogenetic tree when the Diamond analysis was extended to a maximum of 5,000 hits (**Figure 2C**). However, the five Metazoa sequences did not group with the *X. index* sequence. An alternative topology, constrained to support the monophyly of all Metazoa sequences, was significantly less likely than the unconstrained topology, supporting an HGT of plant origin for the *X. index* sequence **(Figure 2C)**. Additionally, we identified six CAZymes likely acquired via HGT in four independent events, all originating from bacteria (**Figure 2D-G** and **Table S4**). These CAZymes belonged to the three families with known activities on plant sugars described above: GH28, GH32, and GH12, as well as a GH31 of unknown function **(Figure 2D-G)**. We confirmed the bacterial origin of the GH12 and GH32 previously reported in *X. index* **(Figure 2E-F)**. However, no GH28 **(Figure 2D)** was identified in the previous transcriptome analysis of *X. index* and *Longidorus elongatus* [12]. Overall, HGT events have substantially contributed to the parasitic arsenal of *X. index*, and our genome annotation improved HGT detection as only 12 out of the 44 HGTs were identified in the previous transcriptomic analysis (**Table S4)**.

**Figure 2:**
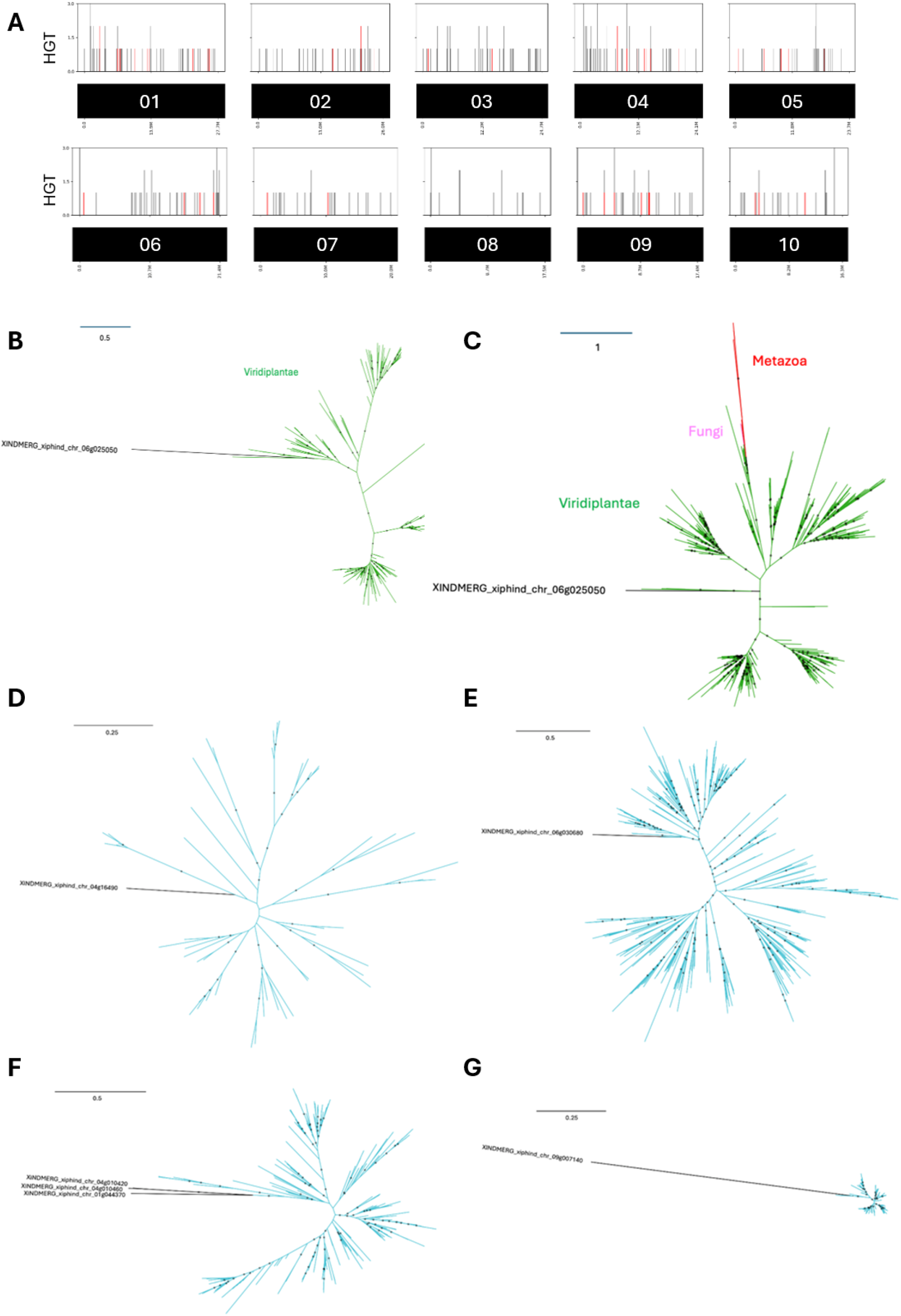
Genome-wide distribution of HGTs and phylogenetic trees of the plant-origin HGT and of CAZyme HGTs. (A) Distribution of the 224 AvP-predicted HGTs (black lines) and the 44 validated HGTs (red lines) across the 10 chromosomes. The number of HGTs range from 0 to 3 per 100 kb window. (B-G) The *X. index* sequences are shown in black. Green branches represent plant sequences. Other branch colors are as follows: red = Metazoa sequences, pink = fungal sequences, blue = bacterial sequences. Gray circles indicate nodes with support values greater than or equal to 80% and 90% for SH-aLRT and UFboot, respectively. (B) Phylogenetic tree from Diamond analysis with a maximum of 500 hits. (C) Phylogenetic tree from Diamond analysis with a maximum of 5,000 hits and the use of a gap threshold of 0.5 for trimAl instead of the -automated1 option. (D) XindB05 HGT event corresponding to GH28. (E) XindB07 HGT event corresponding to GH32. (F) XindB12 HGT event corresponding to GH12. (G) XindB17 HGT event corresponding to GH31. For XindB17, the CD-HIT analysis was performed with an identity threshold of 98%, as all bacterial sequences from the Diamond analysis were very similar. While bacterial GH31 sequences are highly conserved, the divergence in the *X. index* sequence explains the length of the corresponding branch in the tree.

To our knowledge, our study presents the first documented case of HGT of plant origin in a nematode [38]. Although the exact plant donor species is unknown, the closest sequences are from Dicotyledons. RNA-seq reads analysis indicates that *xiphind_chr_06g025050* is expressed **(Figure S6).** Furthermore, we successfully amplified its genomic insertion in an independent *X. index* population, using a genomic marker (named HGT1) designed in the first intron of this gene, thus ruling out a hypothetical contamination **(Figure S6)**. This atypical HGT encodes a putative calcium binding protein (calmodulin), while various enzymes represent the large majority of the rest of the HGTs. The functional annotation of xiphind_chr_06g025050 revealed a signal peptide for secretion followed by a consensus disordered predicted region and the canonical 4 EF-Hand domains found in calmodulin proteins that are known to bind calcium ions **(Figure S6)**. The alignment between this candidate and its 10 closest sequences in databases showed that (i) it is the only one to harbor a signal peptide, (ii) it displayed the conserved binding pockets including amino acids involved in calcium binding. Future studies dedicated to this putative effector are required to fully functionally assess the mechanism behind it.

### The epigenetic toolkit of *X. index* is unprecedented in nematodes

As *X. index* represents the most basal plant parasitic nematode with a T2T genome currently available [26], it represents a key resource for investigating the evolution of fundamental biological processes and putting into perspective the knowledge acquired with the extensively studied model nematode *C. elegans*. We therefore characterized the *X. index* epigenetic toolkit, focusing on components that have been reported as lost during nematode diversification and may retain signatures of ancestral regulatory mechanisms. We first mined the predicted proteome and genome assembly of *X. index* using the human (*Homo sapiens*) DNMTs as queries (hDNMT1: P26358; hDNMT2: O14717; hDNMT3: Q9UBC3) to identify canonical DNMT candidates in nematodes. Notably, we identified three DNMT1 homologs in *X. index*: xiphind_chr_02g008810 (dnmt1a), xiphind_chr_03g023520 (dnmt1b) and xiphind_chr_01g045810 (dnmt1c). These findings were unexpected, as DNMT1 genes have only been reported in a few nematode species to date, and in unrelated clades [16–20]. To untangle this apparent discrepancy, we reanalysed these previously reported nematode DNMT1 candidates in the light of the three *X. index* homologs. Protein domain analysis revealed that the previously reported nematode DNMT1 candidates all lacked important characteristic domains of canonical DNMT1 proteins, including the foci-targeting domain (DNMT1-RFD) (**Figure S7)**. In addition, half of these also lacked the C-terminal BAH and DNA methylase domains. In contrast, dnmt1a, dnmt1b and dnmt1c from *X. index* contained both the RFD domain and the canonical C-terminal domains of DNMT1 (**Figure 3A and 3B** and **Figure S7)**. Besides, in *X. index*, we identified one DNMT2 (xiphind_chr_02g036660) and one DNMT3 (xiphind_chr_02g037650) candidate proteins, thus completing the whole DNMT arsenal. In the light of these surprising findings, we mined *X. index* genome for other major epigenetic players, such as the chromatin remodeler ATRX. To date, nematodes are generally considered to possess only a truncated version of ATRX [39], referred to as XNP-1 in *C. elegans* [40]. XNP-1 lacks the N-terminal ATRX-DNMT3-DNMT3L (ADD/PHD) domain, which is also a characteristic domain of DNMT3 proteins. In addition to the xnp-1 ortholog (xiphind_chr_02g004130), the genome of *X. index* contained one candidate gene coding for a full ATRX protein (xiphind_chr_03g035120) comprising the ADD/PHD N-terminal domain (**Figure 3A and 3C and Figure S8)**. To our knowledge, *X. index* is the first organism reported to have a full version of the chromatin remodeler ATRX in addition to the xnp-1 ortholog. As for *X. index* RFD in DNMT1 candidates, domain analysis and 3D structure prediction of *X. index* ADD domain in ATRX candidate showed a strong similarity with those of Human (**Figure 3B-C, Figure S7, S8)**. Using the PacBio HiFi long reads, we double checked that the contiguous presence of the characteristic domains of these key epigenetic players was supported in the genome assembly (**Figure S9A**). Furthermore, analysis of RNA-seq data showed that the dnmt1a, dnmt1b, dnmt1c, dnmt2, dnmt3 and *ATRX* gene predictions were all well supported by transcriptomic data and were substantially expressed (**Figure S9B**).

**Figure 3:**
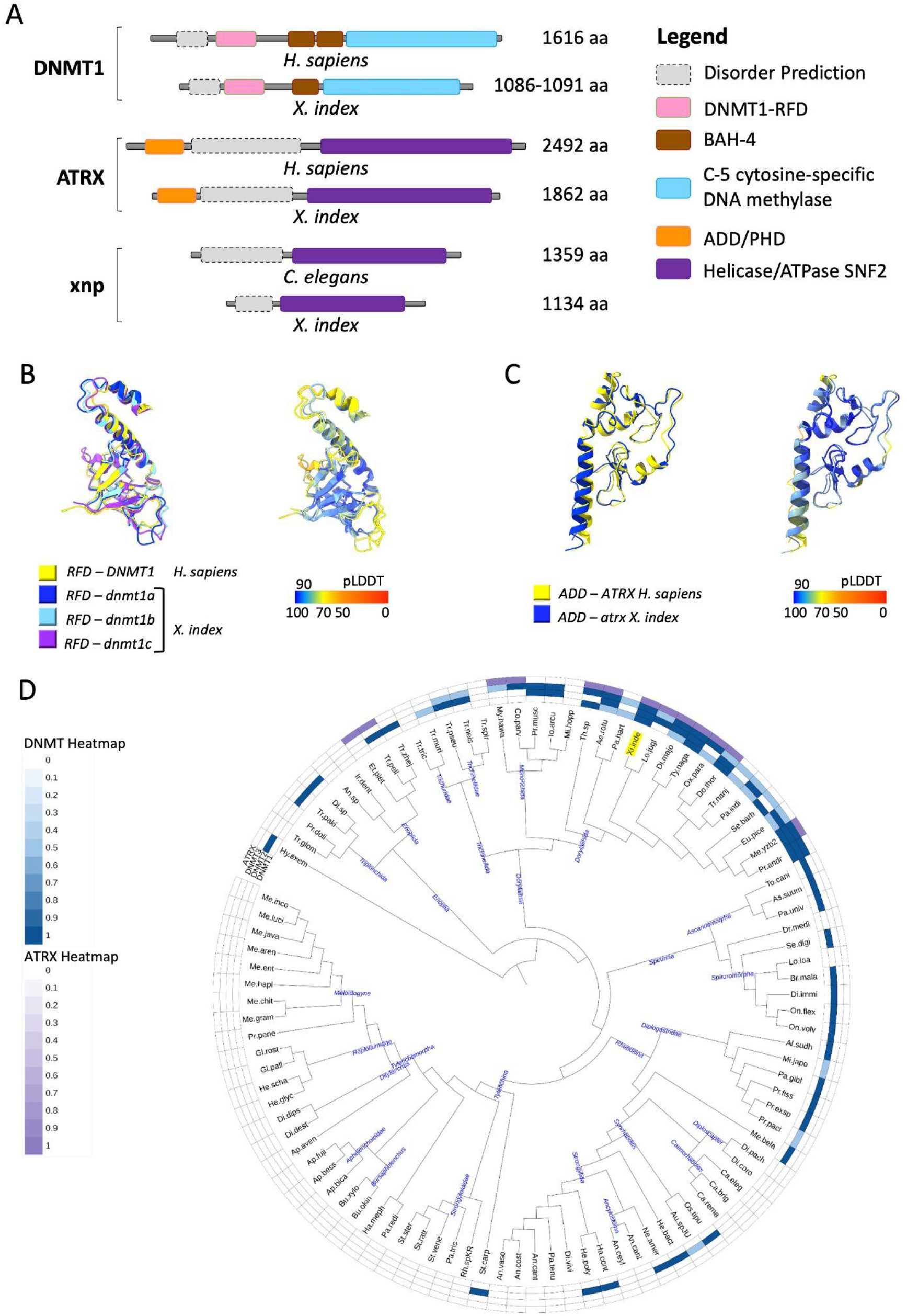
**A.** Domain structure of the canonical ATRX and DNMT1 proteins. The illustration highlights characteristic crucial domains that define these proteins (i.e. RFD and C-5 DNA methylase, and ADD/PHD and Helicase/ATPase SNF2 domains for DNMT1 and ATRX proteins, respectively). The binding sites localization of the interactors DMAP-1, CBX5, MECP2, EZH2 and DAXX, which have been characterized in humans, are indicated. **B.** Predicted 3D structures of the RFD from the human DNMT1 and *X. index dnmt1a,b and c*, respectively. Left structure colored by samples and right structure color based on pLDDT scores. **C.** Predicted 3D structures of the ADD from the human ATRX and *X. index atrx,* respectively. Left structure colored by samples and right structure color based on pLDDT scores. **D**. Phylogenetic tree illustrating the presence/absence of canonical DNMT1, 2 and 3, and ATRX proteins in 103 nematode species (the 71 included in the orthofinder analysis and 32 draft nematode genomes from Dorylaimia and Enoplia produced in [26]. Dark blue (DNMTs) and purple (ATRX) represent the presence of the corresponding full-length protein-coding gene. Light blue indicates partial evidence for the presence. *X. index* is highlighted in yellow.

We then projected these findings in the context of the whole Nematoda phylum evolution. Using a combination of orthology search at the protein level, tBLASTn, and HMM analyses, we thus investigated the presence/absence of canonical *DNMTs* and *ATRX* in our dataset of 71 nematode genomes and proteomes to obtain a comprehensive representation of the distribution of these genes in the nematode tree of life. Due to the lack of genomic resources, the most basal nematode clades, Enoplia and Dorylaimia, were under-represented in our initial dataset of well assembled and annotated nematode genomes. Therefore, to more precisely trace the evolutionary history of the epigenetic protein toolkit, we further complemented this study with 60 additional draft nematode genomes produced from a recent phylogenomic analysis of the phylum [26]. We detected the presence of full-length sequences of ATRX only in Enoplia and Dorylaimia, the most basal clades (**Figure 3D, Table S5**). In addition, we identified six species with full-length DNMT1s and seven with partial evidence for DNMT1s (with at least the presence of the RFD domain). Interestingly all these species belong to the Dorylaimida order. No other full-length DNMT1 was detected in other orders. We complemented these results with a DNMT1 phylogeny analysis based on referenced vertebrate and plant sequences, as well as all putative DNMT1 identified to date in nematodes. The resulting phylogeny largely recapitulated the species phylogeny, with the exception of the sequences from *Bursaphelenchus xylophilus* and *Trichinella spiralis* [17,19], which have been previously reported as candidate DNMT1s **(Figure S10).** The outgroup phylogenetic position and lack of canonical DNMT1 domains of *B. xylophilus* and *T. spiralis* sequences strongly suggest these proteins are actually not DNMT1 and only have distant similarities with canonical ones. These results further supported the specific presence of full-length DNMT1 in Dorylaimida only among nematodes examined to date. The unexpected presence of both canonical *DNMT1* and *ATRX* genes in Dorylaimida drove us to investigate the presence of known interactors of these two well-characterized proteins. This analysis revealed a mixed pattern, with some interactors well conserved across nematodes, whereas others appear to be restricted to basal clades. Overall, we found that most of the known interactors of ATRX and DNMT1, including interactants shared between these two proteins, have an ortholog in *X. index* suggesting fully conserved pathways **(Figure S8F)**. Similarly to ATRX, the distribution of canonical DNMT3 proteins was restricted to the basal Enoplia and Dorylaimia clades, suggesting a loss in the last common ancestor of all the other clades. Candidate DNMT2s were found in some Spirurina, Rhabditina and up to some basal Tylenchina, but not in Tylenchymorpha, suggesting a more recent loss (**Figure 3D, Table S5**). Taken together, these findings suggest a sequential loss of epigenetic players including the DNMT repertoire during nematode evolution, introduced by the loss of DNMT1, followed by DNMT3 and, finally, DNMT2.

Finally, we investigated whether the idiosyncratic epigenetic toolkit encoded in the *X. index* genome could be extended beyond DNA and chromatin modifications to the regulation of genome architecture. In mammals, the insulator CCCTC-binding factor (CTCF), along with cohesin, plays crucial roles in the three-dimensional genome organisation [41]. Although we identified components of the cohesin complex such as SMC1 and SMC3 in most nematode orders **(Figure S8F)**, CTCF is not present in *C. elegans* [42]. Thus, using human CTCF as reference, we identified the xiphind_chr_10g011800 gene as *X. index CTCF* ortholog. Its functional annotation was further supported by sequence alignment with human CTCF, which revealed strong conservation of the characteristic array of 11 zinc-finger (ZNF) domains involved in DNA binding **(Figure S11).** Similarly to ATRX, CTCF orthologs were only detected in Dorylaimida and other basal orders. Altogether these findings reveal a broader evolutionary pattern in which the epigenetic toolkit of early-branching nematodes retains components that are absent from broadly studied model species. During nematode evolution, DNMTs, ATRX, and CTCF were probably parts of a radical reshuffling that occurred in the epigenetic toolkit. In this context, these results highlight the key phylogenetic position of the Dorylaimida to more comprehensively investigate epigenetic toolkit shifts in invertebrates.

### CpG DNA methylation around TSS in *X. index* is inversely correlated with chromatin accessibility and gene transcription

The presence of a full DNMT arsenal in *X. index* suggests that DNA CpG methylation might take place in this species, a rare feature in nematodes. Thus, we took advantage of nanopore sequencing data to assess the presence of methylated CpGs in the *X. index* genome [43]. In addition, we performed Assay for Transposase-Accessible Chromatin using Sequencing (ATAC-seq) experiments on three biological replicates of *X. index* to assess the genome-wide chromatin accessibility pattern. DNA methylation within the promoter region being largely associated with gene silencing and reduced chromatin accessibility [44], we investigated the gene expression levels, the methylation status at CpGs and the chromatin accessibility (ATAC-seq) surrounding the transcription start sites (TSS) genome wide. A k-means clustering based on the percentage of methylated CpGs highlighted three clusters (**Figure 4**). In the meta-analysis, cluster 1 contained lowly-expressed genes with a high percentage of CpGs in the TSS region, a high proportion of which being methylated. In line with gene regulation observed in vertebrates [44,45], the chromatin accessibility was relatively low in this cluster. By contrast, cluster 3 included highly expressed genes associated with low to null proportion of methylated CpGs and an increased chromatin accessibility across the TSS regions (**Figure 4).** The observed differences between k-means-based clusters are statistically supported (**Figure S12**). Altogether, these results confirmed the presence of DNA methylation in *X. index* and suggest an association between the level of CpG methylation at TSS, chromatin accessibility status and gene expression, reminiscent of the orchestration described in vertebrates.

**Figure 4:**
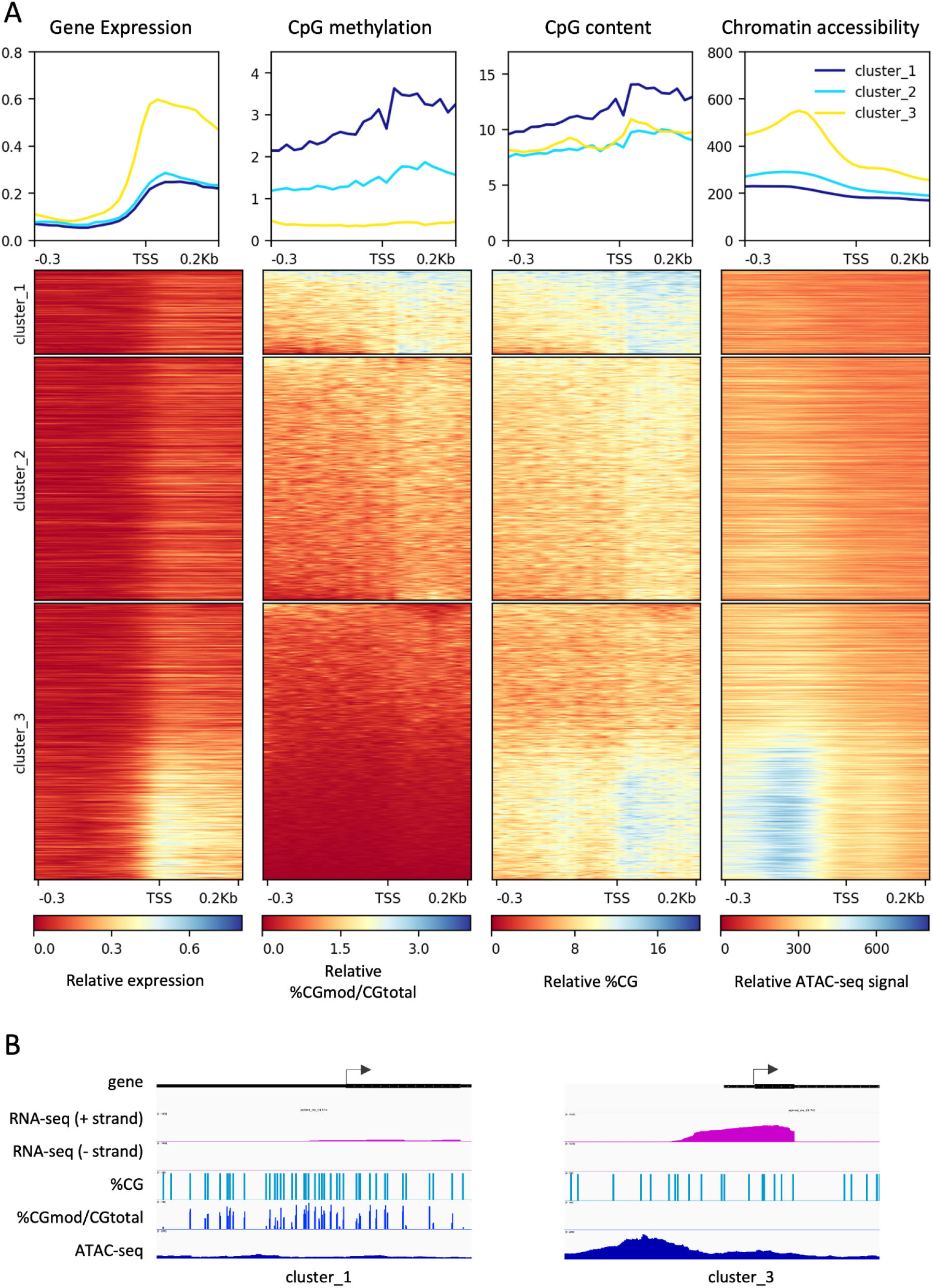
DNA C5 methylation status at CpGs in *X. index* anti-correlates with gene expression and chromatin openness. **A**. heatmaps and metaplots showing the relative levels of gene expression, percentage of CpGs methylated (CGmod) over total CGs, %CGs and chromatin accessibility from 300 bp upstream to 200bp downstream of the start codon of protein coding genes, used as a proxy for TSS, based on k-means clustering performed on CGmod. **B**. Examples of candidate regions (500bp windows) surrounding the TSS (arrow showing the TSS and the gene strand).

## Discussion

In this study, we produced the first T2T genome for the basal nematode order Dorylaimida. Beyond providing an essential genomic resource for *X. index*, a virus vector that threatens the wine industry, our work highlights how sampling poorly explored branches of the tree of life can reveal unexpected evolutionary features. In particular, our study provides evidence for a plant-to-nematode horizontal gene transfer (HGT) and reveals an epigenetic toolkit of unexpected completeness for a nematode, including regulatory features reminiscent of those described in vertebrates.

More than 30,000 protein-coding genes were predicted in the *X. index* genome, and 10% of them encode predicted secreted proteins. Furthermore, ca. 400 proteins were identified as CAZymes. This ensemble of proteins certainly contributes to the parasitic arsenal of *X. index*. Among the protein-coding genes, 224 were putatively acquired via HGTs of non-animal origin, a number consistent with previous estimates in other plant-parasitic nematodes [38]. Expert curation narrowed down the list to 44 genes classified as highly likely HGTs, predominantly originating from bacteria and fungi. Several of them encode CAZyme families known to be involved in plant parasitism, such as GH12 cellulases, GH28 polygalacturonases and GH32 invertases. Such CAZymes with digestive activities on plant sugars have also been previously reported as acquired via HGT in distantly related cyst and root-knot plant-parasitic nematodes, although from different donor species [9–11,37]. The recurrent acquisition of CAZyme families from distinct donors supports the hypothesis that parallel HGT events contributed to the independent emergence of plant parasitism in nematodes. Notably, our analysis also provides, to our knowledge, the first compelling evidence for a plant-to-nematode HGT. Surprisingly, despite the intricate interaction between plants and nematodes, no convincing evidence for HGT events of plant origin has been reported so far in PPNs [38]. The transferred gene in *X. index* encodes a calmodulin that contains a predicted signal peptide for secretion, whereas no signal peptide is present in its closest plant homologs. This raises the possibility that the protein may function as a secreted effector. Many calcium-related processes in eukaryotic organisms depend on calmodulins, which act as sensors by binding to diverse signaling proteins, such as kinases. Among others, calmodulin-dependent protein kinases have been implicated in plant stress response signaling [46]. A recent study in rice showed that a root-knot nematode effector can interact with a host calmodulin and interfere with its interaction with a conserved DNA-binding protein, thereby promoting plant susceptibility [47]. By analogy, the plant-derived calmodulin identified in *X. index* could potentially interfere with endogenous plant calmodulin-dependent signaling in a putative “back to sender” model, an appealing hypothesis to investigate further in future functional studies.

Beyond the analysis of the parasitic arsenal of *X. index*, the most striking feature we uncovered in this study is the composition of the epigenetic toolkit encoded in its genome. In vertebrates, genome-wide CpG methylation mediated by a complete repertoire of DNMTs plays a central role in the epigenetic regulation of gene expression. However, in invertebrates, which represent the vast majority of animals, the picture is much more heterogeneous. In nematodes, complete repertoires of DNMTs and consistent 5mC levels are scarce [16,20]. While entire clades such as Rhabditina and Tylenchida seem to have lost both DNMTs and CpG methylation, the alternative epigenetic marks N6-adenine methylation has been reported in *C. elegans* and, more recently, in root-knot nematodes [18,21,22]. In more basal clades, DNMT1 candidates have been previously claimed. However, our in-depth analysis revealed that all these proteins lack the N-terminal domains characteristic of canonical DNMT1, and most also lack some important C-terminal domains. This suggests that CpG methylation is unlikely to represent a conserved regulatory mechanism across nematodes. In sharp contrast, our thorough analysis of *X. index* points out (i) a complete and expanded DNMT repertoire, (ii) genome-wide CpG methylation and (iii) an inverse association around TSS between 5mC levels and gene expression as well as chromatin accessibility. Collectively, these observations reveal surprising similarities between many aspects of the epigenetic machinery of *X. index* and that of vertebrates [32]. However, as a microscopic plant ectoparasite, which can only be produced in soil with a moderate multiplication rate, *X. index* is a challenging model to work on, especially given the quantity required to generate high-quality genomic, transcriptomic and epigenetic resources. Due to these limitations, our study is based on bulk mixed developmental stages which introduces biological heterogeneity and therefore consequently restricts our analyses to global observations and trends. Thus, beyond the clear pattern observed between methylation, gene expression, and chromatin accessibility around TSS in our results, future studies based on specific separate developmental stages would indubitably refine our knowledge of the involvement of epigenetic modifications in gene regulation during nematode development and parasitism.

The presence of three expressed DNMT1 paralogs in *X. index* is particularly intriguing, as only a single copy is generally present in most vertebrates [48]. One hypothesis is that some paralogs could be specific to particular cell lineages (somatic/germline), mimicking DNMT1 isoforms in humans, or, as for DNMT3, a copy may serve as a regulator of other DNMT1s. The functional significance of this expansion will require in-depth functional studies. Beyond DNA methylation, *X. index* possesses a full-length version of the chromatin remodeler ATRX, an uncommon finding outside vertebrates. This key epigenetic player is well known in research on genetic diseases and cancer. The association between ATRX deficiency and aberrant DNA methylation was first raised decades ago [49] and has been strengthened since then [45,50], suggesting that ATRX and DNMTs may be intrinsically linked to maintain chromatin integrity and the fine tuning of specific genomic targets. Completing this epigenetic toolkit, we identified a putative CTCF-coding gene in the *X. index* genome. This transcription factor and insulator is involved, along with the cohesin complex, in the three-dimensional organization of the genome [41]. While cohesin is broadly conserved across nematodes, CTCF appears to be restricted to basal nematode lineages, mirroring the distribution of ATRX. In mammals, ATRX deficiency alters CTCF occupancy and chromatin interactions [51–54]. The co-occurrence of DNMTs, ATRX and CTCF in *X. index* therefore points to a potentially integrated regulatory framework linking DNA methylation, chromatin state and genome architecture in a manner surprisingly reminiscent of vertebrates. Whether these components functionally interact in *X. index* remains, however, to be determined.

The phylogenetic distribution of DNMTs, ATRX, and CTCF uncovered in this study indicates that a major reshuffling of epigenetic components occurred after the divergence of nematode clades from the Dorylaimida order. In particular, our data are consistent with extensive erosion of this toolkit after the last common ancestor of Chromadorea, which encompasses the Spirurina, Rhabditina and Tylenchina orders, and which can be dated to ca. 482 MYA (408-561 MYA) [26]. Taken together, our results reveal striking differences in epigenetic organization across nematodes. Dorylaimida have retained an epigenetic repertoire that, in terms of gene complement, DNA methylation patterns and associated regulatory features, shows notable similarities to vertebrate systems, whereas members of Chromadorea appear to have undergone extensive loss of these components. These findings probably make Dorylaimida the most basal animal lineage known to possess this full complement of epigenetic regulators and position them as strategic models for studying the evolution and function of epigenetic systems. By contrast, members of the Chromadorea class may have evolved an alternative epigenetic regulation or may share some similarities with other invertebrate organisms. Future studies based on high-quality genomic resources from other early branching nematodes within Chromadorea (*e.g.,* Chromadorida, Monhysterida and Plectida) would likely confirm this shift in the phylum.

## Conclusions

The chromosome-scale genome assembly of *Xiphinema index* provides an unprecedented view of the genomic and epigenetic features of a poorly explored lineage of the Dorylaimida and offers new insights into the evolution of genome regulation in plant-parasitic nematodes. It also represents an essential resource for studying the biology of *X. index* and, ultimately, for understanding the molecular basis of the tripartite interactions between *X. index*, grapevine fanleaf virus and grapevine.

Taken together, our findings reveal two striking features of *X. index*: an unprecedented case of plant-to-nematode horizontal gene transfer and an exceptionally comprehensive epigenetic toolkit within the Nematoda phylum, in terms of gene repertoire completeness, DNA methylation and chromatin accessibility patterns. The latter points to a regulatory landscape sharing notable similarities with that of vertebrates, while highlighting the diversity of epigenetic systems across nematodes. Hence, this species constitutes a key evolutionary model for investigating the origins, plasticity and functional consequences of epigenetic and host–parasite regulatory innovations in animals.

## Methods

### Biological material preparation and sequencings

#### Nematode production and sample preparation

The *Xiphinema index* population ‘Samos’ and ‘Iran’ (both populations were produced independently from a single female) were reared on fig trees at Institut Sophia Agrobiotech’s greenhouse facility, which was maintained between 5-32°C for decades and regularly verified using molecular markers [55,56]. Nematodes were extracted from soil using Oostenbrink elutriator according to the protocol defined by Hooper [57]. For all the experiments related to genome sequencing and assembly, *X. index* individuals, at different stages, were manually picked up to avoid contamination with other nematodes. Approximately 70,000 mixed-stages of *X. index* were collected in 1X antimycotic solution (Sigma) to produce all the samples for long-reads (PACBIO HiFi, ONT) and short-reads (Hi-C, RNA and ATAC) sequencings.

#### High molecular weight DNA extraction and library preparation for long read sequencing

DNA extraction for subsequent long read sequencing was performed using the MasterPure™ Complete DNA and RNA Purification Kit from Biosearch Technologies (formerly epicentre) following supplier’s instructions. Nematodes were ground in liquid nitrogen using a micro pestle directly in the microtube. All homogenisation steps were performed gently to avoid DNA fragmentation. DNA was resuspended in 120 μL of EB buffer (Qiagen). DNA was analyzed for quality and quantity controls using nanodrop, Qubit and fragment analyser.

For PACBIO HiFi sequencing, the final sample used for library preparation contained 16.4 µg of gDNA with an average fragment size of 36,789 bp. Library preparation and sequencing were performed according to the manufacturer’s instructions “Procedure & Checklist Preparing HiFi SMRTbell Libraries using SMRTbell Express Template Prep Kit 2.0”. At each step, DNA was quantified using the Qubit dsDNA HS Assay Kit (Life Technologies). DNA purity was tested using the nanodrop (Thermofisher) and size distribution and degradation assessed using the Femto pulse Genomic DNA 165 kb Kit (Agilent). Purification steps were performed using AMPure PB beads (PacBio). 15µg of DNA was double purified then sheared at 20kb using the Megaruptor3 system (Diagenode). Using SMRTbell Express Template prep kit 2.0 on 5µg of sample, the library was treated with an exonuclease cocktail to digest unligated DNA fragments. A size selection step using a 8kb cutoff was performed on the BluePippin Size Selection system (Sage Science) with “0.75% DF Marker S15-20 kb Improved Recovery” protocol. Using Binding kit 2.0 kit and sequencing kit 2.0, the primer V2 annealed and polymerase 2.0 bounded library was sequenced by diffusion loading onto 1 SMRTcell on Sequel2 instrument at 80pM with a 2 hours pre-extension and a 30 hours movie.

Regarding the ONT sequencing, library preparation and sequencing were performed according to the manufacturer’s instructions “1D gDNA selecting for long reads (SQK-LSK109)”. At each step, DNA was quantified using the Qubit dsDNA HS Assay Kit (Life Technologies). DNA purity was tested using the nanodrop (Thermofisher) and size distribution and degradation assessed using the Fragment analyzer (Agilent) DNF-464 HS Large Fragment Kit. Purification steps were performed using AMPure XP beads (Beckman Coulter). For 1 Flowcell, 3µg of DNA was purified then sheared at 25kb using the megaruptor system (diagenode). Using SQK-LSK 109 kit (ONT) on 2µg of sample, library was loaded onto 1 R9.4.1 flowcell and sequenced on GridION instrument at 30 fmol within 48H. Basecalling was done in super-high accuracy mode with Dorado.

#### Extraction of nuclei, library preparation and illumina sequencing for Hi-C sequencing

Nematodes were crushed for 10 minutes on ice in a dounce tissue grinder tube with 2 ml of extraction buffer containing KCl, NaCl, MgCl2, EDTA, Tris/HCl, Protease inhibitor, sucrose, sodium butyrate, PMSF, DTT, and IGEPAL (at the final concentrations 60 mM, 15 mM, 5 mM, 0.1 mM, 15 mM, 1 g/L, 300 mM, 5 mM, 0.1 mM, 0.5 mM and 0.4%, respectively). The resulting solution was loaded in a Corex centrifugation tube containing 8 mL of a separation buffer made of KCl, NaCl, MgCl2, EDTA, Tris/HCl, Protease inhibitor, sucrose, sodium butyrate, PMSF, and DTT (at the final concentrations 60 mM, 15 mM, 5 mM, 0.1 mM, 15 mM, 1 g/L, 1.2 M, 5 mM, 0.1 mM, and 0.5 mM, respectively). Centrifugation at 10,000 g for 30 minutes at 4°C with slow deceleration was applied. The supernatant was removed entirely. The pellet was carefully resuspended by pipetting in 5 mL of PBS 1X. Then, 500 µL of freshly prepared buffer TC containing NaCl, 100 mM; EDTA-pH8, 1 mM; EGTA-pH8, 0.5 mM; HEPES-pH8, 50 mM and Formaldehyde 22 %, was added. Samples were treated using the reagents of the Arima - Hi-C kit. 289 µL of the Stop Solution 1 (from the kit) was added and the tube was incubated at RT for 5 minutes. The tube was centrifuged at 2000 g for 15 min at RT and the supernatant was removed. At this step an aliquot of 10% of the initial volume was prepared and stored at -80°C for future estimation of the initial quantity of nuclei. Then, the pellet was resuspended in 24 µL of conditioning solution and incubated at 62°C for 10 minutes. The final steps of the protocol followed the instructions provided by the Arima-Hi-C kit (ref. 510008) along with the TruSeq DNA PCR-Free Kit and TruSeq DNA UD Indexes (Illumina, ref. 20015962, ref. 20020590) and the KAPA library amplification kit (Roche, ref. KK2620). The Hi-C library was then sequenced in paired-end 2x150nt mode on a NovaSeq6000 (Illumina).

#### RNA extraction, library preparation and illumina transcriptome sequencing

For the transcriptome, three batches of *X. index* were soaked either in tap water, in 0.2 M NaCl for 1h or in 0.1% acetic acid for 1 h. The combination of standard and stressed individuals was made to maximize the number of expressed genes. The samples were ground in liquid nitrogen using a micro pestle directly in the microtube. 800 µl of extraction buffer (CTAB 2.5%, PVPP 2%, Tris-HCL 100mM, EDTA 25 mM, NaCl 2M, β-mercaptoethanol 2%) was added to the tube. After 30 minutes of incubation at 65°C, 800 µl of Chloroform/Isoamyl alcohol (CI; v/v; 24/1) was added, and after homogenization, the tube was centrifuged (16,000g - 8 min - 4°C). The supernatant was retrieved and the same volume of water-saturated Phenol (pH 4.5-5) / Chloroform / Isoamyl alcohol (PCI; v/v; 25/24/1) (ca 700 µl) was added and centrifuged. A second step with CI was carried out and the supernatant was retrieved and mixed with 500 µl NaCl 5M and 500 µl of isopropanol and stored overnight at -20°C. After centrifugation (16,000g - 20 min - 4°C), two cleaning steps were carried out with ethanol 70%. The pellet was resuspended in 30 µl of buffer EB (Qiagen). Genomic DNA was removed from the sample using the kit TURBO DNA-free (Ambion) following the supplier’s instructions. Samples purity and quality were assessed with a nanodrop and Qubit before library preparation. RNA-seq libraries have been prepared according to Illumina’s protocols using the Illumina TruSeq Stranded mRNA sample prep kit to analyze mRNA. Briefly, mRNA were selected using poly-T beads. Then, RNAs were fragmented to generate double stranded cDNA and adaptors were ligated to be sequenced. 11 cycles of PCR were applied to amplify libraries. Library quality was assessed using a Fragment Analyser and libraries were quantified by QPCR using the Kapa Library Quantification Kit. RNA-seq experiments have been performed on an Illumina NovaSeq 6000 using a paired-end read length of 2 × 150 bp with the Illumina NovaSeq. 6000 sequencing kits.

#### Chromatin accessibility assay and data analysis

For ATAC-seq experiments, three replicates of mixed stages *X. index* were collected after overnight filtration on tissues and washed with 1X PBS before flash frozen in liquid nitrogen and stored at -80°C before nucleic acid extractions. ATAC-seq experiments were performed using ATAC-Seq Kit from Active Motif (53150) following manufacturer’s instructions. Briefly nuclei from *X. index* were extracted with Douncer in ATAC lysis buffer before being filtered on 40 µm followed by 10 µm mesh strainers. 200 000 nuclei were tagmented in Tagmantation Master Mix for 30 min at 37°C at 800 rpm. Fragmented DNA was purified using DNA purification columns. PCR amplification and indexing was performed using i7 and i5 Indexed Primers. Samples were then purified with SPRI Beads. Library quality was assessed using a Fragment Analyser and sequenced using a paired-end read length of 2 × 75 bp with AVITIDY chemistry.

### The genome assembly of *X. index*

#### Genome size, ploidy and heterozygosity estimation

We used JellyFish version 2.3.0 [58] with a k-mer size of 21 to enumerate k-mers on the *X. index* HiFi reads. We generated histograms using the jellyfish histo command and adjusted the -h parameter to allow k-mer repeated up to 1 million times. To infer the ploidy level, we used SmudgePlot (v.0.2.5) [59] with default parameters. Finally, to estimate genome size and heterozygosity, we used GenomeScope2 [59] with the previously obtained ploidy levels and k-mer histograms.

#### Genome assembly and scaffolding

First, Nanopore reads were filtered for a minimum quality of Q10 and a minimum length of 30kb using Nanofilt 2.8.0 [60]. Then, we used HiFiasm [61] version 0.19.8 with the PacBio HiFi long reads combined with the Nanopore longest reads (-ul option) and the Hi-C Illumina paired-end short reads (-h1, -h2 options) with the other parameters left to default. We used TiDK (v0.1.5) [62] to search the canonical nematode telomeric repeat ‘TTAGGC’ in the primary contigs assembly (detection window of 1kb). Then we used Juicer (v 1.6) from 3D-DNA [63] to produce initial contact maps and scaffolding, using Hi-C data. The final scaffolding was manually edited using JuiceBox (v2.13.7) with the help of nematode telomere motifs identified using TIDK (converted in wig files). Scaffolding of the HiFi contigs with HiC data introduced gaps, so we attempted to fill the gaps with the ONT data. For that, we used tgs-gapcloser v1.2.1 [64] without error correction and all the other parameters left to default. This tool resolved 51 % of the gaps in the pseudo-chromosomes. To identify and remove debris that could represent duplicated information already present in the pseudo-chromosomes, we used minimap2 v2.24 [65] to map the debris against all the chromosomes. Then, we calculated the debris coverage using bedtools v2.30.0 genomecov [66] and removed the debris that aligned on more than 90% of their length on pseudo-chromosomes. We used BlobTools [67,68] to identify and remove mitochondrial scaffolds as well as possible contaminations. Because there is a lack of closely related nematode sequence data in public libraries, two sources of evidence were used as hit files. A diamond (v 2.0.8) blastx search against the NCBI’s NR library as a first source and a NCBI blastn search against the NT library as a second source, both with an e-value threshold of 1e-25 (options were -k 5 -e 1e-25 -b 6 -c 1 --sensitive -F 15 --range-culling). Finally, pseudo-chromosomes were ordered by size and orientated according to telomere repeats, starting with reverse motif GCCTAA and ending with forward motif TTAGGC. A schematic representation of the assembly generation of the *X. index* genome is available in **Figure S13**.

#### Mitochondrial genome assembly and NUMT

The mitochondrial genome was assembled using mitochondrial circular DNA reconstitution (https://github.com/GDKO/aladin; ALADIN v1.1) with -d N -m M options. The seed used for the assembly was the NADH dehydrogenase subunit 4 (ND4) partial gene sequence that was available from the same *X. index* population (NCBI accession number: LT996799). The assembly was performed using PacBio HiFi reads and verifications were conducted by aligning ONT reads. NUMTs were detected using a Blastn approach between the mitogenome and the nuclear genome assembly and results were filtered by alignment size (>500 bp) and percentage of identity (>90%). The main NUMT detected in chromosome 10 was validated using specific primers. Two sets of primers were designed to flank the 5’ and the 3’ regions of the insertion: RWI5’_F2: CAGTCTCTTTGGCCTGGTCT / RWI5’_R2: CGGGGATCCTTTACTGTTTCA and RWI3’_F1: TCCAAAACATGATCCGCTCT / RWI3’_R1: GTTGGAATCCTGACCTGGGT. MyTaq™ red mix kit was used following the supplier’s instructions. Genomic DNA of two different populations of *X. index* (‘Samos’, which is the one we have sequenced and ‘Iran’) were used to perform PCR (94°C-5min; 35 cycles of 94°C-30s, 58°C-30s, 72°C-1min; 72°C-5min) and gave amplicons of 470bp and 413bp for *X. index* Iran’ and 494bp and 425bp for *X. index* ‘Samos’. PCR products were sequenced and aligned to the genome sequence for final validation.

#### Gene models

Gene models prediction was done with the fully automated pipeline EuGene-EP (v2.0.3) [69]. EuGene has been configured to integrate similarities with known proteins of Caenorhabditis elegans (PRJNA13758) from WormBase Parasite [70] and “nematoda” section of UniProtKB/Swiss-Prot library [71], with the prior exclusion of proteins that were similar to those present in RepBase [72]. The dataset of Xiphinema index transcribed sequences generated in this study were aligned on the genome and used by EuGene as transcription evidence. For this, we first assembled de novo using Trinity [73] the transcriptome of *X. index* obtained from the one above-described condition and for a given trinity locus we only retained the transcript returning the longest ORF. Finally, only de novo assembled transcripts that aligned on the genome on at least 30% of their length with at least 97% identity were retained. The EuGene default configuration was edited to set the “preserve” parameter to 1 for all datasets, the “gmap_intron_filter” parameter to 1 and the minimum intron length to 35 bp. Finally, the Nematodes-specific Weight Array Method matrices were used to score the splice sites (available at this URL: http://eugene.toulouse.inrae.fr/WAM/).

#### Functional annotation, BUSCO completeness and orthology

We used InterProScan v5.68-100.0 [74] to detect conserved domains in the predicted protein set of *X. index*. We used the options --goterms and -pa to allow assignment of corresponding Gene Ontology (GO) terms as well as prediction of corresponding MetaCyc and Reactome pathways.

We used BUSCO v5.3.1 with the metazoa.odb10 dataset and default parameters to estimate completeness both at the genome assembly level (--genome) and on the set of predicted proteins (--proteome). We compared *X. index* with 5 other Dorylaimia based on the following criteria. From the WormBase ParaSite [70] resource, we selected Dorylaimia genomes with a minimum of 70% BUSCO [75] nematoda completeness at the genome level according to WormBase. For each genus, we selected up to two species that presented the highest N50 at the genome assembly level. This ended up with a selection of five species: *Trichinella spiralis* [76], *Trichinella nelsoni* [77], *Trichuris muris* [78]*, Trichuris trichiura* [78], and *Mesodorylaimus sp. YZB*2_*4* [79].

We used OrthoFinder v2.5.4 [80] to identify homologs of *X. index* predicted proteins in other nematode predicted proteomes. We started from a previous OrthoFinder analysis that included 68 nematode predicted proteomes and one tardigrade proteome as an outgroup [34]. In brief, the list was made according to the genome completeness and the representativity of the different nematode clades. A tardigrade was used as an outgroup as tardigrades are the closest group of animals to nematodes. We added the *X. index* proteome as well as those of *Trichuris trichiura*, and *Mesodorylaimus sp. YZB2_4* that were not previously included. The full list of species is available in **Table S2.** We used the option -M msa to generate multiple-sequence alignments and phylogenetic trees for all orthogroups. By analyzing the produced orthogroups we then identified *X. index* proteins conserved with other nematodes as well as those specific to this species.

### Transcription, chromatin accessibility and methylome analyses

To determine the expression of protein-coding genes, we used RNA-seq data mapped to the genome with STAR (Version 2.7.8a) using more stringent end-to-end parameters [81]. Expected read counts, FPKM and TPM values were calculated on the predicted genes from the *Xiphinema index* GFF annotation using RSEM version 1.3.1 [82]. We considered as supported by transcription data any gene with an FPKM equal or higher than the value of the first quartile of the FPKM distribution for the whole set of predicted genes. BigWig (bw) files mapped RNA-seq data were generated with a data normalization by Counts Per Million mapped reads (CPM) using Deeptools (version 3.5.6) bamCoverage with the options -bs 1 --normalizeUsing CPM --ignoreDuplicates. Similarly strand specific bw files were generated with the extra option --filterRNAstrand forward or --filterRNAstrand reverse. Data were visualized using IGV 2.16.2.

ATAC-seq data were trimmed with fastp (v0.23.4) and mapped using Bowtie2 (v2.5.1). Bam files were generated with samtools (v1.20). bigwig files were generated using bamCoverage from deeptools (deeptools_3.5.6--pyhdfd78af_0) with the options -bs 1 -p 4 --normalizeUsing RPKM --ignoreDuplicates and visualized on IGV 2.16.2.

The detection of methylated CpGs was performed by basecalling the nanopore long-read sequencing data with the option --modified-bases 5mCG_5hmCG and aligning the reads with dorado v0.9.0 on *X. index* assembly. A bedMethyle file was generated using Modkit v0.4.3 pileup with the option --cpg. The methylated CpG data were extracted from the bedMethyle and CpG with a score lower than 5 (Nvalid_cov) were removed. CpG with only one call as modified (Nmod=1) were considered as background and thus considered as unmodified for downstream analyses. Bw files were generated from bedgraph of the percentage of CGmethylated/CGtotal (%CGmod/CGtotal) and percentage of CG (%CG) using ucsc bedGraphToBigWig (ucsc-bedgraphtobigwig:445--h954228d_0) and visualised with IGV (2.16.2). The association between the relative level of gene expression and the status of CpG methylation was assessed by isolating the translation start sites (ATG) of protein coding genes from the gene annotation analyses. Overlapping genes were excluded from the analysed list to limit the generation of artefacts in the downstream analyses. Metaplots and heatmaps of the -300/+200bp surrounding the ATG as a proxy for the transcription start site (TSS) region were performed using the matrix generated with Deeptools (v3.5.6) computeMatrix using the options reference-point --beforeRegionStartLength 300 --afterRegionStartLength 200 -bs 20 –missingDataAsZero --transcript_id_designator “transcript_id”. Metaplots and heatmaps were plotted using plotHeatmap. A k-means clustering based on the %CGmod/CGtotal was performed using the --kmeans and --clusterUsingSamples options and the regions were sorted based on descending values of %CGmod/CGtotal using --sortRegions descend and --sortUsingSamples options. Violin plots of the -300/+200bp surrounding the TSS splitted by the k-means algorithm based clusters were generated based on the Metaplot and heatmap datasets using ggplot2 (v3.5.0) and statistical tests were performed using wilcox.test() on R (v4.3.3).

### Genome mining for key epigenetic players

#### Identification of candidate DNMT1 proteins and phylogenetic analysis

We used complementary proteome and genome-based strategies to identify candidate DNMT1 genes and proteins in the 71 nematode species analyzed. First, we used HMMER to search the PANTHER domain PTHR10629 with an e-value threshold of 1e-80. Second, we checked whether orthologs of the candidate DNMT1 identified by HMM were present in other nematodes using the orthogroups generated by OrthoFinder. Third, we used the nematode protein with the lowest e-value returned by HMM as a query to search using tBLASTn the 71 genome sequences. This last step was to make sure the absence of identification at the protein level was not simply due to lack of prediction of the gene model in the genome. To further extend the nematode dataset of candidate DNMT1 proteins, we repeated this same tBLASTn search in the 60 draft genomes that cover 11 orders, recently produced as part of a comprehensive phylogenomics analysis of the phylum Nematoda [26]. The tBLASTn match regions were extracted and we used AUGUSTUS and FGENSEH+ [83] with the query protein to predict genes and the corresponding amino acid sequences at the corresponding loci. The amino acid sequences were then analysed with interProScan to assess the tBLASTn results and find the corresponding protein domains. In the end, we complemented the collection of candidate Ecdysozoa DNMT1 proteins identified in [16] by the newly identified ones. For consistency of inclusion criteria, we only kept those proteins which had a match with the PTHR10629 PANTHER domain with an e-value threshold of 1e-80. The same approach was used to identify ATRX, DNMT2, DNMT3 and CTCF orthologs. Identified candidates were screened for presence of the following canonical domains : DNMT1-RFD (PF12047), BAH (PF01426), CYTOSINE-SPECIFIC METHYLTRANSFERASE (PTHR10629), ADD (PF17981), SNF2 (PF00176), HELICASE (PF00271), TRNA C5METHYLTRANSFERASE (PTHR46098), PWWP (PF00855), ADD_dnmt3 (PF17980 / PF21255), and DNA METHYLASE (PF00145).

Orthologs of ATRX and DNMT1 interactors in *X. index* were searched using reciprocal blastP analysis with the human sequences as queries. Then, the *X. index* ortholog sequences identified were used as queries in BlastP analysis against ClusteredNR database restricted or not to Nematoda (taxid:6231).

#### 3D protein structure prediction

Interproscan annotated RFD domain from DNMT1 and ADD domain from ATRX were retrieved from *H. sapiens* and *X. index* amino acid sequences. Fasta sequences were then analysed using AlphaFold3 with default parameters but by setting seed. Three independent predictions were made for each domain sequence to assess reproducibility before running Matchmaker on UCSF ChimeraX (version 1.12).

### Carbohydrate-active enzymes (CAZymes) and horizontal gene transfer (HGT) detection

#### CaZome prediction

All the predicted proteins from the *X. index*, *Mesodorylaimus sp. YZB2_4*, *Trichinella spiralis*, *Trichinella nelsoni*, *Trichuris muris*, and *Trichuris trichiura* genomes were compared with entries in the CAZy database [35]. A homemade pipeline combining the BlastP (https://blast.ncbi.nlm.nih.gov/Blast.cgi) and HMMER3 (http://hmmer.janelia.org/) tools was used to compare protein models with the sequences of the CAZy modules. Proteins with E-values less than 0.1 were further screened by a combination of BlastP searches against libraries generated from the sequences of the catalytic and non-catalytic modules. HMMER3 was used to search against a collection of custom Hidden Markov Model (HMM) profiles constructed for each CAZy family. Expert curators then performed manual inspection to resolve borderline cases.

#### Detection of HGT candidates

The *X. index* proteome was searched against the NCBI non-redundant (NR) protein database using DIAMOND (v2) [84] (more-sensitive mode, e-value threshold 1.0e^−3^, maximum 500 hits). The results were submitted to AvP [85], ignoring self-hits to Longidoridae (txid46001). AvP computes the Alien Index (AI) and the Aggregate Hit Support (AHS) for each query sequence, based respectively on the single best animal (NCBI:txid33208) and non-animal hits and on the normalized sum of their scores. Proteins with an AI or AHS > 0, i.e. more similar to non-animal than to animal sequences, were then processed by AvP as follows:. (i) queries were clustered according to shared hits (50% overlap threshold instead of the default 70%); (ii) hit sequences were retrieved from NR (the 50 best hits if no animal hit was found, otherwise all hits preceding the first animal hit plus the next 20); (iii) sequences were aligned using MAFFT (v7) (--auto option) [86] and phylogenies inferred with FastTree (v2) [87]; (iv) HGT candidates and their putative origin were identified from the taxonomy of the sister and ancestral sister branches; (v) HGT-supporting topologies were compared with a constrained topology forcing the monophyly of query and animal sequences using an approximately unbiased (AU) test [88]; and (vi) a local score (-1 to +1) was computed from the genomic environment, negative when flanking genes are themselves HGT candidates (indicating a possible contamination, duplications after an HGT event, or multiple HGTs) and positive when they are likely to be vertically inherited (scaffolds with ≥ 5 genes only).

#### Validation of HGT candidates

HGT candidates from AvP were manually discarded if:,(i) fewer than three donors were present in the sister plus ancestral sister branches; (ii) the HGT-supporting topology was not significantly more likely than the constrained alternative one, unless the animal sequences were taxonomically misannotated (contamination by a donor species) or themselves likely acquired by HGT (BLASTP r against NR); (iii) at least one donor sequence shared > 70% identity with the HGT candidate (possible contamination) (iv) identity with donor sequences was < 30% (low quality alignments); (v) the average alignment length in the DIAMOND homology search results was <100 amino acids with < 50% coverage; or (vi) the local score was < 0 without evidence of duplication after an HGT event or of multiple HGTs, indicating a possible contamination.

#### Phylogeny reconstruction for validated HGT candidates

For each validated HGT candidate, the sequences of all 500 Diamond hits were retrieved by AvP, clustered with CD-HIT at 90% identity [89] and aligned with MAFFT (v7) (--auto) [86]. Highly divergent sequences were removed with trimAl (v1.4) (-resoverlap 0.8 -seqoverlap 75) [90], followed by poorly aligned regions (-automated1). Alignments that had lost all *X. index* sequences or retained < 100 sites were discarded. After removal of duplicate sequences with SeqKit (v2) (rmdup) [91], maximum likelihood trees were inferred with IQ-TREE (v2) with automated model selection as implemented in ModelFinder [92]. Support values were based on a Shimodaira-Hasegawa approximate likelihood ratio test (SH-aLRT) [93] combined with an ultrafast bootstrap (Ufboot2) approximation with 1,000 replicates [94].

Only support values greater than or equal to 80% and 90% for SH-aLRT and UFboot, respectively, were considered. Phylogenies were visualized with iTol [95].

When Metaozan sequences, other than from *X. index*, were present in the phylogenetic tree, we forced a constrained topology supporting monophyly of animal sequences, and determined whether the unconstrained topology was significantly more likely than the constrained alternative one using an approximately unbiased (AU) test [96] with IQ-TREE (v2).

#### PCR validation of the plant-derived HGT

A molecular marker named HGT1 was designed in the first intron of the gene xiphind_chr_06g025050 (putative HGT from plants). We amplified HGT1 using the following primers HGT1-F: GCCACGTAAAGCTCAGCAAT and HGT1-R: GAGTCCAGACCCAAATCCCA with a classical PCR program (tm=59°C; 35 cycles) and MyTaq™ red mix kit. Genomic DNA of two different populations of *X. index* (‘Samos’, which is the one we have sequenced and ‘Iran’) were used as template.

## Data availability

All the raw data produced and used in this study are available in the Bioproject PRJEB75225 (https://www.ebi.ac.uk/ena/browser/view/PRJEB75225). This includes the RNAseq data (ERR12949737), the PacBio Hifi data (ERR14872773), the Hi-C data (ERR14872772), the ONT data (ERR14872784) and the ATAC-seq data (ERR16714051, ERR16714052, ERR16714053). The full set of analyses including the final assembly with the functional annotation are available at https://entrepot.recherche.data.gouv.fr/dataverse/XiphiGeno. The genome resources related to *Xiphinema index* can be viewed at the BIG portal (https://big.plantbios.sophia.inrae.fr).

## Acknowledgements

The authors want to thank Christophe Klopp for fruitful discussions and help in Hi-C contact map analysis. Authors are thankful to Carole Belliardo, Georgios D Koutsovoulos and Djampa K L Kozlowski for help in HGT detection, preliminary genome assembly assessment, and basecalling on ONT data, respectively.

All library preparations and sequencings were performed at the GeT-PlaGe core facility (https://doi.org/10.17180/nvxj-5333), INRAE Toulouse. The authors are grateful to the bioinformatics and genomics platform, BIG, Sophia Antipolis (ISC plantBIOs, https://doi.org/10.15454/qyey-ar89) for computing and storage resources and the genotoul bioinformatics platform Toulouse Occitanie (Bioinfo Genotoul, https://doi.org/10.15454/1.5572369328961167E12) for computing resources. This work was supported by France Génomique National infrastructure, funded as part of “Investissement d’avenir” program managed by Agence Nationale pour la Recherche (ANR) (contract ANR-10-INBS-09), the French government through the France 2030 investment plan managed by the ANR, as part of the Initiative of Excellence Université Côte d’Azur under reference number ANR-15-IDEX-01 and ANR-13-JSV7-0006 ASEXEVOL for funding the sequencing.

## Authors contributions

JT: Epigenetic experiments and data analyses, contributed to the final manuscript KRS: Genome assembly, data analyses, contributed to the final manuscript

AP: Genome assembly, data analyses

DC: HGT predictions and analyses, contributed to the final manuscript

CR: Transcriptome assembly and gene prediction, mitochondrial genome assembly, contributed to the final manuscript, data submission

ED: Predictions and analysis of CAZymes MDR: Transcriptomic analyses

LPB: Production of genomic material ES: Data analyses

UJP: Nematode production

CLR: Sequencing supervision, contributed to the final manuscript CI: Performed Hi-C library and sequencing

AS: Bioinformatic pre-treatment of ONT data MG: Performed PACBIO library and sequencing

RB: Performed ONT libraries and sequencing

LL: Quality controls and sequencing of ATAC libraries DE: Nematode production and expertise

EGJD: Project coordination, funding acquisition, manuscript writing, data analyses

CVG: Nematode production, Project coordination, production of genomic and transcriptomic material, molecular biology validations, manuscript writing, data analyses, data submission

## References

1. Kiontke K, Fitch DHA. Nematodes. Current Biology. 2013;23:R862–4. 10.1016/j.cub.2013.08.009

2. Bardgett RD, van der Putten WH. Belowground biodiversity and ecosystem functioning. Nature. Nature Publishing Group; 2014;515:505–11. 10.1038/nature13855

3. Hoogen J van den, Geisen S, Routh D, Ferris H, Traunspurger W, Wardle DA, et al. Soil nematode abundance and functional group composition at a global scale. Nature. 2019;572:194–8. 10.1038/s41586-019-1418-6

4. van Megen H, van den Elsen S, Holterman M, Karssen G, Mooyman P, Bongers T, et al. A phylogenetic tree of nematodes based on about 1200 full-length small subunit ribosomal DNA sequences. Nematology. 2009;11:927–50.

5. Ahmed M, Roberts NG, Adediran F, Smythe AB, Kocot KM, Holovachov O. Phylogenomic Analysis of the Phylum Nematoda: Conflicts and Congruences With Morphology, 18S rRNA, and Mitogenomes. Frontiers in Ecology and Evolution [Internet]. 2022 [cited 2022 Jan 31];9. https://www.frontiersin.org/article/10.3389/fevo.2021.769565. Accessed 31 Jan 2022

6. Qing X, Zhang YM, Sun S, Ahmed M, Lo W-S, Bert W, et al. Phylogenomic Insights into the Evolution and Origin of Nematoda. Systematic Biology. 2024;syae073. 10.1093/sysbio/syae073

7. Holterman M, Karegar A, Mooijman P, Megen H van, Elsen S van den, Vervoort MTW, et al. Disparate gain and loss of parasitic abilities among nematode lineages. PLOS ONE. 2017;12:e0185445. 10.1371/journal.pone.0185445

8. Bird DM, Jones JT, Opperman CH, Kikuchi T, Danchin EGJ. Signatures of adaptation to plant parasitism in nematode genomes. Parasitology. 2015;142:S71–84. 10.1017/S0031182013002163

9. Danchin EG, Rosso MN, Vieira P, de Almeida-Engler J, Coutinho PM, Henrissat B, et al. Multiple lateral gene transfers and duplications have promoted plant parasitism ability in nematodes. Proceedings of the National Academy of Sciences of the United States of America. 2010;107:17651–6.

10. Haegeman A, Jones JT, Danchin EG. Horizontal gene transfer in nematodes: a catalyst for plant parasitism? Mol Plant Microbe Interact. 2011;24:879–87. 10.1094/MPMI-03-11-0055

11. Lai C-K, Lee Y-C, Ke H-M, Lu MR, Liu W-A, Lee H-H, et al. The Aphelenchoides genomes reveal substantial horizontal gene transfers in the last common ancestor of free-living and major plant-parasitic nematodes. Molecular Ecology Resources. 2023;23:905–19. 10.1111/1755-0998.13752

12. Danchin EGJ, Perfus-Barbeoch L, Rancurel C, Thorpe P, Da Rocha M, Bajew S, et al. The Transcriptomes of Xiphinema index and Longidorus elongatus Suggest Independent Acquisition of Some Plant Parasitism Genes by Horizontal Gene Transfer in Early-Branching Nematodes. Genes. 2017;8:287. 10.3390/genes8100287

13. Jones PA. Functions of DNA methylation: islands, start sites, gene bodies and beyond. Nat Rev Genet. Nature Publishing Group; 2012;13:484–92. 10.1038/nrg3230

14. Luo C, Hajkova P, Ecker JR. Dynamic DNA methylation: In the right place at the right time. Science. American Association for the Advancement of Science; 2018;361:1336–40. 10.1126/science.aat6806

15. Brethouwer T, de Mendoza A, Bogdanovic O. Non-CG DNA methylation in animal genomes. Nat Genet. Nature Publishing Group; 2025;57:2395–407. 10.1038/s41588-025-02303-1

16. Engelhardt J, Scheer O, Stadler PF, Prohaska SJ. Evolution of DNA Methylation Across Ecdysozoa. J Mol Evol. 2022;90:56–72. 10.1007/s00239-021-10042-0

17. Gao F, Liu X, Wu X-P, Wang X-L, Gong D, Lu H, et al. Differential DNA methylation in discrete developmental stages of the parasitic nematode Trichinella spiralis. Genome Biology. 2012;13:R100. 10.1186/gb-2012-13-10-r100

18. Greer EL, Blanco MA, Gu L, Sendinc E, Liu J, Aristizábal-Corrales D, et al. DNA Methylation on N6-Adenine in C. elegans. Cell. Elsevier; 2015;161:868–78. 10.1016/j.cell.2015.04.005

19. Liu Z, Li Y, Zhang X. DNA methylation on C5-Cytosine and N6-Adenine in the Bursaphelenchus xylophilus genome. BMC Genomics. 2023;24:671. 10.1186/s12864-023-09783-7

20. Rošić S, Amouroux R, Requena CE, Gomes A, Emperle M, Beltran T, et al. Evolutionary analysis indicates that DNA alkylation damage is a byproduct of cytosine DNA methyltransferase activity. Nat Genet. Nature Publishing Group; 2018;50:452–9. 10.1038/s41588-018-0061-8

21. Dai D, Liao Y, Bo D, Siddique S, Zhou Y, Hu B, et al. 6mA demethylase Mi-NMAD-1 modulates nematode development and parasitism through the transcription factor Mi-nhr-118 [Internet]. bioRxiv; 2025 [cited 2026 Jan 29]. p. 2025.12.16.694795. 10.64898/2025.12.16.694795

22. Dai D, Zhang S, Hu B, Zhou Y, Cui S, Sun J, et al. N6-methyladenine DNA modification modulates pathogen virulence in nematodes. Proceedings of the National Academy of Sciences. Proceedings of the National Academy of Sciences; 2026;123:e2525035123. 10.1073/pnas.2525035123

23. Clapier CR, Iwasa J, Cairns BR, Peterson CL. Mechanisms of action and regulation of ATP-dependent chromatin-remodelling complexes. Nat Rev Mol Cell Biol. Nature Publishing Group; 2017;18:407–22. 10.1038/nrm.2017.26

24. Garrick D, Samara V, McDowell TL, Smith AJH, Dobbie L, Higgs DR, et al. A conserved truncated isoform of the ATR-X syndrome protein lacking the SWI/SNF-homology domain. Gene. 2004;326:23–34. 10.1016/j.gene.2003.10.026

25. Argentaro A, Yang J-C, Chapman L, Kowalczyk MS, Gibbons RJ, Higgs DR, et al. Structural consequences of disease-causing mutations in the ATRX-DNMT3-DNMT3L (ADD) domain of the chromatin-associated protein ATRX. Proceedings of the National Academy of Sciences. Proceedings of the National Academy of Sciences; 2007;104:11939–44. 10.1073/pnas.0704057104

26. Qing X, Zhang YM, Sun S, Ahmed M, Lo W-S, Bert W, et al. Phylogenomic Insights into the Evolution and Origin of Nematoda. Struck T, editor. Systematic Biology. 2024;syae073. 10.1093/sysbio/syae073

27. Esmenjaud D. Les nématodes de la vigne. Ravageurs de la vigne. Bordeaux: Editions Féret; 2008. p. 21–45.

28. Hewitt W, Raski D, Goheen A. Nematode Vector of Soil-Borne Fanleaf Virus of Grapevines. Phytopathology. St Paul: Amer Phytopathological Soc; 1958;48:586–95.

29. Andret-Link P, Laporte C, Valat L, Ritzenthaler C, Demangeat G, Vigne E, et al. Grapevine fanleaf virus: still a major threat to the grapevine industry. Journal of Plant Pathology. 2004;86:183–95. 10.2307/41992424

30. Nguyen VC, Villate L, Gutierrez-Gutierrez C, Castillo P, Van Ghelder C, Plantard O, et al. Phylogeography of the soil-borne vector nematode Xiphinema index highly suggests Eastern origin and dissemination with domesticated grapevine. Sci Rep. Nature Publishing Group; 2019;9:7313. 10.1038/s41598-019-43812-4

31. Jones JT, Haegeman A, Danchin EGJ, Gaur HS, Helder J, Jones MGK, et al. Top 10 plant-parasitic nematodes in molecular plant pathology. Molecular Plant Pathology. 2013;14:946–61. 10.1111/mpp.12057

32. Long HK, Sims D, Heger A, Blackledge NP, Kutter C, Wright ML, et al. Epigenetic conservation at gene regulatory elements revealed by non-methylated DNA profiling in seven vertebrates. Ferguson-Smith A, editor. eLife. eLife Sciences Publications, Ltd; 2013;2:e00348. 10.7554/eLife.00348

33. Dalmasso A. Cytogenetics and Reproduction in Xiphinema and Longidorus. Nematode Vectors of Plant Viruses [Internet]. Springer, Boston, MA; 1975 [cited 2017 Aug 24]. p. 139–51. 10.1007/978-1-4684-0841-6_9

34. Mota APZ, Koutsovoulos GD, Perfus-Barbeoch L, Despot-Slade E, Labadie K, Aury J-M, et al. Unzipped genome assemblies of polyploid root-knot nematodes reveal unusual and clade-specific telomeric repeats. Nat Commun. Nature Publishing Group; 2024;15:773. 10.1038/s41467-024-44914-y

35. Drula E, Garron M-L, Dogan S, Lombard V, Henrissat B, Terrapon N. The carbohydrate-active enzyme database: functions and literature. Nucleic Acids Research. 2022;50:D571–7. 10.1093/nar/gkab1045

36. Li J, Wiebenga A, Lipzen A, Ng V, Tejomurthula S, Zhang Y, et al. Comparative Genomics and Transcriptomics Analyses Reveal Divergent Plant Biomass-Degrading Strategies in Fungi. Journal of Fungi. Multidisciplinary Digital Publishing Institute; 2023;9:860. 10.3390/jof9080860

37. Danchin EGJ, Guzeeva EA, Mantelin S, Berepiki A, Jones JT. Horizontal Gene Transfer from Bacteria Has Enabled the Plant-Parasitic Nematode Globodera pallida to Feed on Host-Derived Sucrose. Molecular Biology and Evolution. 2016;33:1571–9. 10.1093/molbev/msw041

38. Ko I, Kranse OP, Senatori B, Eves-van den Akker S. A Critical Appraisal of DNA Transfer from Plants to Parasitic Cyst Nematodes. Molecular Biology and Evolution. 2024;41:msae030. 10.1093/molbev/msae030

39. Aguilera P, López-Contreras AJ. ATRX, a guardian of chromatin. Trends in Genetics. 2023;39:505–19. 10.1016/j.tig.2023.02.009

40. Villard L, Fontès M, Ewbank JJ. Characterization of xnp-1, a Caenorhabditis elegans gene similar to the human XNP/ATR-X gene. Gene. 1999;236:13–9. 10.1016/S0378-1119(99)00279-6

41. Dehingia B, Milewska M, Janowski M, Pękowska A. CTCF shapes chromatin structure and gene expression in health and disease. EMBO Rep. 2022;23:EMBR202255146. 10.15252/embr.202255146

42. Sawh AN, Mango SE. Chromosome organization in 4D: insights from *C. elegans* development. Current Opinion in Genetics & Development. 2022;75:101939. 10.1016/j.gde.2022.101939

43. Halliwell DO, Honig F, Bagby S, Roy S, Murrell A. Double and single stranded detection of 5-methylcytosine and 5-hydroxymethylcytosine with nanopore sequencing. Commun Biol. Nature Publishing Group; 2025;8:243. 10.1038/s42003-025-07681-0

44. Blattler A, Farnham PJ. Cross-talk between Site-specific Transcription Factors and DNA Methylation States*. Journal of Biological Chemistry. 2013;288:34287–94. 10.1074/jbc.R113.512517

45. Truch J, Downes DJ, Scott C, Gür ER, Telenius JM, Repapi E, et al. The chromatin remodeller ATRX facilitates diverse nuclear processes, in a stochastic manner, in both heterochromatin and euchromatin. Nat Commun. Nature Publishing Group; 2022;13:3485. 10.1038/s41467-022-31194-7

46. Bredow M, Monaghan J. Cross-kingdom regulation of calcium- and/or calmodulin-dependent protein kinases by phospho-switches that relieve autoinhibition. Current Opinion in Plant Biology. 2022;68:102251. 10.1016/j.pbi.2022.102251

47. Liu J, Zhang J, Wei Y, Su W, Li W, Wang B, et al. The nematode effector calreticulin competes with the high mobility group protein OsHMGB1 for binding to the rice calmodulin-like protein OsCML31 to enhance rice susceptibility to Meloidogyne graminicola. Plant, Cell & Environment. 2024;47:1732–46. 10.1111/pce.14848

48. Alvarez-Ponce D, Torres-Sánchez M, Feyertag F, Kulkarni A, Nappi T. Molecular evolution of DNMT1 in vertebrates: Duplications in marsupials followed by positive selection. PLOS ONE. Public Library of Science; 2018;13:e0195162. 10.1371/journal.pone.0195162

49. Gibbons RJ, McDowell TL, Raman S, O’Rourke DM, Garrick D, Ayyub H, et al. Mutations in ATRX, encoding a SWI/SNF-like protein, cause diverse changes in the pattern of DNA methylation. Nat Genet. Nature Publishing Group; 2000;24:368–71. 10.1038/74191

50. Voon HPJ, Hughes JR, Rode C, De La Rosa-Velázquez IA, Jenuwein T, Feil R, et al. ATRX Plays a Key Role in Maintaining Silencing at Interstitial Heterochromatic Loci and Imprinted Genes. Cell Reports. 2015;11:405–18. 10.1016/j.celrep.2015.03.036

51. Babikir H, Wang L, Shamardani K, Catalan F, Sudhir S, Aghi MK, et al. ATRX regulates glial identity and the tumor microenvironment in IDH-mutant glioma. Genome Biol. 2021;22:311. 10.1186/s13059-021-02535-4

52. Kernohan KD, Vernimmen D, Gloor GB, Bérubé NG. Analysis of neonatal brain lacking ATRX or MeCP2 reveals changes in nucleosome density, CTCF binding and chromatin looping. Nucleic Acids Res. 2014;42:8356–68. 10.1093/nar/gku564

53. Maurano MT, Wang H, John S, Shafer A, Canfield T, Lee K, et al. Role of DNA Methylation in Modulating Transcription Factor Occupancy. Cell Reports. 2015;12:1184–95. 10.1016/j.celrep.2015.07.024

54. Roseman SA, Siegenfeld AP, Lee C, Lue NZ, Waterbury AL, Liau BB. DNA methylation insulates genic regions from CTCF loops near nuclear speckles. Dalal Y, editor. eLife. eLife Sciences Publications, Ltd; 2025;13:RP102930. 10.7554/eLife.102930

55. Villate L, Esmenjaud D, Coedel S, Plantard O. Development of nine polymorphic microsatellite markers for the phytoparasitic nematode Xiphinema index, the vector of the grapevine fanleaf virus. Molecular Ecology Resources. 2009;9:229–32. 10.1111/j.1755-0998.2008.02411.x

56. Van Ghelder C, Reid A, Kenyon D, Esmenjaud D. Detection of Nepovirus Vector and Nonvector Xiphinema Species in Grapevine. In: Lacomme C, editor. Plant Pathology: Techniques and Protocols [Internet]. New York, NY: Springer; 2015 [cited 2026 Jan 21]. p. 149–59. 10.1007/978-1-4939-2620-6_12

57. Hooper D, Southey J. Laboratory Methods for Work with Plant and Soil Nematodes. Ministry of Agriculture, Fisheries and Food; 1986. p. 30.

58. Marçais G, Kingsford C. A fast, lock-free approach for efficient parallel counting of occurrences of k-mers. Bioinformatics. 2011;27:764–70. 10.1093/bioinformatics/btr011

59. Ranallo-Benavidez TR, Jaron KS, Schatz MC. GenomeScope 2.0 and Smudgeplot for reference-free profiling of polyploid genomes. Nat Commun. Nature Publishing Group; 2020;11:1–10. 10.1038/s41467-020-14998-3

60. De Coster W, D’Hert S, Schultz DT, Cruts M, Van Broeckhoven C. NanoPack: visualizing and processing long-read sequencing data. Bioinformatics. 2018;34:2666–9. 10.1093/bioinformatics/bty149

61. Cheng H, Concepcion GT, Feng X, Zhang H, Li H. Haplotype-resolved de novo assembly using phased assembly graphs with hifiasm. Nature Methods. Nature Publishing Group; 2021;18:170–5. 10.1038/s41592-020-01056-5

62. Brown M, González De la Rosa PM, Mark B. A Telomere Identification Toolkit [Internet]. Zenodo; 2023 [cited 2024 Oct 24]. 10.5281/zenodo.10091385

63. Dudchenko O, Batra SS, Omer AD, Nyquist SK, Hoeger M, Durand NC, et al. De novo assembly of the Aedes aegypti genome using Hi-C yields chromosome-length scaffolds. Science. American Association for the Advancement of Science; 2017;356:92–5. 10.1126/science.aal3327

64. Xu M, Guo L, Gu S, Wang O, Zhang R, Peters BA, et al. TGS-GapCloser: A fast and accurate gap closer for large genomes with low coverage of error-prone long reads. GigaScience. 2020;9:giaa094. 10.1093/gigascience/giaa094

65. Li H. Minimap2: pairwise alignment for nucleotide sequences. Bioinformatics. 2018;34:3094–100. 10.1093/bioinformatics/bty191

66. Quinlan AR, Hall IM. BEDTools: a flexible suite of utilities for comparing genomic features. Bioinformatics. 2010;26:841–2. 10.1093/bioinformatics/btq033

67. Kumar S, Jones M, Koutsovoulos G, Clarke M, Blaxter M. Blobology: exploring raw genome data for contaminants, symbionts and parasites using taxon-annotated GC-coverage plots. Front Genet [Internet]. 2013 [cited 2019 Mar 19];4. 10.3389/fgene.2013.00237

68. Laetsch DR, Blaxter ML. BlobTools: Interrogation of genome assemblies. F1000Research. 2017;6:1287. 10.12688/f1000research.12232.1

69. Sallet E, Gouzy J, Schiex T. EuGene: An Automated Integrative Gene Finder for Eukaryotes and Prokaryotes. Methods Mol Biol. 2019;1962:97–120. 10.1007/978-1-4939-9173-0_6

70. Howe KL, Bolt BJ, Shafie M, Kersey P, Berriman M. WormBase ParaSite - a comprehensive resource for helminth genomics. Mol Biochem Parasitol. 2017;215:2–10. 10.1016/j.molbiopara.2016.11.005

71. UniProt Consortium T. UniProt: the universal protein knowledgebase. Nucleic Acids Res. 2018;46:2699–2699. 10.1093/nar/gky092

72. Bao W, Kojima KK, Kohany O. Repbase Update, a database of repetitive elements in eukaryotic genomes. Mobile DNA. 2015;6:11. 10.1186/s13100-015-0041-9

73. Haas BJ, Papanicolaou A, Yassour M, Grabherr M, Blood PD, Bowden J, et al. De novo transcript sequence reconstruction from RNA-seq using the Trinity platform for reference generation and analysis. Nat Protocols. 2013;8:1494–512. 10.1038/nprot.2013.084

74. Mitchell A, Chang H-Y, Daugherty L, Fraser M, Hunter S, Lopez R, et al. The InterPro protein families database: the classification resource after 15 years. Nucl Acids Res. 2015;43:D213–21. 10.1093/nar/gku1243

75. Manni M, Berkeley MR, Seppey M, Simão FA, Zdobnov EM. BUSCO update: novel and streamlined workflows along with broader and deeper phylogenetic coverage for scoring of eukaryotic, prokaryotic, and viral genomes. Molecular Biology and Evolution [Internet]. 2021 [cited 2021 Aug 2]; 10.1093/molbev/msab199

76. Mitreva M, Jasmer DP, Zarlenga DS, Wang Z, Abubucker S, Martin J, et al. The draft genome of the parasitic nematode Trichinella spiralis. Nat Genet. 2011;43:228–35. 10.1038/ng.769

77. Korhonen PK, Pozio E, Rosa GL, Chang BCH, Koehler AV, Hoberg EP, et al. Phylogenomic and biogeographic reconstruction of the Trichinella complex. Nat Commun. Nature Publishing Group; 2016;7:1–8. 10.1038/ncomms10513

78. Foth BJ, Tsai IJ, Reid AJ, Bancroft AJ, Nichol S, Tracey A, et al. Whipworm genome and dual-species transcriptome analyses provide molecular insights into an intimate host-parasite interaction. Nat Genet. Nature Publishing Group; 2014;46:693–700. 10.1038/ng.3010

79. Lee Y-C, Ke H-M, Liu Y-C, Lee H-H, Wang M-C, Tseng Y-C, et al. Single-worm long-read sequencing reveals genome diversity in free-living nematodes. Nucleic Acids Research. 2023;51:8035–47. 10.1093/nar/gkad647

80. Emms DM, Kelly S. OrthoFinder: phylogenetic orthology inference for comparative genomics. Genome Biology. 2019;20:238. 10.1186/s13059-019-1832-y

81. Dobin A, Davis CA, Schlesinger F, Drenkow J, Zaleski C, Jha S, et al. STAR: ultrafast universal RNA-seq aligner. Bioinformatics. 2013;29:15–21. 10.1093/bioinformatics/bts635

82. Li B, Dewey CN. RSEM: accurate transcript quantification from RNA-Seq data with or without a reference genome. BMC Bioinformatics. 2011;12:323. 10.1186/1471-2105-12-323

83. Solovyev V. Statistical Approaches in Eukaryotic Gene Prediction. Handbook of Statistical Genetics [Internet]. John Wiley & Sons, Ltd; 2004 [cited 2025 Apr 2]. 10.1002/0470022620.bbc06

84. Buchfink B, Reuter K, Drost H-G. Sensitive protein alignments at tree-of-life scale using DIAMOND. Nat Methods. Nature Publishing Group; 2021;18:366–8. 10.1038/s41592-021-01101-x

85. Koutsovoulos GD, Noriot SG, Bailly-Bechet M, Danchin EGJ, Rancurel C. AvP: A software package for automatic phylogenetic detection of candidate horizontal gene transfers. PLOS Computational Biology. Public Library of Science; 2022;18:e1010686. 10.1371/journal.pcbi.1010686

86. Katoh K, Standley DM. MAFFT multiple sequence alignment software version 7: improvements in performance and usability. Mol Biol Evol. 2013;30:772–80. 10.1093/molbev/mst010

87. Price MN, Dehal PS, Arkin AP. FastTree 2 – Approximately Maximum-Likelihood Trees for Large Alignments. PLOS ONE. 2010;5:e9490. 10.1371/journal.pone.0009490

88. Shimodaira H, Hasegawa M. CONSEL: for assessing the confidence of phylogenetic tree selection. Bioinformatics (Oxford, England). 2001;17:1246–7.

89. Li W, Godzik A. Cd-hit: a fast program for clustering and comparing large sets of protein or nucleotide sequences. Bioinformatics. 2006;22:1658–9. 10.1093/bioinformatics/btl158

90. Capella-Gutiérrez S, Silla-Martínez JM, Gabaldón T. trimAl: a tool for automated alignment trimming in large-scale phylogenetic analyses. Bioinformatics. 2009;25:1972–3. 10.1093/bioinformatics/btp348

91. Shen W, Sipos B, Zhao L. SeqKit2: A Swiss army knife for sequence and alignment processing. iMeta. 2024;3:e191. 10.1002/imt2.191

92. Kalyaanamoorthy S, Minh BQ, Wong TKF, von Haeseler A, Jermiin LS. ModelFinder: fast model selection for accurate phylogenetic estimates. Nat Methods. Nature Publishing Group; 2017;14:587–9. 10.1038/nmeth.4285

93. Guindon S, Dufayard J-F, Lefort V, Anisimova M, Hordijk W, Gascuel O. New Algorithms and Methods to Estimate Maximum-Likelihood Phylogenies: Assessing the Performance of PhyML 3.0. Syst Biol. 2010;59:307–21. 10.1093/sysbio/syq010

94. Hoang DT, Chernomor O, von Haeseler A, Minh BQ, Vinh LS. UFBoot2: Improving the Ultrafast Bootstrap Approximation. Molecular Biology and Evolution. 2018;35:518–22. 10.1093/molbev/msx281

95. Letunic I, Bork P. Interactive Tree Of Life (iTOL) v4: recent updates and new developments. Nucleic Acids Res. Oxford Academic; 2019;47:W256–9. 10.1093/nar/gkz239

96. Shimodaira H. An Approximately Unbiased Test of Phylogenetic Tree Selection. Syst Biol. 2002;51:492–508. 10.1080/10635150290069913

