## Supplementary Material for "T2T genome of the basal nematode *Xiphinema index* reveals major adaptive events associated with plant parasitism and epigenetic regulation"

| | Assembly |
|---|---|
| **Contigs**<br><br>**(largest Mb)** | 73<br><br>(27.75) |
| **Size (Mb)** | 225.83 |
| **N50 (Mb)** | 23.66 |
| **N90 (Mb)** | 16.34 |
| **L50** | 5 |
| **L90** | 10 |
| **GC%** | 39.82 |
| **Total number of genes** | 33,647 |
| **Number of protein coding genes** | 30,614 |
| **Number of TE assigned to TE families** | 45,590 |
| **Number of genes completely included in family-assigned TE** | 1282 |

**Table S1:** Assembly metrics related to the *Xiphinema index* genome. Transposable element (TE) analysis was made using Earl Grey[1] version 4.2.4 with the options -m and "RepeatMasker search term" (-r).

| Species name | Short name | Clade | B Clade | (TTAGGC) n | Telomerase |
|---|---|---|---|---|---|
| *Xiphinema index* | Xi.inde | Longidoridae | Clade I | Y | Y |
| *Mesodorylaimus sp. YZB2_4* | Me.yzb2 | Mesodorylaimus | Clade I | Y | Y |
| *Trichinella nelsoni* | Tr.nels | Trichinellidae | Clade I | N | N |
| *Trichinella pseudospiralis* | Tr.pseu | Trichinellidae | Clade I | N | N |
| *Trichinella spiralis* | Tr.spir | Trichinellidae | Clade I | N | N |
| *Trichuris muris* | Tr.muri | Trichuridae | Clade I | Y | N |
| *Trichuris trichiura* | Tr.tric | Trichuridae | Clade I | Y | Y |
| *Ascaris suum* | As.suum | Ascaridomorpha | Clade III | Y | Y |
| *Parascaris univalens* | Pa.univ | Ascaridomorpha | Clade III | Y | Y |
| *Toxocara canis* | To.cani | Ascaridomorpha | Clade III | Y | Y |
| *Dracunculus medinensis* | Dr.medi | Dracunculidae | Clade III | Y | Y |
| *Brugia malayi* | Br.mala | Spiruromorpha | Clade III | Y | Y |
| *Dirofilaria immitis* | Di.immi | Spiruromorpha | Clade III | Y | Y |
| *Loa loa* | Lo.loa | Spiruromorpha | Clade III | Y | Y |
| *Onchocerca flexuosa* | On.flex | Spiruromorpha | Clade III | ? | Y |
| *Onchocerca volvulus* | On.volv | Spiruromorpha | Clade III | Y | Y |
| *Setaria digitata* | Se.digi | Spiruromorpha | Clade III | N | Y |
| *Rhabditophanes sp. KR3021* | Rh.spKR | Alloionematidae | Clade IV | N | N |
| *Ditylenchus destructor* | Di.dest | Anguinidae | Clade IV | Y | Y |
| *Ditylenchus dipsaci* | Di.dips | Anguinidae | Clade IV | Y | Y |
| *Aphelenchus avenae* | Ap.aven | Aphelenchidae | Clade IV | Y | Y |
| *Aphelenchoides besseyi* | Ap.bess | Aphelenchoididae | Clade IV | N | Y |
| *Aphelenchoides bicaudatus* | Ap.bica | Aphelenchoididae | Clade IV | N | N |
| *Aphelenchoides fujianensis* | Ap.fuji | Aphelenchoididae | Clade IV | Y | Y |
| *Bursaphelenchus okinawaensis* | Bu.okin | Aphelenchoididae | Clade IV | Y | ? |
| *Bursaphelenchus xylophilus* | Bu.xylo | Aphelenchoididae | Clade IV | Y | Y |
| *Globodera pallida* | Gl.pall | Hoplolaimidae | Clade IV | Y | Y |
| *Globodera rostochiensis* | Gl.rost | Hoplolaimidae | Clade IV | Y | Y |
| *Heterodera glycines* | He.glyc | Hoplolaimidae | Clade IV | Y | Y |
| *Heterodera schachtii* | He.scha | Hoplolaimidae | Clade IV | Y | Y |
| *Meloidogyne arenaria* | Me.aren | Meloidogynidae | Clade IV | N | N |
| *Meloidogyne chitwoodi-braker* | Me.chit.b | Meloidogynidae | Clade IV | N | N |
| *Meloidogyne graminicola* | Me.gram | Meloidogynidae | Clade IV | N | N |
| *Meloidogyne hapla* | Me.hapl | Meloidogynidae | Clade IV | N | N |
| *Meloidogyne incognita* | Me.inco | Meloidogynidae | Clade IV | N | N |
| *Meloidogyne javanica* | Me.java | Meloidogynidae | Clade IV | N | N |
| *Meloidogyne luci* | Me.luci | Meloidogynidae | Clade IV | N | N |
| *Meloidogyne enterolobii* | Me.ent | Meloidogynidae | Clade IV | N | N |
| *Halicephalobus mephisto* | Ha.meph | Panagrolaimidae | Clade IV | N | Y |
| *Panagrellus redivivus* | Pa.redi | Panagrolaimidae | Clade IV | N | Y |
| *Pratylenchus penetrans* | Pr.pene | Pratylenchidae | Clade IV | Y | Y |
| *Steinernema carpocapsae* | St.carp | Steinernematidae | Clade IV | Y | Y |
| *Parastrongyloides trichosuri* | Pa.tric | Strongyloididae | Clade IV | N | N |

| | | | | | |
|---|---|---|---|---|---|
| *Strongyloides ratti* | St.ratt | Strongyloididae | Clade IV | N | N |
| *Strongyloides stercoralis* | St.ster | Strongyloididae | Clade IV | N | N |
| *Strongyloides venezuelensis* | St.vene | Strongyloididae | Clade IV | N | N |
| *Caenorhabditis briggsae* | Ca.brig | Anarhabditis | Clade V | Y | Y |
| *Caenorhabditis elegans* | Ca.eleg | Anarhabditis | Clade V | Y | Y |
| *Caenorhabditis remanei* | Ca.rema | Anarhabditis | Clade V | Y | Y |
| *Diploscapter coronatus* | Di.coro | Anarhabditis | Clade V | N | Y |
| *Diploscapter pachys* | Di.pach | Anarhabditis | Clade V | N | Y |
| *Allodiplogaster sudhausi* | Al.sudh | Diplogastridae | Clade V | Y | ? |
| *Micoletzkya japonica* | Mi.japo | Diplogastridae | Clade V | Y | N |
| *Parapristionchus giblindavisi* | Pa.gibl | Diplogastridae | Clade V | Y | N |
| *Pristionchus exspectatus* | Pr.exsp | Diplogastridae | Clade V | Y | ? |
| *Pristionchus fissidentatus* | Pr.fiss | Diplogastridae | Clade V | Y | N |
| *Pristionchus pacificus* | Pr.paci | Diplogastridae | Clade V | Y | N |
| *Heterorhabditis bacteriophora* | He.bact | Heterorhabditidae | Clade V | ? | ? |
| *Mesorhabditis belari* | Me.bela | Pleiorhabditis | Clade V | Y | ? |
| *Ancylostoma caninum* | An.cani | Strongylida | Clade V | Y | Y |
| *Angiostrongylus cantonensis* | An.cant | Strongylida | Clade V | Y | ? |
| *Ancylostoma ceylanicum* | An.ceyl | Strongylida | Clade V | Y | Y |
| *Angiostrongylus costaricensis* | An.cost | Strongylida | Clade V | Y | ? |
| *Angiostrongylus vasorum* | An.vaso | Strongylida | Clade V | ? | ? |
| *Dictyocaulus viviparus* | Di.vivi | Strongylida | Clade V | Y | N |
| *Haemonchus contortus* | Ha.cont | Strongylida | Clade V | Y | Y |
| *Heligmosomoides polygyrus* | He.poly | Strongylida | Clade V | Y | Y |
| *Necator americanus* | Ne.amer | Strongylida | Clade V | ? | ? |
| *Parelaphostrongylus tenuis* | Pa.tenu | Strongylida | Clade V | Y | Y |
| *Auanema sp. JU1783* | Au.spJU | Synrhabditis | Clade V | Y | Y |
| *Oscheius tipulae* | Os.tipu | Synrhabditis | Clade V | Y | Y |
| *Hypsibius exemplaris* | Hy.exem | Tardigrada | Outgroup | ? | Y |

**Table S2:** List of 71 nematode species and the tardigrada species used in the orthofinder analysis showing the presence/absence of the canonical telomeric repeat (TTAGGC)n and the telomerase candidate (Yes/No).

| BUSCO met n=954 | %Compl.[Single, Duplic.] | %Frag. | %Miss. | Geno. size / Prot. # |
|---|---|---|---|---|
| ***Xiphinema index* (genome)** | C:84.5 [S:78.7,D:5.8] | F:2.0 | M:13.5 | 226 Mb |
| **(predicted proteins)** | C:86.9 [S:81.8,D:5.1] | F:3.5 | M:9.6 | 30,614 |
| ***Mesodorylaimus sp. YZB2_4*** | C:86.5 [S:84.3,D:2.2] | F:1.8 | M:11.7 | 142 Mb |
| | C:91.0 [S:88.7,D:2.3] | F:1.9 | M:7.1 | 22,630 |
| ***Trichinella spiralis*** | C:74.7 [S:69.7,D:5.0] | F:2.4 | M:22.9 | 63 Mb |
| | C:62.4 [S:59.4;D:3.0] | F:6.3 | M:31.3 | 16,380 |
| ***Trichinella nelsoni*** | C:75.0 [S:74.0,D:1.0] | F:2.1 | M:22.9 | 47 Mb |
| | C:73.3 [S:47.5,D:25.8] | F:2.5 | M:24.2 | 13,229 |
| ***Trichuris muris*** | C:71.0 [S:59.5,D:11.5] | F:2.1 | M:26.9 | 112 Mb |
| | C:77.1 [S:65.4,D:11.7] | F:3.0 | M:19.9 | 14,995 |
| ***Trichuris trichiura*** | C:71.8 [S:68.9,D:2.9] | F:2.8 | M:25.4 | 81 Mb |
| | C:76.8 [S:69.1,D:7.7] | F:3.0 | M:20.2 | 12,671 |

**Table S3:** BUSCO v5.3.1[2] metazoa (odb10) completeness assessment of *X. index* with five other Dorylaimia. First line at the genome level, second line at the protein level.

| HGT event | Sequence name | Origin of donor sequences | Alternative topology | Similarities with the donor sequences | | Length and coverage of the alignment with the donor sequences | | Local score for genomic environment | Representative InterPro annotation | | CAZyme | HGT candidate in[3] |
|---|---|---|---|---|---|---|---|---|---|---|---|---|
| | | | | max id | average id | average aln length | average coverage | | | | | |
| XindB01 | XINDMERG_xiphind_chr_04g018470 | Bacteria | / | 52.0 | 48.0 | 434 | 92.7 | 0.56 | IPR000801 | Esterase-like | | Yes |
| XindB02 | XINDMERG_xiphind_chr_01g006360 | Bacteria | 0 | 52.8 | 51.0 | 453 | 96.0 | 0.4 | IPR029058 | Alpha/Beta hydrolase fold | | |
| | XINDMERG_xiphind_chr_09g014320 | | | 47.6 | 46.3 | 645 | 94.4 | 0.68 | | | | |
| | XINDMERG_xiphind_chr_09g014080 | | | 54.2 | 51.5 | 332 | 92.0 | 0.59 | | | | |
| | XINDMERG_xiphind_chr_01g006350 | | | 50.9 | 47.3 | 162 | 78.7 | 0.4 | | | | |
| XindB03 | XINDMERG_xiphind_chr_05g013770 | Bacteria | / | 38.4 | 35.3 | 487 | 88.9 | 0.36 | IPR050416 | FAD-linked Oxidoreductases in Biosynthetic Pathways | | Yes |
| | XINDMERG_xiphind_chr_05g007790 | | | 38.1 | 35.6 | 470 | 84.8 | 0.6 | | | | |
| | XINDMERG_xiphind_chr_10g005810 | | | 41.0 | 36.4 | 262 | 62.4 | 0.45 | | | | |
| XindB04 | XINDMERG_xiphind_chr_02g031130 | Bacteria | / | 47.2 | 43.9 | 245 | 86.0 | 0.45 | IPR051206 | N-acetylmuramoyl-L-alanine amidase 2 | | Yes |
| | XINDMERG_xiphind_chr_02g031100 | | | 46.5 | 42.6 | 249 | 88.2 | 0.53 | | | | |
| | XINDMERG_xiphind_chr_04g020390 | | | 48.3 | 44.3 | 249 | 83.2 | 0.58 | | | | |
| XindB05 | XINDMERG_xiphind_chr_04g016490 | Bacteria | / | 66.7 | 62.2 | 508 | 93.1 | 0.52 | IPR000743 | Glycoside hydrolase, family 28 | GH28 | |
| XindB06 | XINDMERG_xiphind_chr_01g023500 | Bacteria | / | 36.2 | 33.8 | 374 | 92.3 | 0.35 | IPR019405 IPR050282 | Lactonase, 7-bladed beta propeller Cycloisomerase 2 | | Yes |
| | XINDMERG_xiphind_chr_01g022900 | | | 37.1 | 33.9 | 372 | 91.2 | 0.6 | | | | |
| XindB07 | XINDMERG_xiphind_chr_06g030680 | Bacteria | / | 49.0 | 46.7 | 502 | 95.1 | 0.72 | IPR001362 | Glycoside hydrolase, family 32 | GH32 | Yes |

| | | | | | | | | | | | | |
|---|---|---|---|---|---|---|---|---|---|---|---|---|
| XindB08 | XINDMERG_xiphind_chr_01g043960 | Bacteria | / | 61.0 | 56.5 | 188 | 90.0 | 0.72 | IPR014509 | Uncharacterised conserved protein UCP020606 | | Yes |
| XindB09 | XINDMERG_xiphind_chr_06g027900 | Bacteria | / | 49.3 | 44.8 | 237 | 77.0 | 0.56 | IPR050261 | FrsA esterase | | |
| | XINDMERG_xiphind_chr_05g011870 | | | 53.8 | 51.4 | 236 | 81.1 | 0.51 | | | | |
| | XINDMERG_xiphind_chr_01g013830 | | | 57.3 | 52.4 | 178 | 85.6 | 0.68 | | | | |
| XindB10 | XINDMERG_xiphind_chr_09g012500 | Bacteria | / | 40.7 | 37.0 | 189 | 62.1 | 0.56 | IPR028978 | Chorismate pyruvate-lyase/UbiC transcription regulator-associated domain superfamily | | Yes |
| | XINDMERG_xiphind_chr_09g005200 | | | 48.1 | 44.4 | 189 | 82.5 | 0.64 | | | | |
| XindB11 | XINDMERG_xiphind_chr_03g025030 | Bacteria | 0 | 39.2 | 36.4 | 181 | 62.3 | 0.72 | IPR007065 | HPP | | |
| XindB12 | XINDMERG_xiphind_chr_04g010420 | Bacteria | / | 47.3 | 41.7 | 228 | 88.0 | 0.42 | IPR002594 | Glycoside hydrolase family 12 | GH12 | Yes |
| | XINDMERG_xiphind_chr_04g010460 | | | 49.1 | 43.1 | 217 | 78.1 | 0.5 | | | GH12 | |
| | XINDMERG_xiphind_chr_01g044370 | | | 52.6 | 50.2 | 211 | 75.3 | 0.68 | | | GH12 | |
| XindB13 | XINDMERG_xiphind_chr_07g002070 | Bacteria | / | 31.6 | 28.8 | 199 | 63.2 | 0.56 | IPR013078 | Histidine phosphatase superfamily, clade-1 | | |
| XindB14 | XINDMERG_xiphind_chr_06g001180 | Bacteria | / | 67.2 | 65.7 | 261 | 96.0 | 0.44 | IPR003329 | Acylneuraminate cytidylyltransferase | | |
| XindB15 | XINDMERG_xiphind_chr_01g019320 | Bacteria | / | 52.5 | 49.4 | 189 | 69.2 | 0.68 | IPR028978 | Chorismate pyruvate-lyase/UbiC transcription regulator-associated domain superfamily | | |
| XindB16 | XINDMERG_xiphind_chr_05g021260 | Bacteria | / | 31.9 | 29.4 | 370 | 93.8 | 0.68 | IPR019405<br>IPR050282 | Lactonase, 7-bladed beta-propeller<br>Cycloisomerase 2 | | |
| XindB17 | XINDMERG_xiphind_chr_09g007140 | Bacteria | / | 68.7 | 67.8 | 805 | 99.6 | 0.43 | IPR017853 | Glycoside hydrolase superfamily | GH31_2 | Yes |

| | | | | | | | | | | | | |
|---|---|---|---|---|---|---|---|---|---|---|---|---|
| XindB18 | XINDMERG_xiphind_chr_02g023510 | Bacteria | 0 | 50.0 | 47.1 | 239 | 78.2 | 0.68 | IPR007325 | Kynurenine formamidase/cyclase-like | | |
| XindB19 | XINDMERG_xiphind_chr_10g004810 | Bacteria | / | 56.9 | 56.2 | 283 | 99.6 | 0.36 | IPR003721 | Pantoate-beta-alanine ligase | | |
| XindB20 | XINDMERG_xiphind_chr_01g037860 | Bacteria | 1 | 66.1 | 57.2 | 273 | 90.0 | 0.63 | IPR017853 | Glycoside hydrolase superfamily | | |
| XindB21 | XINDMERG_xiphind_chr_04g027920 | Bacteria | / | 32.1 | 29.6 | 168 | 49.9 | 0.72 | IPR039275 | PDZ domain-containing protein 8 | | |
| XindB22 | XINDMERG_xiphind_chr_05g001500 | Bacteria | / | 54.7 | 52.4 | 500 | 88.6 | 0.64 | IPR045851 | AMP-binding enzyme, C-terminal domain superfamily | | |
| XindB23 | XINDMERG_xiphind_chr_07g014580 | Bacteria | / | 51.9 | 50.5 | 510 | 88.7 | 0.64 | IPR045851 | AMP-binding enzyme, C-terminal domain superfamily | | |
| XindF01 | XINDMERG_xiphind_chr_01g012830 | Fungi | 0 | 68.4 | 64.5 | 260 | 63.9 | 0.8 | IPR001905 | Ammonium transporter | | Yes |
| | XINDMERG_xiphind_chr_04g018740 | Fungi | | 66.1 | 64.1 | 425 | 84.3 | 0.52 | | | | |
| | XINDMERG_xiphind_chr_10g014920 | Fungi | | 58.3 | 56.9 | 415 | 90.7 | 0.52 | | | | |
| XindF02 | XINDMERG_xiphind_chr_03g002360 | Fungi | / | 33.8 | 30.7 | 267 | 93.1 | 0.72 | IPR000490 | Glycoside hydrolase family 17 | | |
| XindF03 | XINDMERG_xiphind_chr_04g013470 | Fungi | / | 57.4 | 55.4 | 789 | 99.5 | 0.23 | IPR006276 | Cobalamin-independent methionine synthase | | Yes |
| XindF04 | XINDMERG_xiphind_chr_09g000010 | Fungi | / | 54.8 | 46.9 | 1924 | 97.8 | 0.49 | IPR014084 | Urea carboxylase | | |
| XindF05 | XINDMERG_xiphind_chr_02g036040 | Fungi | ? | 68.5 | 61.3 | 148 | 82.9 | 0.72 | IPR014980 | Dopa 4,5-dioxygenase | | Yes |
| XindV01 | XINDMERG_xiphind_chr_06g025050 | Viridiplantae | / | 39.2 | 35.7 | 144 | 62.2 | 0.76 | IPR011992 | EF-hand domain pair | | |

**Table S4:** Validated HGT candidates from potential bacterial, fungal or *Viridiplantae* donors in the *Xiphinema index* genome

| *Species names* | *Short name* | DNMT1 | DNMT2 | DNMT3 | Full ATRX |
|---|---|---|---|---|---|
| *Aetholaimus rotundicauda* | *Ae.rotu* | 0.5 | 1 | 1 | 1 |
| *Allodiplogaster sudhausi* | *Al.sudh* | 0 | 1 | 0 | 0 |
| *Ancylostoma caninum* | *An.cani* | 0 | 0 | 0 | 0 |
| *Ancylostoma ceylanicum* | *An.ceyl* | 0 | 0 | 0 | 0 |
| *Angiostrongylus cantonensis* | *An.cant* | 0 | 0 | 0 | 0 |
| *Angiostrongylus costaricensis* | *An.cost* | 0 | 0 | 0 | 0 |
| *Angiostrongylus vasorum* | *An.vaso* | 0 | 0 | 0 | 0 |
| *Anticoma sp* | *An.sp* | 0 | 0 | 0 | 0 |
| *Aphelenchoides besseyi* | *Ap.bess* | 0 | 0 | 0 | 0 |
| *Aphelenchoides bicaudatus* | *Ap.bica* | 0 | 0 | 0 | 0 |
| *Aphelenchoides fujianensis* | *Ap.fuji* | 0 | 0 | 0 | 0 |
| *Aphelenchus avenae* | *Ap.aven* | 0 | 0 | 0 | 0 |
| *Ascaris suum* | *As.suum* | 0 | 1 | 0 | 0 |
| *Auanema sp. JU1783* | *Au.spJU* | 0 | 0.5 | 0 | 0 |
| *Brugia malayi* | *Br.mala* | 0 | 1 | 0 | 0 |
| *Bursaphelenchus okinawaensis* | *Bu.okin* | 0 | 0 | 0 | 0 |
| *Bursaphelenchus xylophilus* | *Bu.xylo* | 0 | 0 | 0 | 0 |
| *Caenorhabditis briggsae* | *Ca.brig* | 0 | 0 | 0 | 0 |
| *Caenorhabditis elegans* | *Ca.eleg* | 0 | 0 | 0 | 0 |
| *Caenorhabditis remanei* | *Ca.rema* | 0 | 0 | 0 | 0 |
| *Coomansus parvus* | *Co.parv* | 0 | 0 | 1 | 1 |
| *Dictyocaulus viviparus* | *Di.vivi* | 0 | 0 | 0 | 0 |
| *Diphtherophora sp* | *Di.sp* | 0 | 1 | 0 | 0 |
| *Diploscapter coronatus* | *Di.coro* | 0 | 0 | 0 | 0 |
| *Diploscapter pachys* | *Di.pach* | 0 | 0 | 0 | 0 |
| *Dirofilaria immitis* | *Di.immi* | 0 | 1 | 0 | 0 |
| *Discolaimus major* | *Di.majo* | 0.5 | 1 | 1 | 1 |
| *Ditylenchus destructor* | *Di.dest* | 0 | 0 | 0 | 0 |
| *Ditylenchus dipsaci* | *Di.dips* | 0 | 0 | 0 | 0 |
| *Dorylaimoides thornei* | *Do.thor* | 1 | 1 | 0.5 | 1 |
| *Dracunculus medinensis* | *Dr.medi* | 0 | 0 | 0 | 0 |
| *Etamphidelus pieterloofi* | *Et.piet* | 0 | 0 | 0 | 1 |
| *Eudorylaimus piceae* | *Eu.pice* | 0.5 | 1 | 0.5 | 0 |
| *Globodera pallida* | *Gl.pall* | 0 | 0 | 0 | 0 |
| *Globodera rostochiensis* | *Gl.rost* | 0 | 0 | 0 | 0 |
| *Haemonchus contortus* | *Ha.cont* | 0 | 1 | 0 | 0 |

| | | | | | |
|---|---|---|---|---|---|
| *Halicephalobus mephisto* | *Ha.meph* | 0 | 0 | 0 | 0 |
| *Heligmosomoides polygyrus* | *He.poly* | 0 | 1 | 0 | 0 |
| *Heterodera glycines* | *He.glyc* | 0 | 0 | 0 | 0 |
| *Heterodera schachtii* | *He.scha* | 0 | 0 | 0 | 0 |
| *Heterorhabditis bacteriophora* | *He.bact* | 0 | 1 | 0 | 0 |
| *Hypsibius exemplaris* | *Hy.exem* | 0 | 1 | 0 | 0 |
| *Iotonchus arcuatus* | *Io.arcu* | 0 | 1 | 1 | 0 |
| *Ironus dentifurcatus* | *Ir.dent* | 0 | 0 | 0 | 0 |
| *Loa loa* | *Lo.loa* | 0 | 0 | 0 | 0 |
| *Longidorus juglans* | *Lo.jugl* | 0 | 1 | 0.5 | 1 |
| *Meloidogyne arenaria* | *Me.aren* | 0 | 0 | 0 | 0 |
| *Meloidogyne chitwoodi-braker* | *Me.chit* | 0 | 0 | 0 | 0 |
| *Meloidogyne enterolobii* | *Me.ent* | 0 | 0 | 0 | 0 |
| *Meloidogyne graminicola* | *Me.gram* | 0 | 0 | 0 | 0 |
| *Meloidogyne hapla* | *Me.hapl* | 0 | 0 | 0 | 0 |
| *Meloidogyne incognita* | *Me.inco* | 0 | 0 | 0 | 0 |
| *Meloidogyne javanica* | *Me.java* | 0 | 0 | 0 | 0 |
| *Meloidogyne luci* | *Me.luci* | 0 | 0 | 0 | 0 |
| *Mesodorylaimus sp. YZB2_4* | *Me.yzb2* | 1 | 1 | 1 | 1 |
| *Mesorhabditis belari* | *Me.bela* | 0 | 1 | 0 | 0 |
| *Micoletzkya japonica* | *Mi.japo* | 0 | 0.5 | 0 | 0 |
| *Miconchus hopperi* | *Mi.hopp* | 0 | 0 | 0 | 0 |
| *Mylonchulus hawaiiensis* | *My.hawa* | 0 | 0 | 0.5 | 1 |
| *Necator americanus* | *Ne.amer* | 0 | 1 | 0 | 0 |
| *Onchocerca flexuosa* | *On.flex* | 0 | 1 | 0 | 0 |
| *Onchocerca volvulus* | *On.volv* | 0 | 1 | 0 | 0 |
| *Oscheius tipulae* | *Os.tipu* | 0 | 1 | 0 | 0 |
| *Oxybelondira paraperplexa* | *Ox.para* | 0.5 | 0 | 1 | 1 |
| *Panagrellus redivivus* | *Pa.redi* | 0 | 0 | 0 | 0 |
| *Paractinolaimus indicus* | *Pa.indi* | 0.5 | 1 | 0.5 | 0 |
| *Parapristionchus giblindavisi* | *Pa.gibl* | 0 | 1 | 0 | 0 |
| *Parascaris univalens* | *Pa.univ* | 0 | 1 | 0 | 0 |
| *Parastrongyloides trichosuri* | *Pa.tric* | 0 | 0 | 0 | 0 |
| *Paravulvus hartingii* | *Pa.hart* | 0.5 | 0 | 0.5 | 0 |
| *Parelaphostrongylus tenuis* | *Pa.tenu* | 0 | 0 | 0 | 0 |
| *Pratylenchus penetrans* | *Pr.pene* | 0 | 0 | 0 | 0 |
| *Prionchulus muscorum* | *Pr.musc* | 0 | 1 | 1 | 0 |
| *Prismatolaimus dolichuru* | *Pr.doli* | 0 | 0 | 0 | 0 |
| *Pristionchus exspectatus* | *Pr.exsp* | 0 | 1 | 0 | 0 |
| *Pristionchus fissidentatus* | *Pr.fiss* | 0 | 1 | 0 | 0 |
| *Pristionchus pacificus* | *Pr.paci* | 0 | 0.5 | 0 | 0 |
| *Prodorylaimus andrassyi* | *Pr.andr* | 1 | 1 | 1 | 0 |

| | | | | | |
|---|---|---|---|---|---|
| *Rhabditophanes sp. KR3021* | *Rh.spKR* | 0 | 0 | 0 | 0 |
| *Sectonema barbatoides* | *Se.barb* | 1 | 0 | 0.5 | 0 |
| *Setaria digitata* | *Se.digi* | 0 | 1 | 0 | 0 |
| *Steinernema carpocapsae* | *St.carp* | 0 | 1 | 0 | 0 |
| *Strongyloides ratti* | *St.ratt* | 0 | 0 | 0 | 0 |
| *Strongyloides stercoralis* | *St.ster* | 0 | 0 | 0 | 0 |
| *Strongyloides venezuelensis* | *St.vene* | 0 | 0 | 0 | 0 |
| *Thornia sp* | *Th.sp* | 1 | 0 | 1 | 1 |
| *Toxocara canis* | *To.cani* | 0 | 1 | 0 | 0 |
| *Trachactinolaimus nanjingensis* | *Tr.nanj* | 0.5 | 0 | 0.5 | 0 |
| *Trichinella nelsoni* | *Tr.nels* | 0 | 1 | 0.5 | 0 |
| *Trichinella pseudospiralis* | *Tr.pseu* | 0 | 1 | 0.5 | 0 |
| *Trichinella spiralis* | *Tr.spir* | 0 | 0 | 0 | 0 |
| *Trichodorus pakistanensis* | *Tr.paki* | 0 | 1 | 0 | 0 |
| *Trichuris muris* | *Tr.muri* | 0 | 0.5 | 0 | 0 |
| *Trichuris trichiura* | *Tr.tric* | 0 | 0 | 0 | 0 |
| *Tripyla glomerans* | *Tr.glom* | 0 | 0 | 0 | 0 |
| *Tripylina zhejiangensis* | *Tr.zhej* | 0 | 1 | 0 | 0 |
| *Trischistoma pellucidum* | *Tr.pell* | 0 | 1 | 0 | 1 |
| *Tylencholaimus nagauriensis* | *Ty.naga* | 1 | 1 | 1 | 1 |
| *Xiphinema index* | *Xi.inde* | 1 | 1 | 1 | 1 |

**Table S5:** List of nematode species screened for presence/absence of DNMT1, 2, 3 and ATRX orthologs. 0=no evidence, 0.5=partial evidence, 1=presence of a full predicted protein.

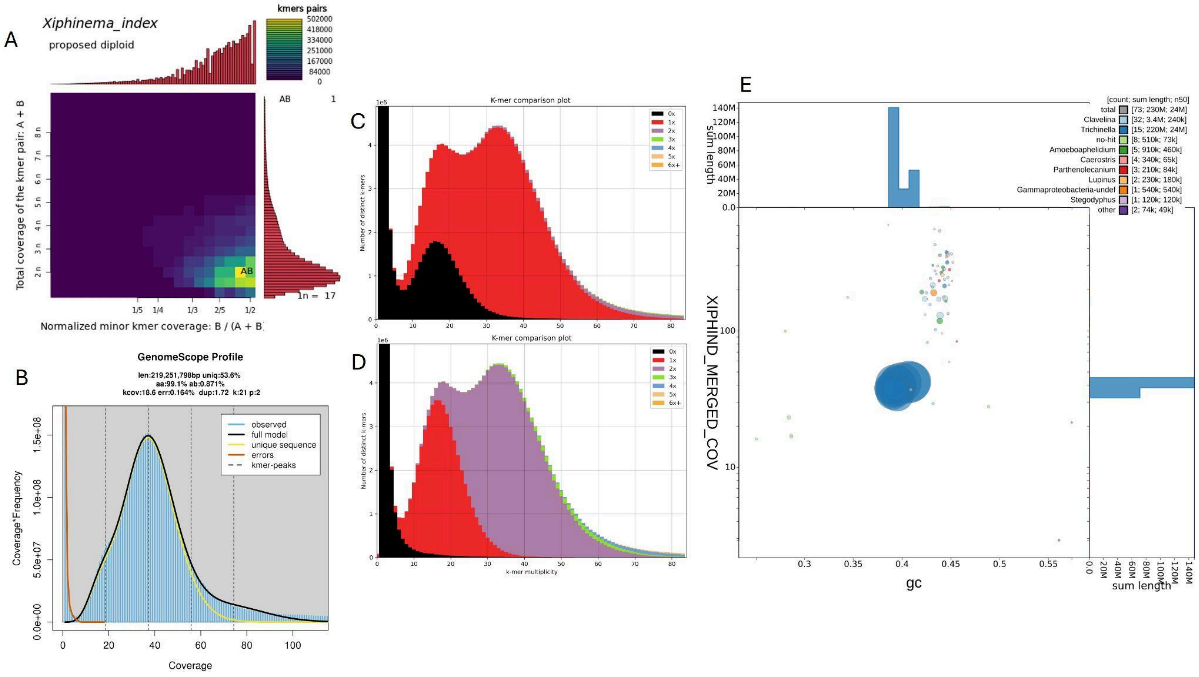


**Figure S1: Genome profiling of *Xiphinema index*.**

(A) Smudgeplot[4] of *X. index* extracting 21-mers from HiFi reads. The color intensity of each smudge reflects the approximate number of k-mers per bin. *X. index* is predicted to have a diploid genome. (B) GenomeScope2[4] k-mer profile and estimated parameters for *X. index*. Coverage (kcov), error rate (err.), haploid genome size estimation (len.), k-mer size (k) and ploidy level (p). *X. index* shows an average heterozygosity of ca. 0.9% with a predicted haploid genome size of 219 Mb. (C) Genome assembly vs. reads k-mer spectra. The Kat[5] spectrum plot using HiFi reads reports the multiplicity of each k-mer detected in the read set and in the assembly. The plot is color-coded according to the number of times a k-mer is found in the assembly. The plot is based on the merged consensus assembly. (D) The same plot but based on concatenation of haplotypes 1 and 2. (E) BlobPlot[6] of *X. index* genome assembly showing taxonomic affiliation at the genus rank level. Sequence length (sum) is mentioned above and on the right panel. Marginal contamination is detected in debris., which were eliminated from the final assembly.

A

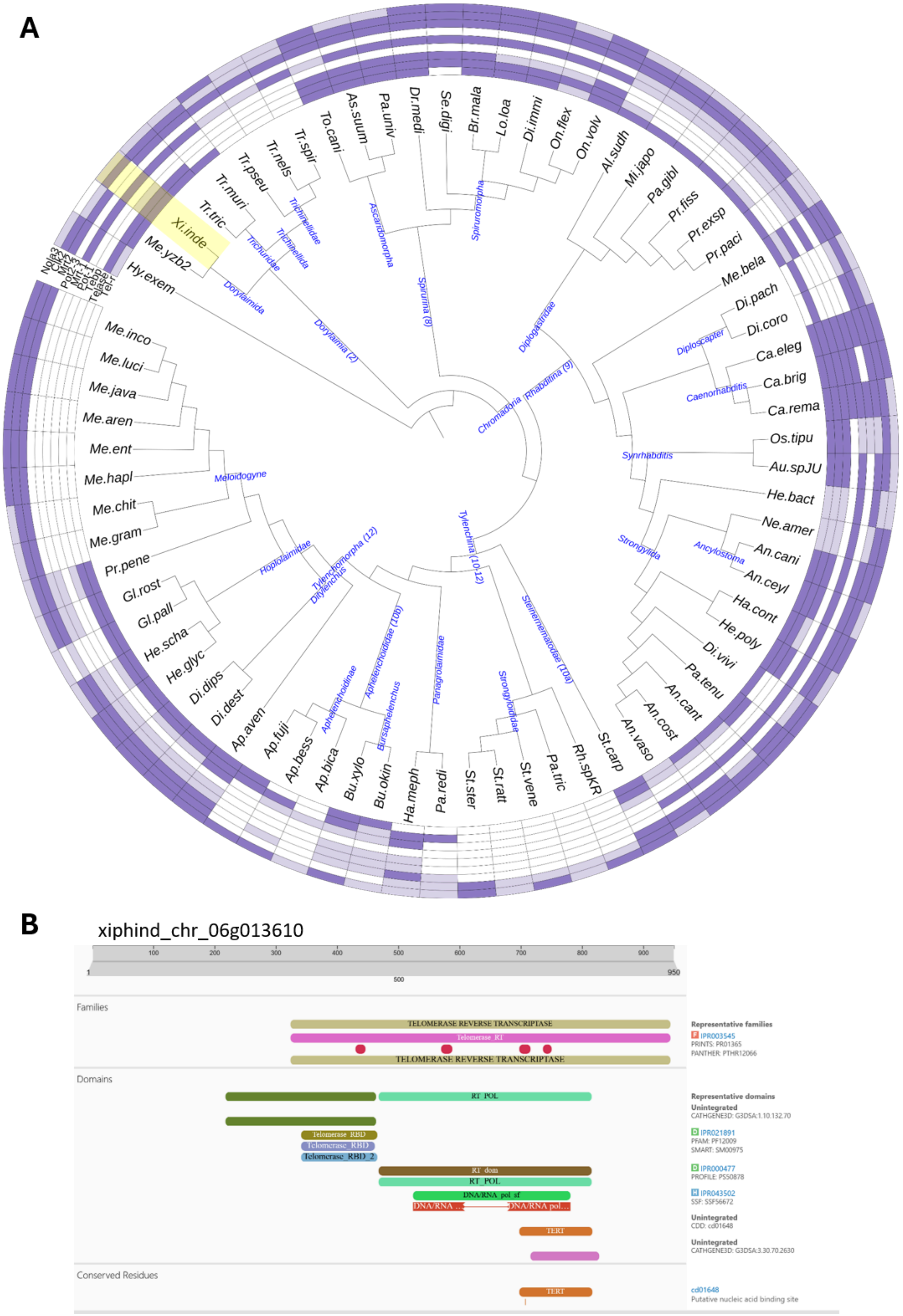
A
Xi.inde
Me.yzb2
Hy.exem
Tr.tric
Tr.muri
Tr.pseu
Tr.nels
Tr.spir
To.cani
As.suum
Pa.univ
Dr.medi
Se.digi
Br.mala
Lo.loa
Di.immi
On.flex
On.volv
Al.sudh
Mi.japo
Pa.gibl
Pr.fiss
Pr.exsp
Pr.paci
Me.bela
Di.pach
Di.coro
Ca.eleg
Ca.brig
Ca.rema
Os.tipu
Au.spJU
He.bact
Ne.amer
An.cani
An.ceyl
Ha.cont
He.poly
Di.vivi
Pa.tenu
An.cant
An.cost
An.vaso
St.carp
Rh.spKR
Pa.tric
St.vene
St.ratt
St.ster
Pa.redi
Ha.meph
Bu.okin
Bu.xylo
Ap.bica
Ap.bess
Ap.fuji
Ap.aven
Di.dest
Di.dips
He.glyc
He.scha
Gl.pall
Gl.rost
Pr.pene
Me.gram
Me.chit
Me.hapl
Me.ent
Me.aren
Me.java
Me.luci
Me.inco
Nola3
Clk2
Mrt-2
Pot-2
Pot-1
Tebp
Telase
Tel-I
Trichinellidae
Trichuridae
Trichinellida
Dorylaimida
Dorylaimia (2)
Ascaridomorpha
Spiruromorpha
Spirurina (8)
Diplogastridae
Rhabditina (9)
Chromadoria
Diploscapter
Caenorhabditis
Synrhabditis
Strongylida
Ancylostoma
Meloidogyne
Hoplolaimidae
Tylenchomorpha (12)
Ditylenchus
Tylenchina (10-12)
Aphelenchoidinae
Aphelenchoididae (10b)
Bursaphelenchus
Panagrolaimidae
Strongyloididae
Steinernematidae (10a)
B
xiphind_chr_06g013610
1
100
200
300
400
500
600
700
800
900
950
Families
TELOMERASE REVERSE TRANSCRIPTASE
Telomerase_RT
TELOMERASE REVERSE TRANSCRIPTASE
Representative families
IPR003545
PRINTS: PR01365
PANTHER: PTHR12066
Domains
RT_POL
Telomerase_RBD
Telomerase_RBD
Telomerase_RBD_2
RT_dom
RT_POL
DNA/RNA_pol_sf
DNA/RNA ...
DNA/RNA_pol...
TERT
Representative domains
Unintegrated
CATHGENE3D: G3DSA:1.10.132.70
IPR021891
PFAM: PF12009
SMART: SM00975
IPR000477
PROFILE: PS50878
IPR043502
SSF: SSF56672
Unintegrated
CDD: cd01648
Unintegrated
CATHGENE3D: G3DSA:3.30.70.2630
Conserved Residues
TERT
cd01648
Putative nucleic acid binding site

**Figure S2: Distribution of telomere repeats and telomere-associated proteins in the phylum Nematoda and functional annotation of the putative *X. index* telomerase enzyme.**
**(A)** Phylogenetic tree showing the presence/absence of canonical telomere repeats and telomere-associated proteins across nematode genomes. Violet color means present, light-violet means partial evidence and blank no evidence. Tel-r: *C. elegans* telomeric repeat (TTAGGC)n except for the outgroup *Hypsibus exemplaris* in which another simple 9-nucleotides repeat has been described[7]. Telase: telomerase reverse transcriptase. Tebp: double strand (ds)-telomeric DNA binding proteins Tebp1 or Tebp2. Pot1: single strand (ss)-telomeric DNA binding protein Pot-1. Mrt1: ss-telomeric DNA binding protein Mrt-1. Pot2-3: ss-telomeric DNA binding proteins Pot2 or Pot3. Mrt2 and Clk2: proteins with a putative function in telomere length regulation. Nola3: H/ACA ribonucleoprotein complex subunit 3. **(B)** Functional annotation of the putative *X. index* telomerase (xiphind_chr_06g013610) displaying the canonical domains found in the human telomerase (TERT-O14746). xiphind_chr_06g013610 shows 24.62% of sequence identity with TERT.

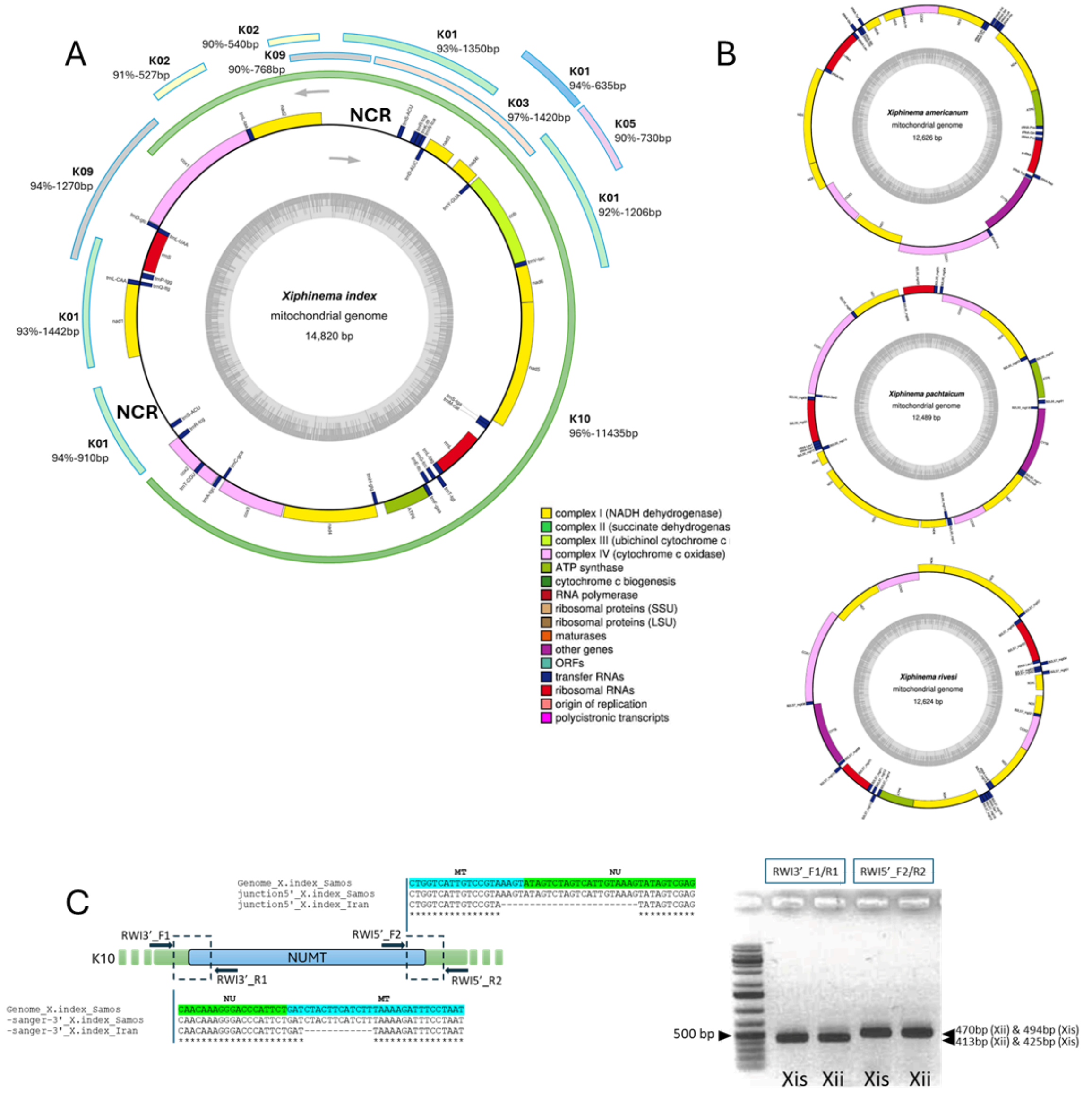


**Figure S3: Mitochondrial genome and nuclear mitochondrial DNA segments (NUMTs) of *Xiphinema index.***

A) mitochondrial genome of *X. index* with the NUMTs positioned on the mitogenome. Their chromosome locations (K1 to K10), lengths and percentages of identity with the corresponding sequences in the mitogenome are indicated. NCR for non coding region. B) Mitogenomes of *X. americanum* (NC_005928.1), *X. rivesi* (NC_033869.1), and *X. pachtaïcum* (NC_033870.1). C) PCR validation of the main NUMT on the chromosome 10 (K10). Two distinct populations of *X. index* were PCR-tested with 5' and 3' flanking primers (RW5' and RW3') and sanger sequenced: *X. index* 'samos' (Xis) corresponding to the genome (Xis) and *X. index* 'Iran' (Xii).

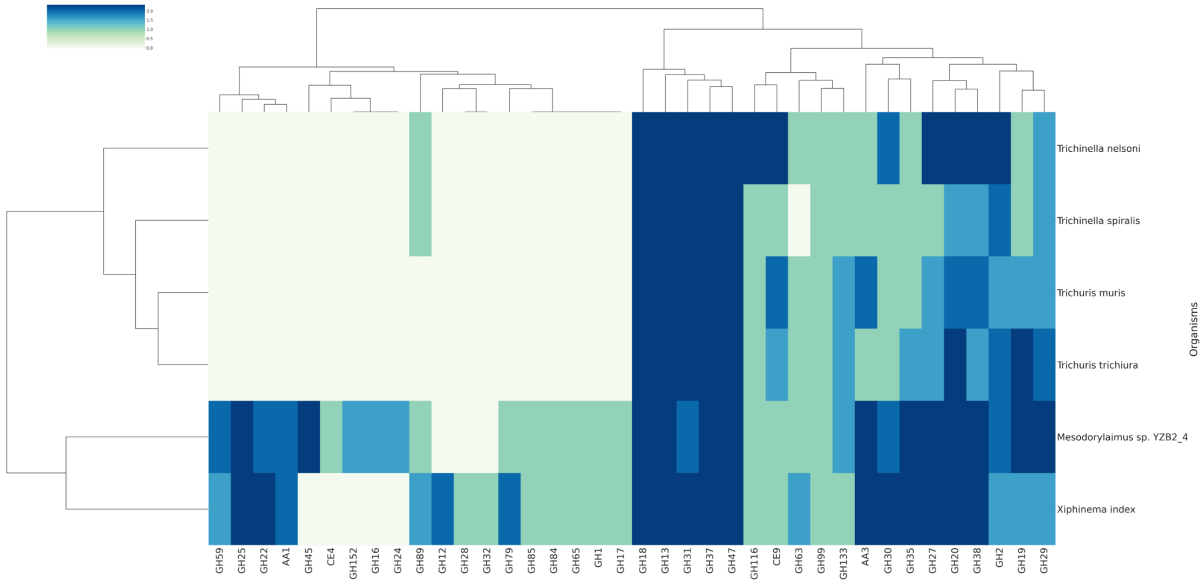


**Figure S4: Clustering of Dorylaimia species according to their degrading CAZome composition.**

On the x-axis the different Glycoside Hydrolase (GH), Auxiliary Activities (AA) and Carbohydrate Esterase (CE) modules are represented (no Polysaccharide Lyase (PL) module present in these species). On the y-axis, the six species: *Trichinella nelsoni*, *Trichinella spiralis*, *Trichuris muris*, *Trichuris trichiura*, *Mesodorylaimus sp*. and *Xiphinema index* are clustered according to the abundance of the CAZyme modules found.

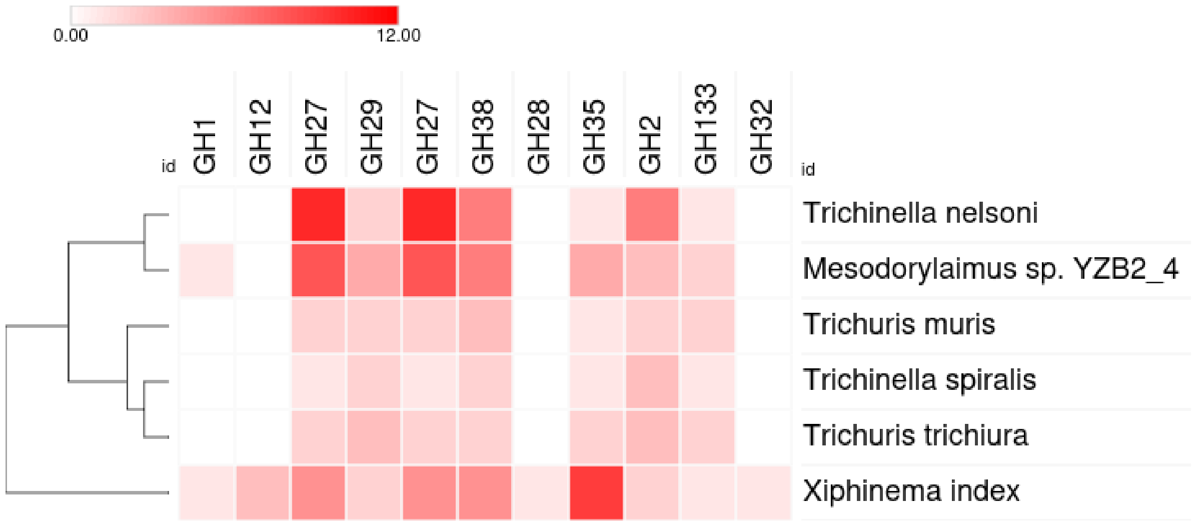


**Figure S5. Clustering of Dorylaimia species according to their abundance in CAZymes with possible degrading activities on plant sugars as substrates.**

The list of CAZymes with such activities was obtained from Figure 1 in supplementary ref[8].

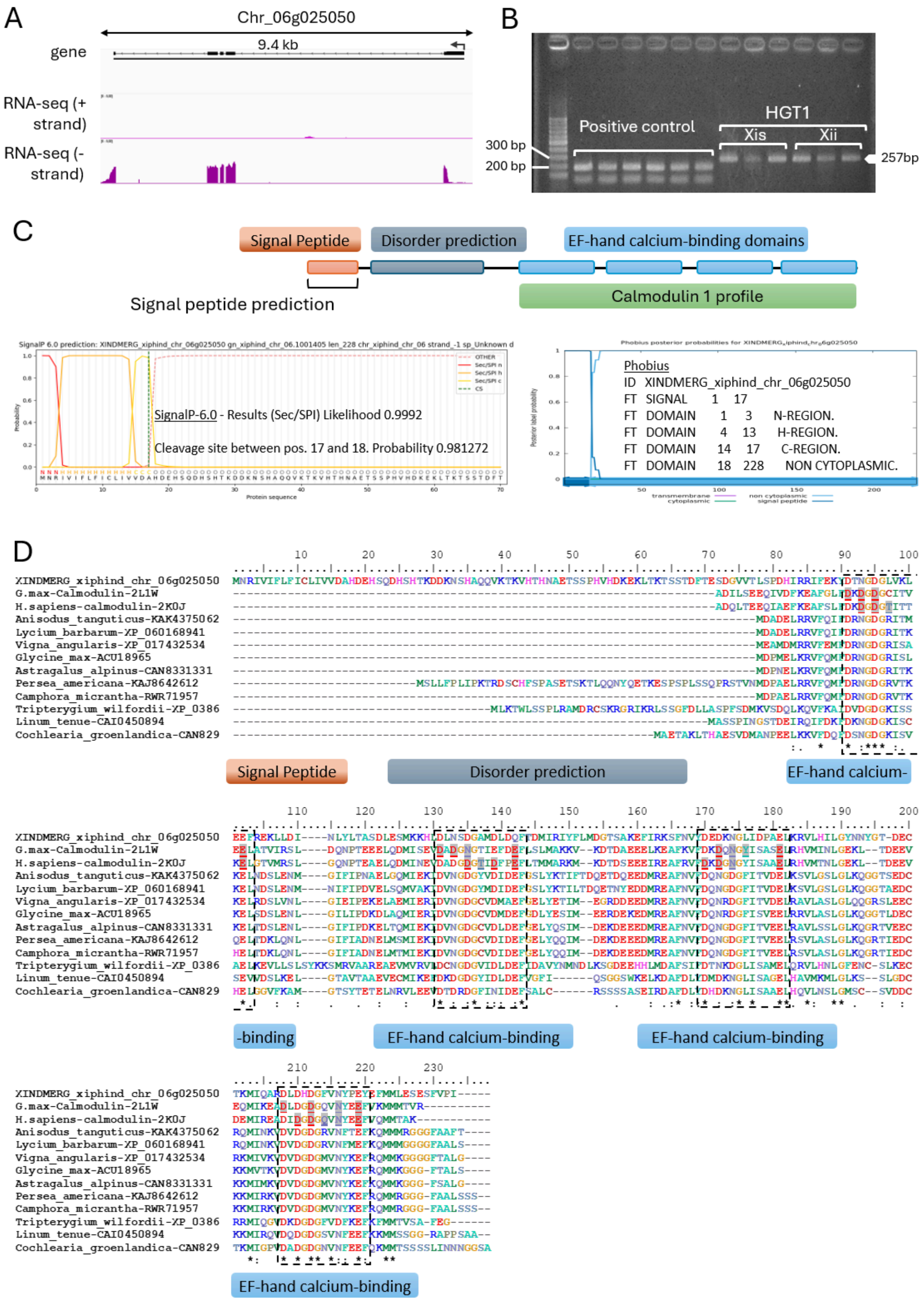


**Figure S6: Characterization of the candidate calmodulin plant-derived HGT (chr_06g_025050).** (A) Structural annotation showing the intron/exon structure and the UTR. The RNA-seq tracks show the gene expression corresponding to each strand (+/-), scale [0-5] CPM. (B) Gel showing the amplification of the marker HGT1, designed in the first intron, from genomic DNA of 2 different populations of *X. index*: samos (Xis) and Iran (Xii). (C) Functional annotation based on InterProScan showing a N-terminal signal peptide, a disordered region and the classical 4 EF-hand calcium binding domains of calmodulin proteins. SignalP 6.0 and Phobius predictions detecting the presence of a signal peptide, with a strong probability, between amino acids 1 to 17. (D) Alignment of chr_06g_025050 with the 10 best hits from BlastP analysis against nr and two crystallized calmodulins from *Glycine max* (L1W) and *Homo sapiens* (2K0J). The alignment shows the domain position, the binding pockets (dotted frames) including amino acids involved in calcium binding (shaded and underlined).

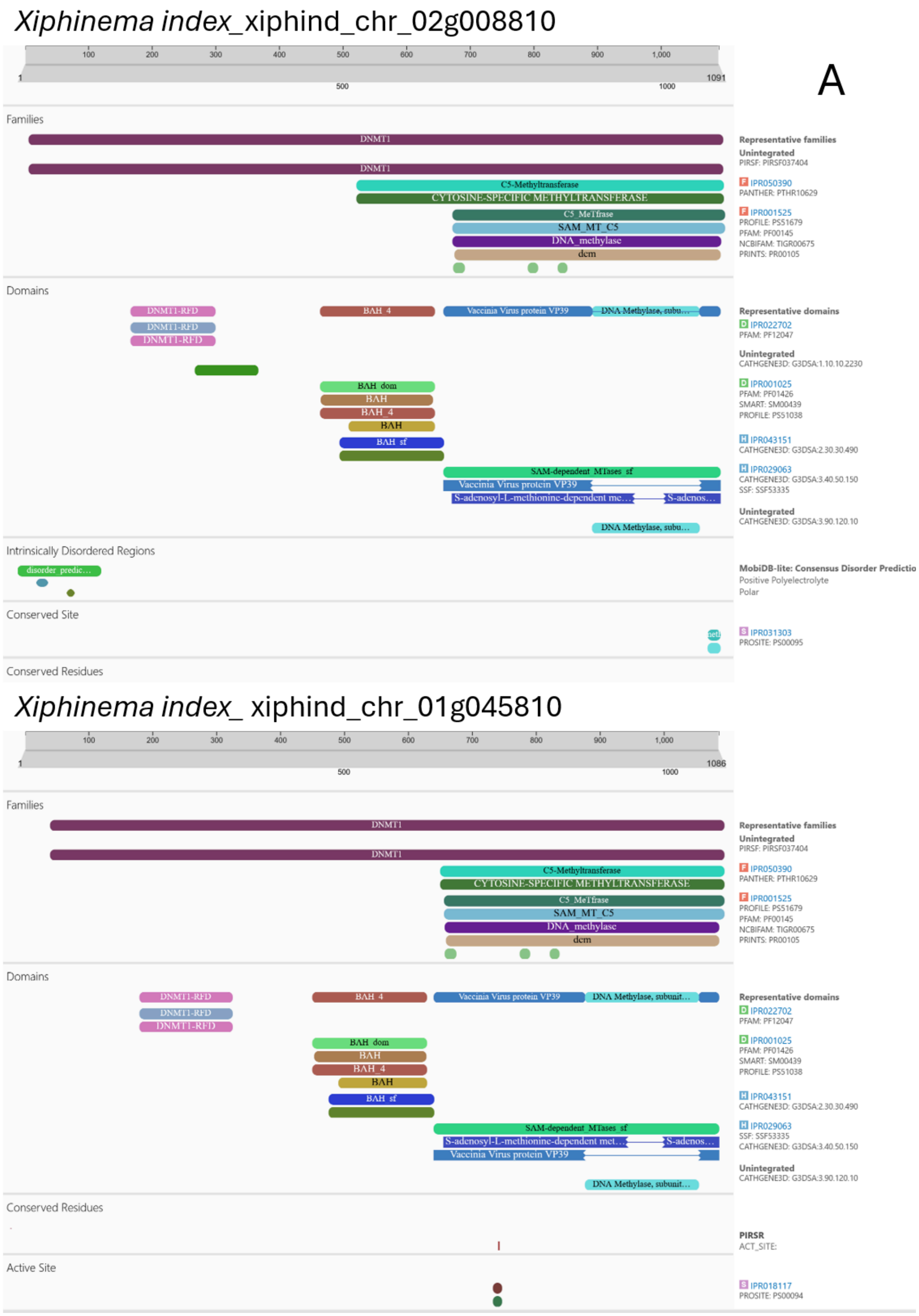
Xiphinema index_xiphind_chr_02g008810
A
Families
DNMT1
DNMT1
C5-Methyltransferase
CYTOSINE-SPECIFIC METHYLTRANSFERASE
C5_MeTfrase
SAM_MT_C5
DNA_methylase
dcm
Representative families
Unintegrated
PIRSF: PIRSF037404
IPR050390
PANTHER: PTHR10629
IPR001525
PROFILE: PS51679
PFAM: PF00145
NCBIFAM: TIGR00675
PRINTS: PR00105
Domains
DNMT1-RFD
BAH_4
Vaccinia Virus protein VP39
BAH_dom
BAH
BAH_sf
SAM-dependent_MTases_sf
S-adenosyl-L-methionine-dependent me…
Representative domains
IPR022702
PFAM: PF12047
Unintegrated
CATHGENE3D: G3DSA:1.10.10.2230
IPR001025
PFAM: PF01426
SMART: SM00439
PROFILE: PS51038
IPR043151
CATHGENE3D: G3DSA:2.30.30.490
IPR029063
CATHGENE3D: G3DSA:3.40.50.150
SSF: SSF53335
Unintegrated
CATHGENE3D: G3DSA:3.90.120.10
Intrinsically Disordered Regions
disorder_predic…
MobiDB-lite: Consensus Disorder Predictio
Positive Polyelectrolyte
Polar
Conserved Site
IPR031303
PROSITE: PS00095
Conserved Residues
Xiphinema index_ xiphind_chr_01g045810
Families
Domains
Conserved Residues
PIRSR
ACT_SITE:
Active Site
IPR018117
PROSITE: PS00094

### *Xiphinema index*_ xiphind_chr_03g023520

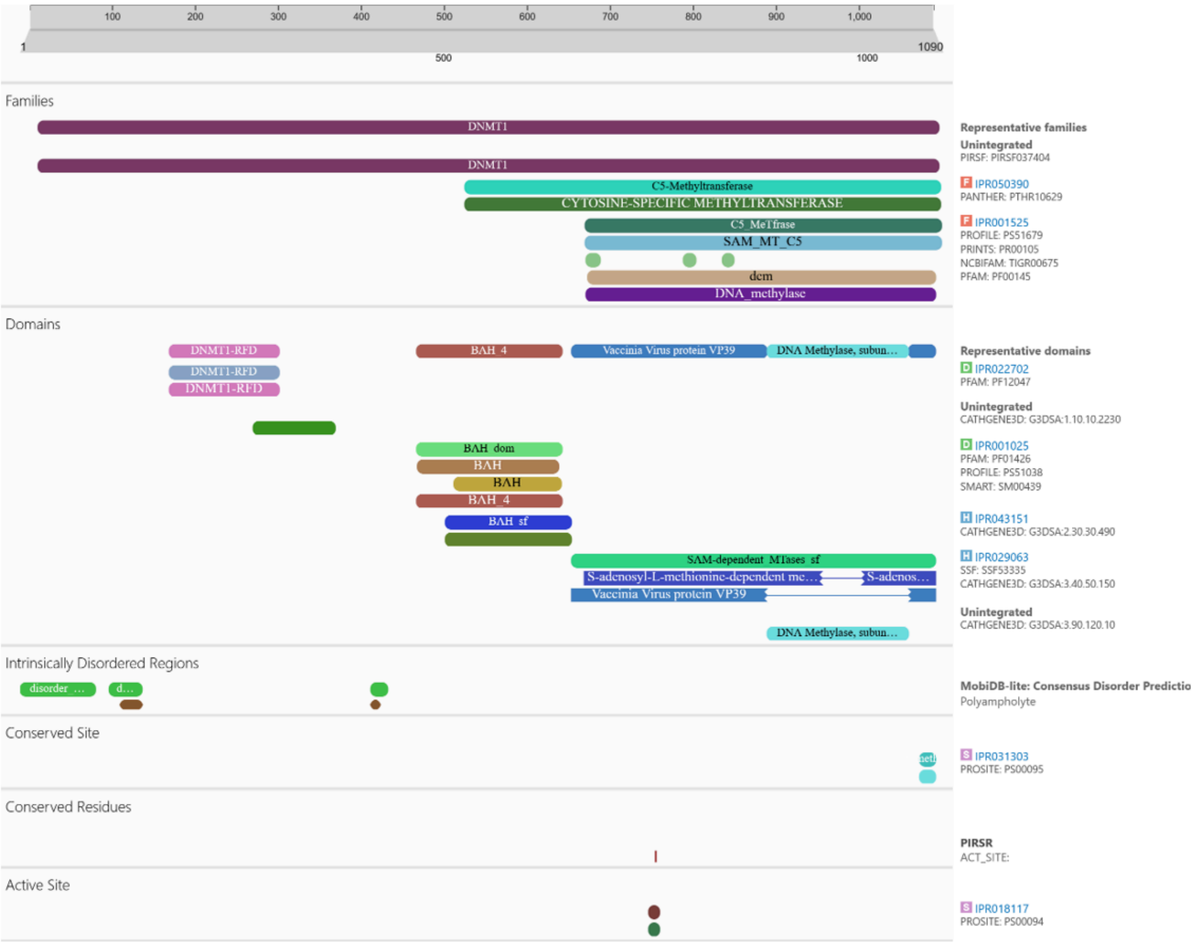


### *Romanomermis culicivorax* (nRc.2.0.1.t46146-RA)

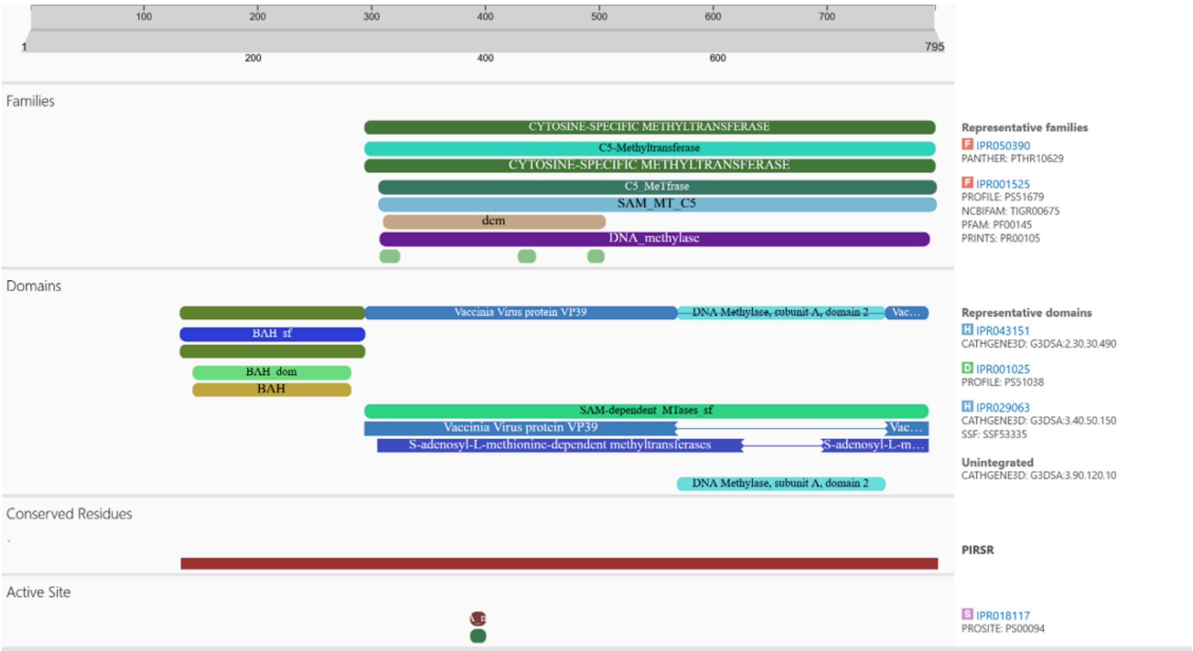

*Plectus sambesii* -PSAMB.scaffold1111size35782.g11016.t1

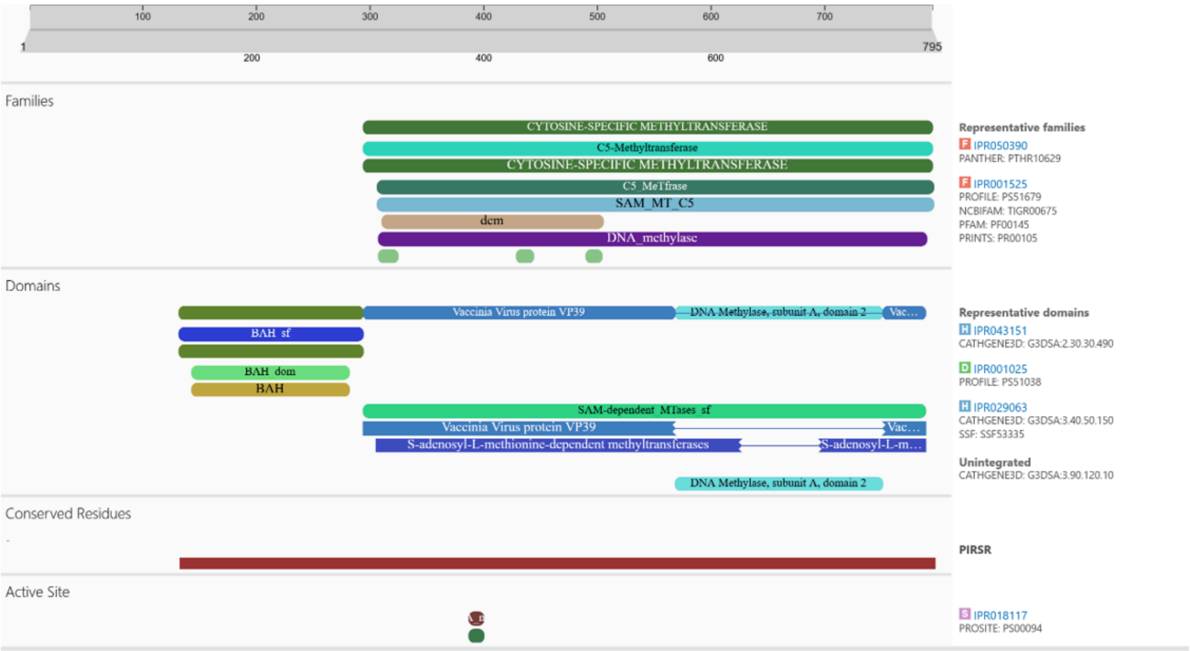


*Trichinella spiralis* putative dnmt1 -EFV58204.1

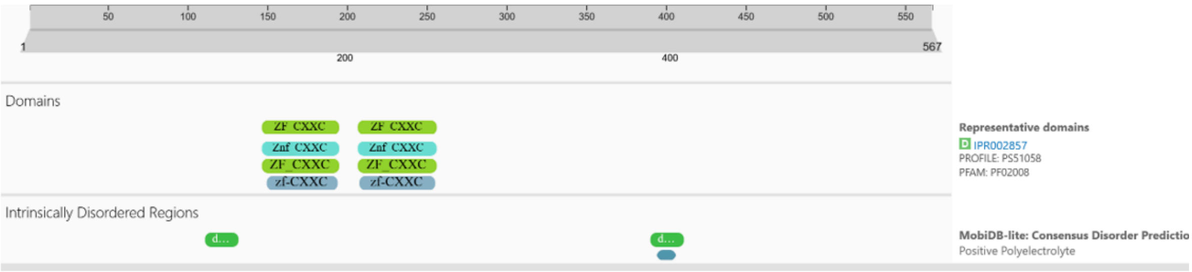


*Bursaphelenchus xylophilus*- putative dnmt1 BUX.s00333.8

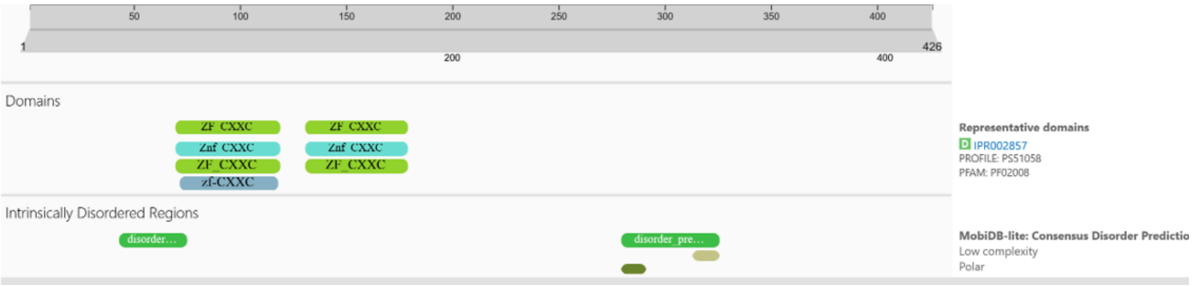

#### *Mesodorylaimus*- QYR22_01172400.t1

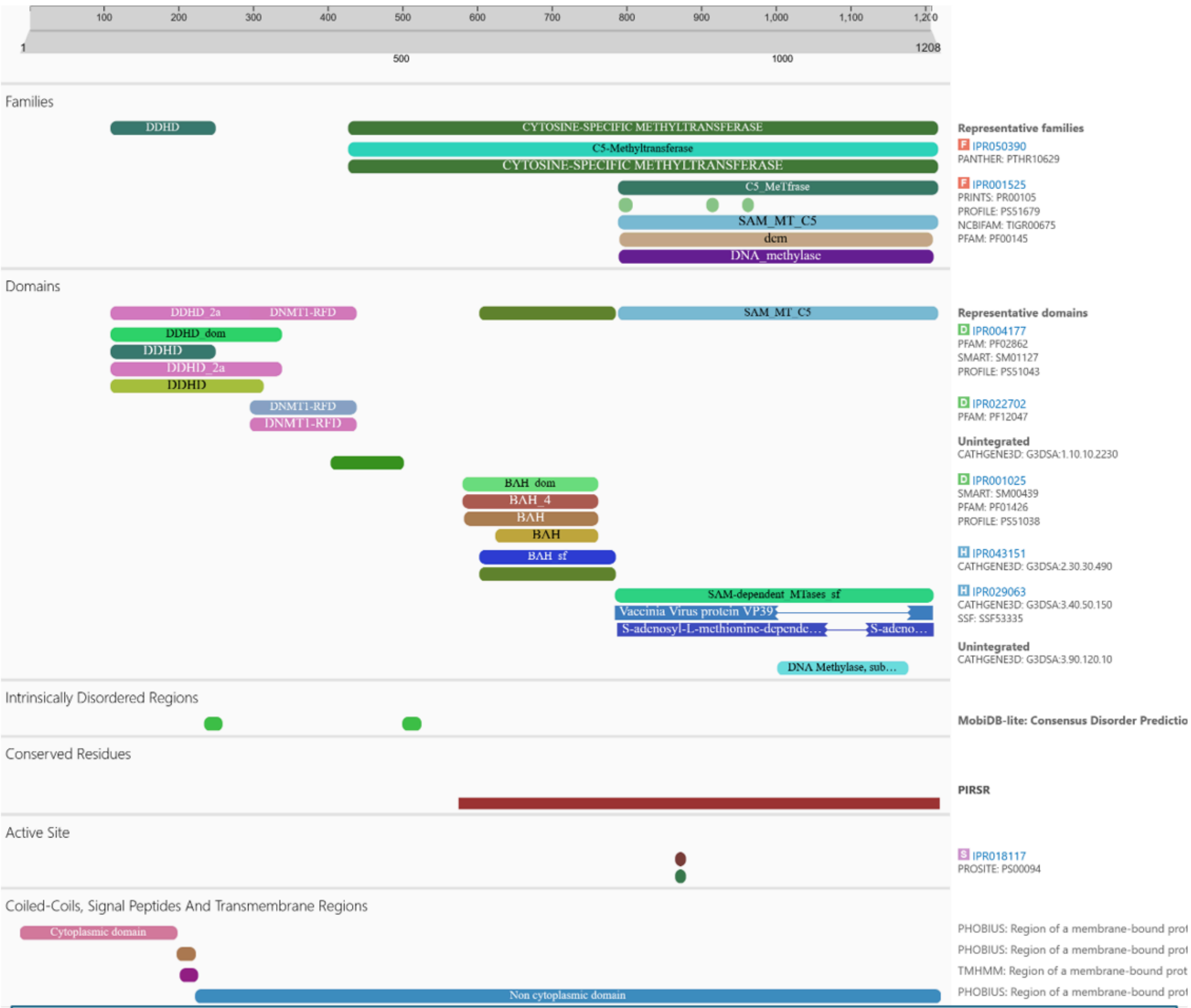


| DNMT1 candidates | DMAP | DNMT1-RFD | Znf-CXXC | BAH | DNA methylase |
|---|---|---|---|---|---|
| *Arabidopsis thaliana-P34881* | | | | | |
| *Homo sapiens-P26358* | | | | | |
| *Gallus gallus-Q92072* | | | | | |
| *Xenopus laevis-P79922* | | | | | |
| *Bombyx mori-Q5W7N6* | | | | | |
| *Xiphinema index-xiphind_chr_01g045810* | | | | | |
| *Xiphinema index-xiphind_chr_03g023520* | | | | | |
| *Xiphinema index-xiphind_chr_02g008810* | | | | | |
| *Mesodorylaimus spp.-QYR22_01172400.t1* | | | | | |
| *Plectus sambesii -PSAMB.scaffold1111size35782.g11016.t1* | | | | | |
| *Romanomermis culicivorax nRc.2.0.1.t46146-RA* | | | | | |
| *Trichinella spiralis -EFV58204.1* | | | | | |
| *Bursaphelenchus xilophilus-BUX.s00333.8* | | | | | |

**DNMT1-RFD domain multiple sequence alignment by MUSCLE (3.8)**

B

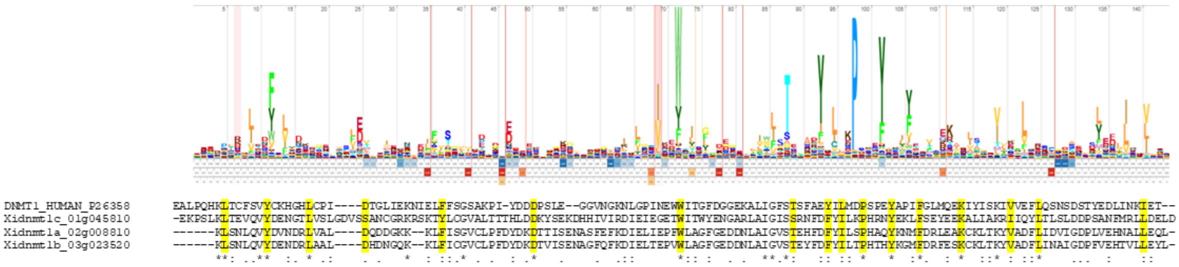

**Figure S7: Functional annotation of the putative dnmt1 homologs in *Xiphinema index* and other nematodes.**

A. comparison of Interproscan domain annotation between the three dnmt1 candidates in *Xiphinema index*, the ortholog in the other *Dorylaimida* species *(Mesodorylaimus sp.)* and the putative nematode dnmt1 described in the literature (*Plectus sambesii, Romanomermis culicivorax, Trichinella spiralis* and *Bursaphelenchus xylophilus)*[9–11]. The table summarizes the domain organization of these putative nematode dnmt1 in a larger context, including a plant (*A. thaliana*), three vertebrates (*H. sapiens, G. gallus, X. laevis*) and an arthropod (*B. mori*) canonical dnmt1 domain organization. B. Sequence alignment of the discriminating DNMT1-RFD domain in human and *X. index* according to the Pfam (PF12047) HMM profile of the domain. Highly conserved amino acids are highlighted.

A

| RFD - DNMT1 *H. sapiens* VS: | sequence alignment score | all pairs | | pruned atom pairs | |
|---|---|---|---|---|---|
| | | pairs | angstroms | pairs | angstroms |
| RFD - dnmt1a *X. index* | 223.2 | 124 | 2.678 | 96 | 0.812 |
| RFD - dnmt1b *X. index* | 226.6 | 124 | 2.263 | 98 | 0.850 |
| RFD - dnmt1c *X. index* | 230.1 | 130 | 2.379 | 104 | 0.866 |

B

| ADD-ATRX *H. sapiens* VS: | sequence alignment score | all pairs | | pruned atom pairs | |
|---|---|---|---|---|---|
| | | pairs | angstroms | pairs | angstroms |
| ADD-atrx *X. index* | 372 | 136 | 5.916 | 102 | 0.890 |

C

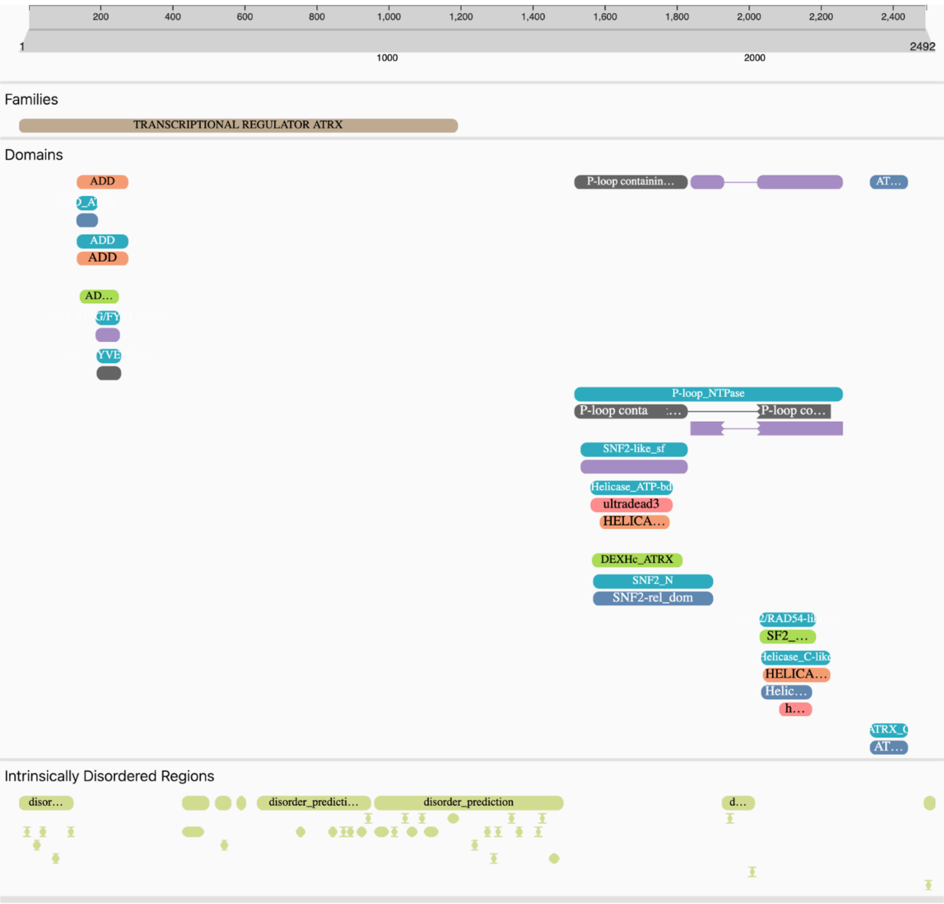

D

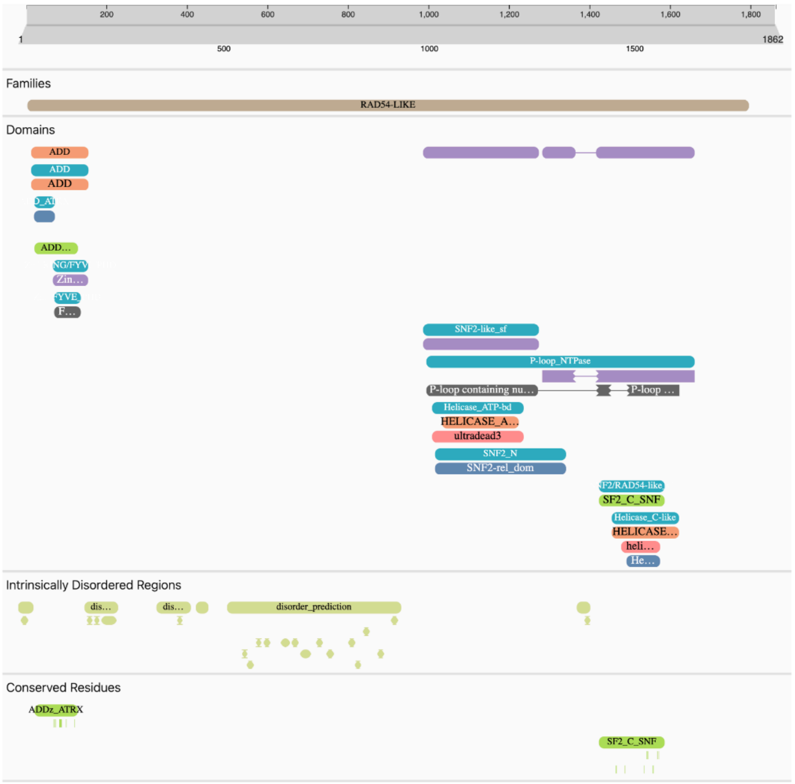


E

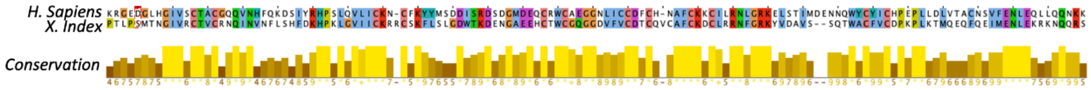


F

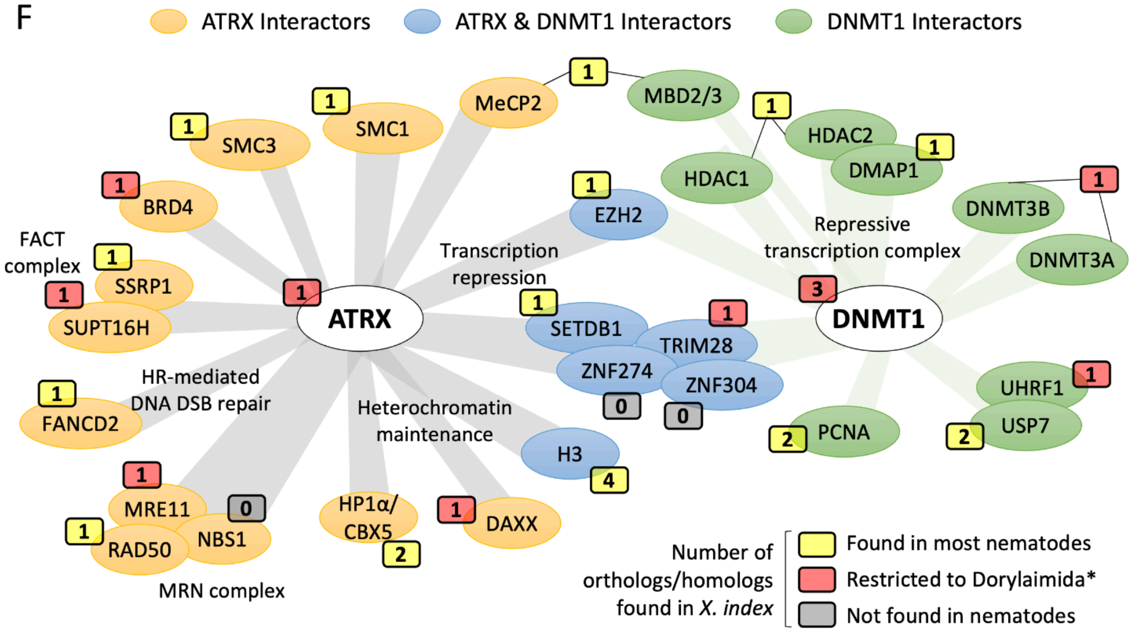


**Figure S8: In-depth functional annotation and putative interactome of DNMT1 and ATRX orthologs in *Xiphinema index***

**A.** Matrix related to superimposes RFD protein structures from the human DNMT1 and *X.index dnmt1a,b and c, respectively.* **B.** Matrix related to superimposes ADD protein structures from the human ATRX and *X.index atrx, respectively.* **C.** Human ATRX Interproscan screenshoot. **D.** *X. index* ATRX Interproscan screenshot. **E.** Amino acid sequence alignment highliting the ADD domain in

Human and *X. index* ATRX. **F.** Main DNMT1 and ATRX interactors. Rectangles indicate the ortholog/homolog numbers found in the predicted proteome of *X. index* and colours indicate if orthologs are detected in other nematode taxa. H3: Histone 3, DAXX: Death domain-associated protein 6, CBX5 (HP1α): Chromobox protein homolog 5, EZH2: Enhancer of Zeste homolog 2, PCNA: proliferating cell nuclear antigen, NBS1: Nibrin, MRE11:Double-strand break repair protein MRE11; MeCP2: Methyl-CpG-binding protein 2;SMC1: Structural maintenance of chromosomes protein 1; SMC3: Structural maintenance of chromosomes protein 3; BRD4: Bromodomain-containing protein 4; SSRP1: FACT complex subunit SSRP1; SUPT16H:FACT complex subunit SPT16; FANCD2:Fanconi anemia group D2 protein; RAD50: DNA repair protein RAD50; SETDB1: Histone-lysine N-methyltransferase SETDB1; TRIM 28: Transcription intermediary factor 1-beta; ZNF274: Neurotrophin receptor-interacting factor homolog; ZNF304: Zinc finger protein 304; USP7: Ubiquitin carboxyl-terminal hydrolase 7; UHRF1: E3 ubiquitin-protein ligase UHRF1; DNMT3A/B: DNA (cytosine-5)-methyltransferase 3A/B; DMAP1: DNA methyltransferase 1-associated protein 1; HDAC1 & 2:Histone deacetylase 1 & 2; MBD2 & 3: Methyl-CpG-binding domain protein 2 & 3; PDS5B:Sister chromatid cohesion protein PDS5 homolog B; ATRX: Transcriptional regulator ATRX; DNMT1: DNA (cytosine-5)-methyltransferase 1. *including or not nematodes from basal clades (1 & 2).

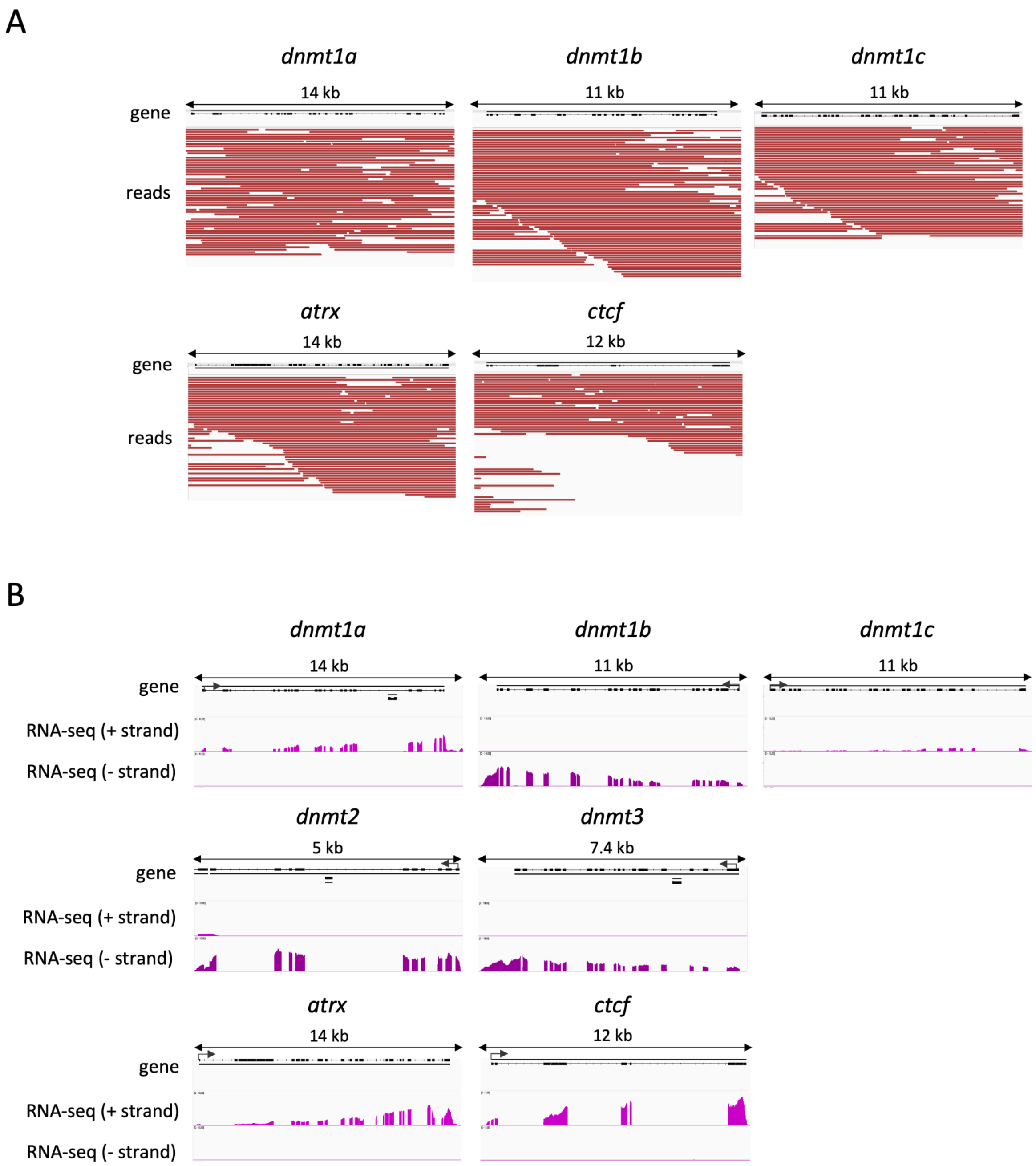


**Figure S9: Visualization of the PacBIO long read coverages and transcriptomic profiles at the loci of dnmts, atrx and ctcf in *X. index.*** A. PacBIO long read coverages at the loci of dnmts1a, dnmt1b, dnmt1c, atrx and ctcf in *X. index.* B.Transcriptomic profiles at the loci of dnmts1a, dnmt1b, dnmt1c, dnmt2, dnmt3, atrx and ctcf in *X. index.* The RNA-seq tracks show the gene expression corresponding to each strand (+/-), scale [0,10] CPM.

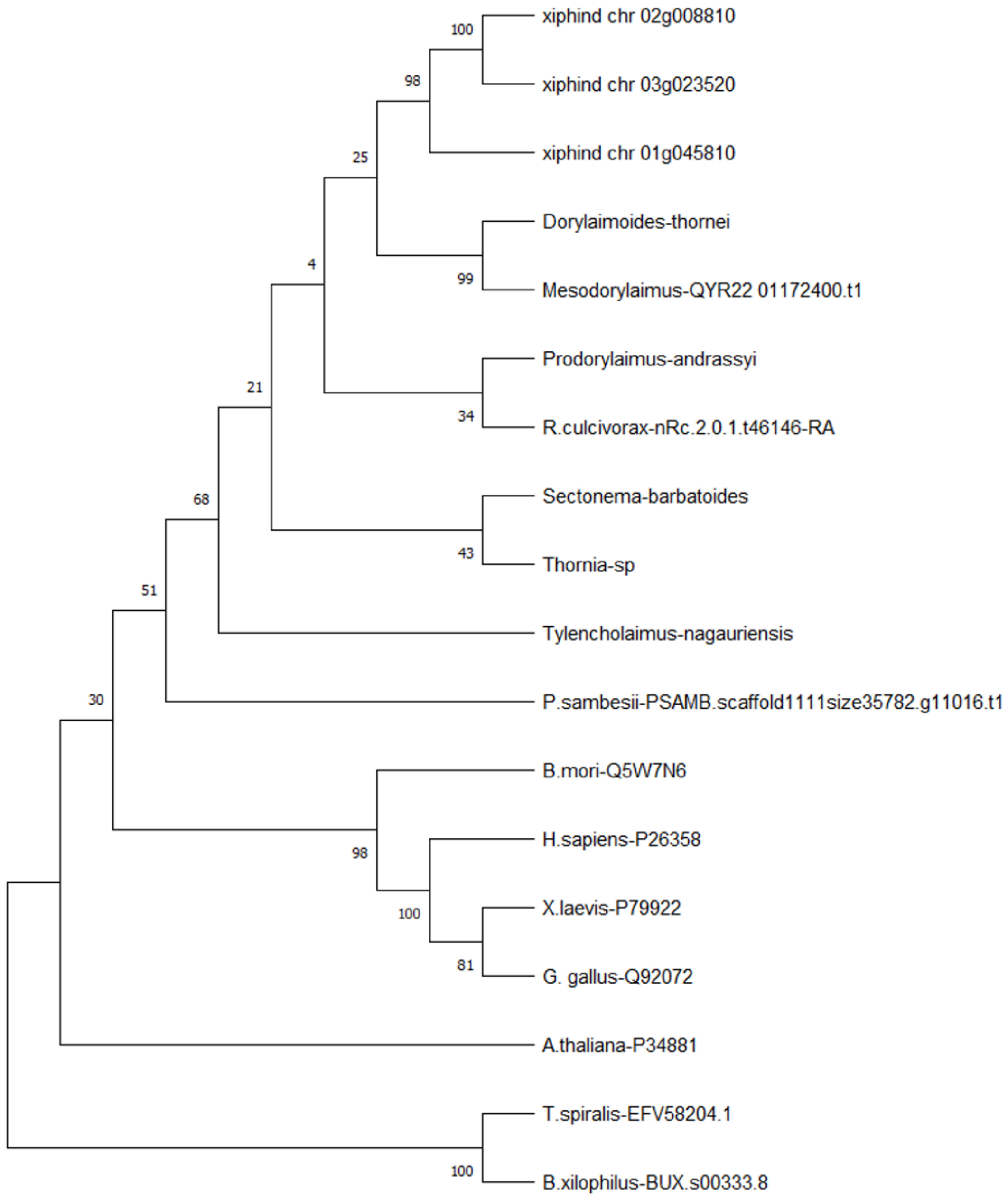


**Figure S10: Phylogenetic tree based on DNMT1 sequences detected in this study and reference sequences.**

The tree was inferred by using the Maximum Likelihood method and JTT matrix-based model. The tree with the highest log likelihood (-38669.76) is shown. The percentage of trees (bootstrap =500) in which the associated taxa clustered together is shown next to the branches. Initial tree(s) for the heuristic search were obtained automatically by applying Neighbor-Joining and BioNJ algorithms to a matrix of pairwise distances estimated using the JTT model, and then selecting the topology with superior log likelihood value. This analysis involved 18 amino acid sequences. There were a total of 1876 positions in the final alignment. Evolutionary analyses were conducted in MEGA X. Sequences include 5 representative sequences (from *Arabidopsis thaliana, Gallus gallus, Xenope laevis, Homo sapiens* and *Bombyx mori,* the 4 sequences extracted from previous studies (from *Trichinella spiralis, Bursaphelenchus xylophilus, Plectus sambesii* and *Romanomermis culcivorax*) and 9 sequences detected in this study (from *X. index, Tylencholaimus nagauriensis, Thornia sp, Sectonema barbatoides, Prodorylaimus andrassyi, Dorylaimoides thornei* and *Mesodorylaimus*).

A

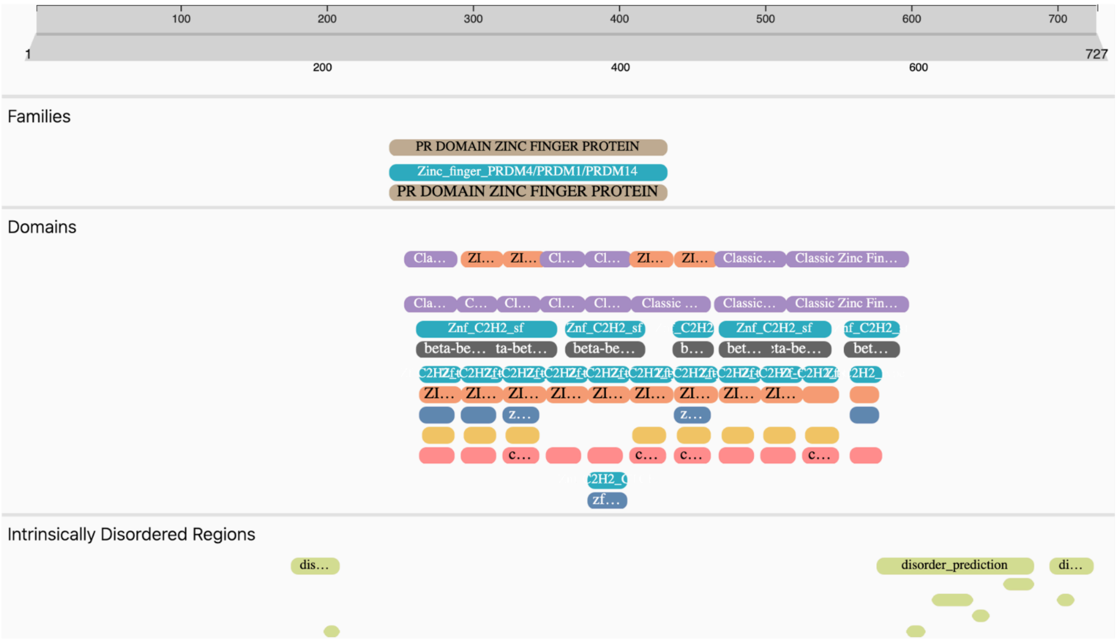

100
200
300
400
500
600
700
1
727
200
400
600
Families
PR DOMAIN ZINC FINGER PROTEIN
Zinc_finger_PRDM4/PRDM1/PRDM14
PR DOMAIN ZINC FINGER PROTEIN
Domains
Classic Zinc Fin…
Znf_C2H2_sf
disorder_prediction
Intrinsically Disordered Regions


B

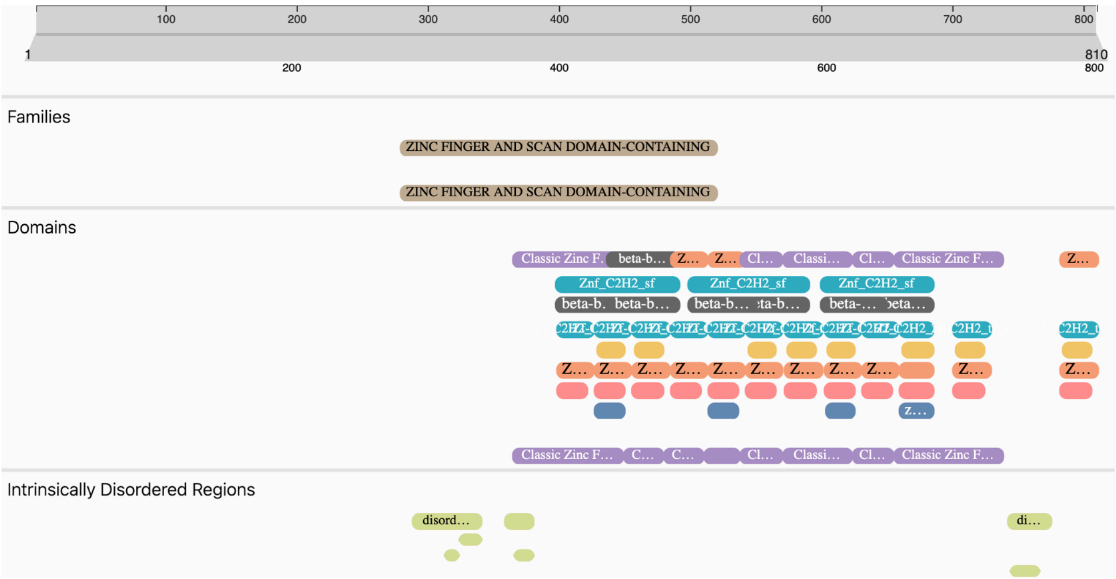

100
200
300
400
500
600
700
800
1
810
200
400
600
800
Families
ZINC FINGER AND SCAN DOMAIN-CONTAINING
ZINC FINGER AND SCAN DOMAIN-CONTAINING
Domains
Classic Zinc F…
Znf_C2H2_sf
Intrinsically Disordered Regions
disord…

C

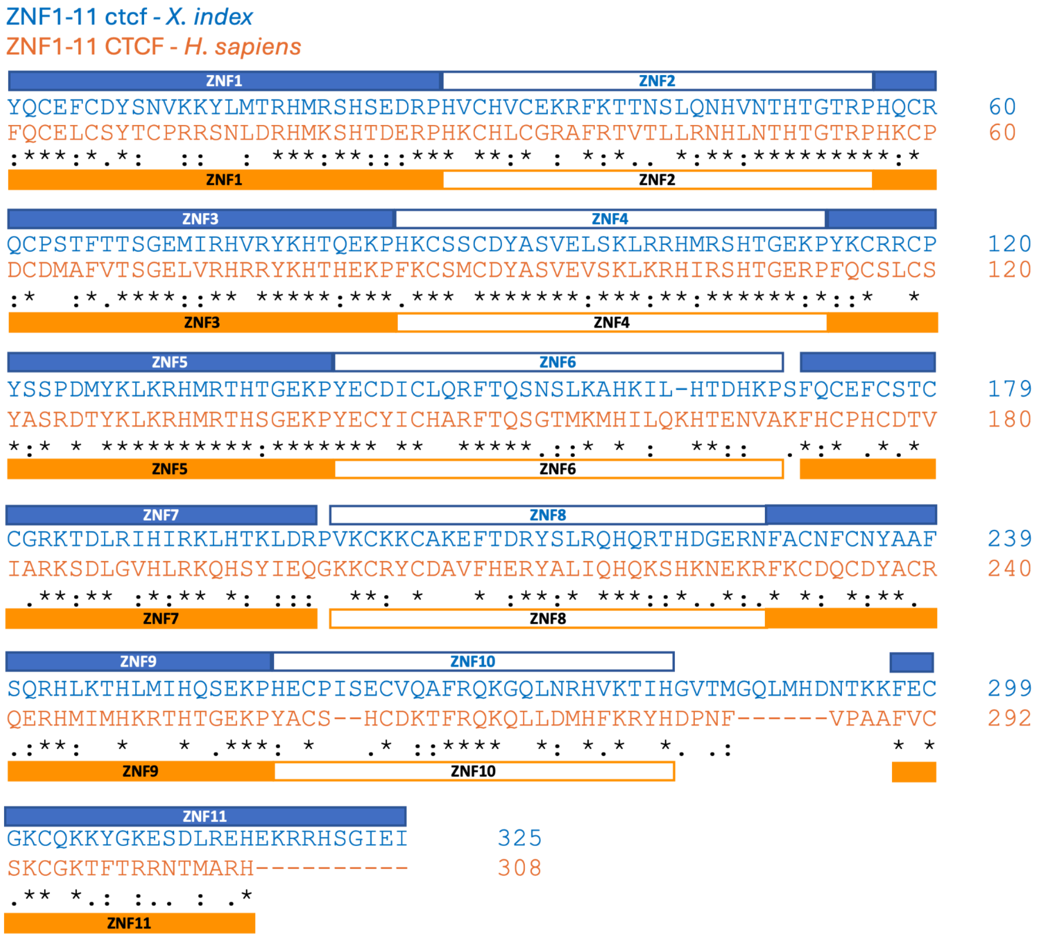


**Figure S11: Functional annotation comparison between the human and *Xiphinema index* CTCF.** A. InterProScan screenshot of *H. sapiens* CTCF. B. InterProScan screenshot of *X. index* CTCF. C. Amino acid sequence alignment of the ZNF1-11 from *X. index* (in blue) and *H. sapiens* (in orange).

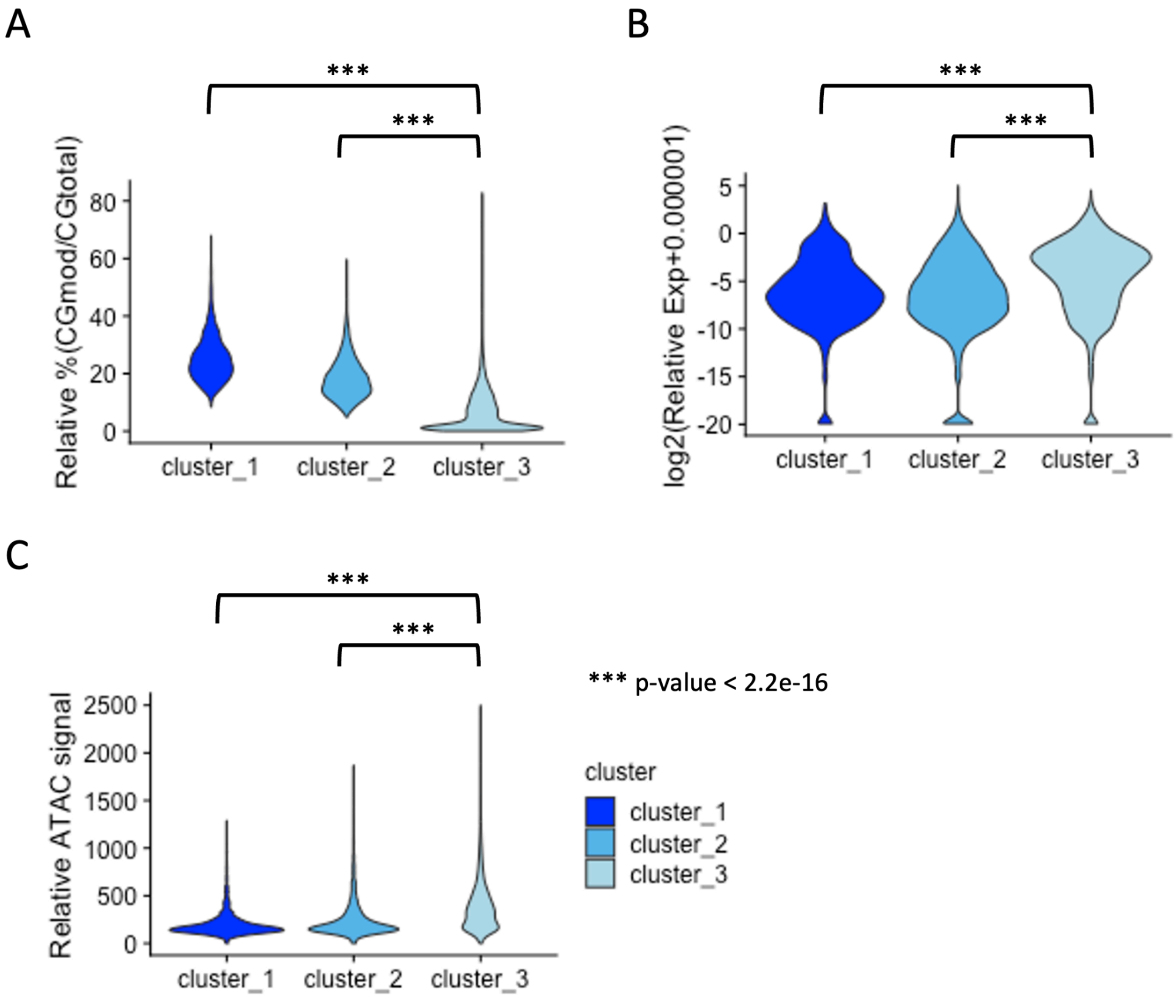


**Figure S12: Violin plots of k-means-based clusters 1 to 3 (-300/+200 bp surrounding TSS) showing the relative % of methylated CG over total CG.**

A. The relative expression of genes. B. The chromatin accessibility based on relative ATAC-seq signal. C. ***p-value<$2.2e^{-16}$, Wilcoxon tests. n=2887 (cluster_1), n=8372 (cluster_2) and n=9524 (cluster_3)

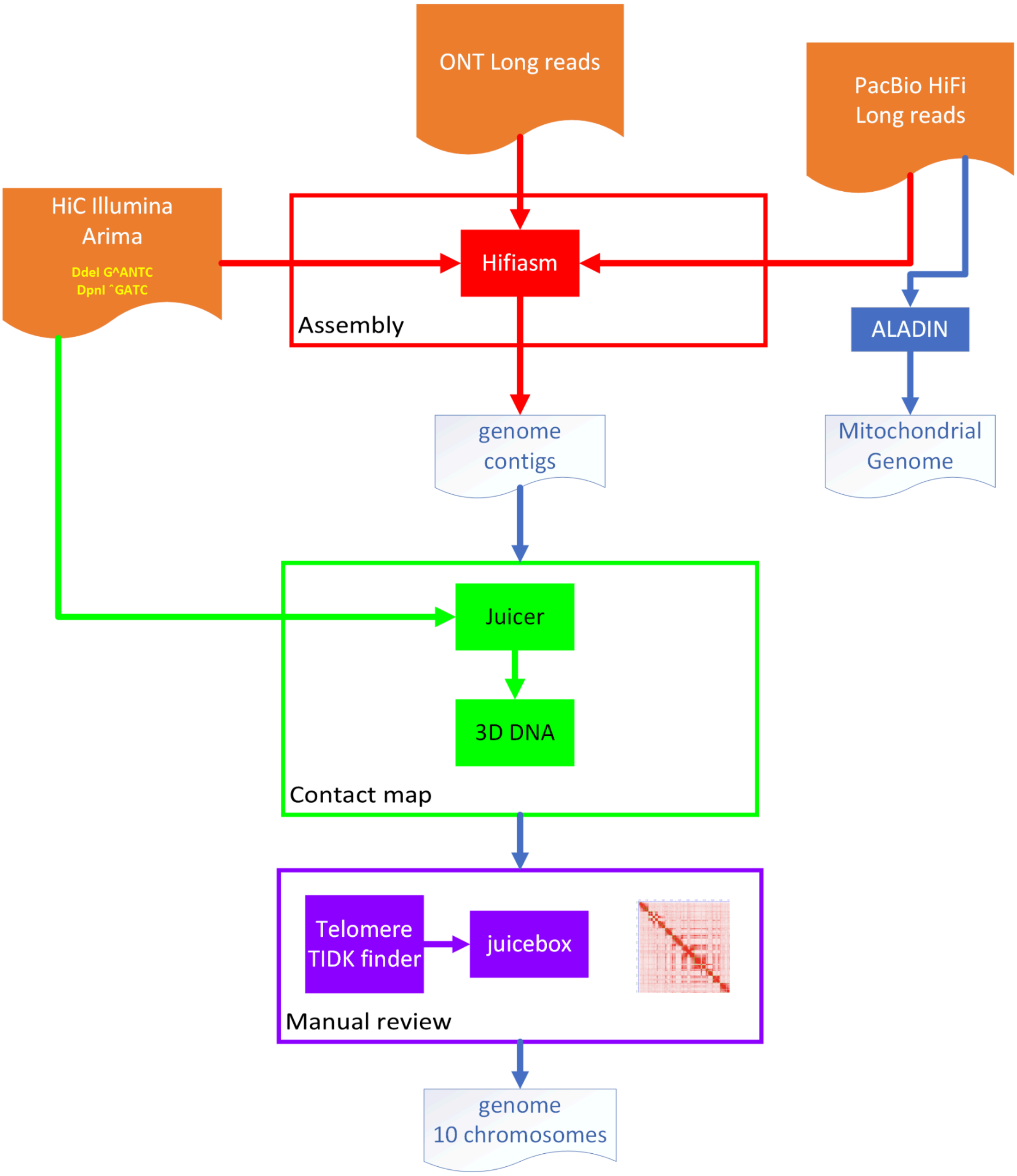


**Figure S13: Flowchart illustrating the production of *Xiphinema index* assembly.**

### Cited references


1. Baril, T., Galbraith, J. & Hayward, A. Earl Grey: A Fully Automated User-Friendly Transposable Element Annotation and Analysis Pipeline. *Mol. Biol. Evol.* **41**, msae068 (2024).
2. Manni, M., Berkeley, M. R., Seppey, M., Simão, F. A. & Zdobnov, E. M. BUSCO update: novel and streamlined workflows along with broader and deeper phylogenetic coverage for scoring of eukaryotic, prokaryotic, and viral genomes. *Mol. Biol. Evol.* https://doi.org/10.1093/molbev/msab199 (2021) doi:10.1093/molbev/msab199.
3. Danchin, E. G. J. *et al.* The Transcriptomes of Xiphinema index and Longidorus elongatus Suggest Independent Acquisition of Some Plant Parasitism Genes by Horizontal Gene Transfer in Early-Branching Nematodes. *Genes* **8**, 287 (2017).
4. Ranallo-Benavidez, T. R., Jaron, K. S. & Schatz, M. C. GenomeScope 2.0 and Smudgeplot for reference-free profiling of polyploid genomes. *Nat. Commun.* **11**, 1–10 (2020).
5. Mapleson, D., Garcia Accinelli, G., Kettleborough, G., Wright, J. & Clavijo, B. J. KAT: a K-mer analysis toolkit to quality control NGS datasets and genome assemblies. *Bioinformatics* **33**, 574–576 (2017).
6. Challis, R., Richards, E., Rajan, J., Cochrane, G. & Blaxter, M. BlobToolKit – Interactive Quality Assessment of Genome Assemblies. *G3 GenesGenomesGenetics* **10**, 1361–1374 (2020).
7. Yoshida, Y. *et al.* Comparative genomics of the tardigrades Hypsibius dujardini and Ramazzottius varieornatus. *PLOS Biol.* **15**, e2002266 (2017).
8. Li, J. *et al.* Comparative Genomics and Transcriptomics Analyses Reveal Divergent Plant Biomass-Degrading Strategies in Fungi. *J. Fungi* **9**, 860 (2023).
9. Engelhardt, J., Scheer, O., Stadler, P. F. & Prohaska, S. J. Evolution of DNA Methylation Across Ecdysozoa. *J. Mol. Evol.* **90**, 56–72 (2022).
10. Gao, F. *et al.* Differential DNA methylation in discrete developmental stages of the parasitic nematode Trichinella spiralis. *Genome Biol.* **13**, R100 (2012).
11. Liu, Z., Li, Y. & Zhang, X. DNA methylation on C5-Cytosine and N6-Adenine in the Bursaphelenchus xylophilus genome. *BMC Genomics* **24**, 671 (2023).
